# Backbone extension acyl rearrangements enable cellular synthesis of proteins with internal β^2^-peptide linkages

**DOI:** 10.1101/2023.10.03.560714

**Authors:** Leah T. Roe, Carly K. Schissel, Taylor L. Dover, Bhavana Shah, Noah X. Hamlish, Shuai Zheng, Diondra A. Dilworth, Nicole Wong, Zhongqi Zhang, Abhishek Chatterjee, Matthew B. Francis, Scott J. Miller, Alanna Schepartz

**Affiliations:** Department of Chemistry, University of California, Berkeley, CA 94720, USA; Department of Chemistry, Yale University, New Haven, CT 06520; Process Development, Attribute Sciences, Amgen Inc., Thousand Oaks, CA 91320; Molecular and Cell Biology, University of California, Berkeley, CA 94720, USA; Department of Chemistry, Boston College, Chestnut Hill, MA; California Institute for Quantitative Biosciences, University of California, Berkeley, CA 94720, USA; Chan Zuckerberg Biohub, San Francisco, CA 94158, USA; Innovation Investigator, ARC Institute, Palo Alto, CA 94304, USA

## Abstract

Proteins and polypeptides containing extended backbone monomers embody highly desirable structures and functions, but they cannot yet be biosynthesized in cells. There are two challenges at work. First is the ribosome, whose ability to promote rapid bond-forming reactions to and from anything other than an α-amino acid or α-hydroxy acid is unknown. The second challenge is the absence of orthogonal enzymes that acylate tRNA with extended backbone monomers. Here we describe a general approach to the programmed cellular synthesis of proteins containing extended backbone monomers that circumvents both of these challenges. Rather than relying on direct and uncharacterized reactions of non-α-amino acid monomers within the ribosomal PTC, we develop a proximity-guided intramolecular rearrangement that effectively edits the protein backbone post-translationally. The method relies on the ability of PylRS-like aminoacyl-tRNA synthetase enzymes to accept diverse α-hydroxy acid monomers, including those whose side chains contain masked nucleophiles. Introduction of such an α-hydroxy acid monomer into a protein translated *in vivo*, followed by nucleophile unmasking, sets up a thermodynamically favored and quantitative intramolecular Backbone Extension Acyl Rearrangement (BEAR) reaction that edits the protein backbone to install an extended backbone monomer. In the examples described here, the intramolecular rearrangement converts an α-peptide backbone directly into a β-backbone. As far as we know, this report represents the first example in which a much-desired expanded backbone β-amino acid linkage has been introduced site-selectively into a protein in a cell.

## Main

There is widespread current interest in the cellular biosynthesis of proteins and polypeptides whose backbones differ from those in extant proteins^1^. Even single-atom substitutions, such as the introduction of an ester^2–8^ or thioester^9–11^ in place of an amide, can promote new chemistry^12^, facilitate mechanistic studies^2^, and generate materials with emergent properties^13–15^. Small molecule synthetic chemists well-appreciate the impact of single atom substitutions^16^. The simplest α-peptide backbone modification, beyond a single-atom substitution, is the addition of a single methylene (CH_2_) unit to generate a β^2^- or β^3^-amino acid (**Figure 1A**). β^2^- and β^3^-amino acid linkages are replete in natural products, such as taxol^17^, andrimid^18^, and guineamide A^19^, that possess profound biological activity. Oligomers containing one or more β-amino acid linkages can function as protein-protein interaction inhibitors^20–26^, antimicrobial agents^27–33^, receptor agonists and antagonists^34–48^, catalysts^49,50^, and other useful tools. Certain β-peptide oligomers assemble into structurally defined protein-like materials^51–54^ that fold cooperatively and with enhanced thermal stability^52,53^. Even a small number of judiciously placed β-linkages within a protein add value, such as improved protease resistance^55–57^, enhanced cellular permeability^58–60^, and altered recognition when presented by class I major histocompatibility complex (MHC) receptors^61,62^.

**Figure 1.**
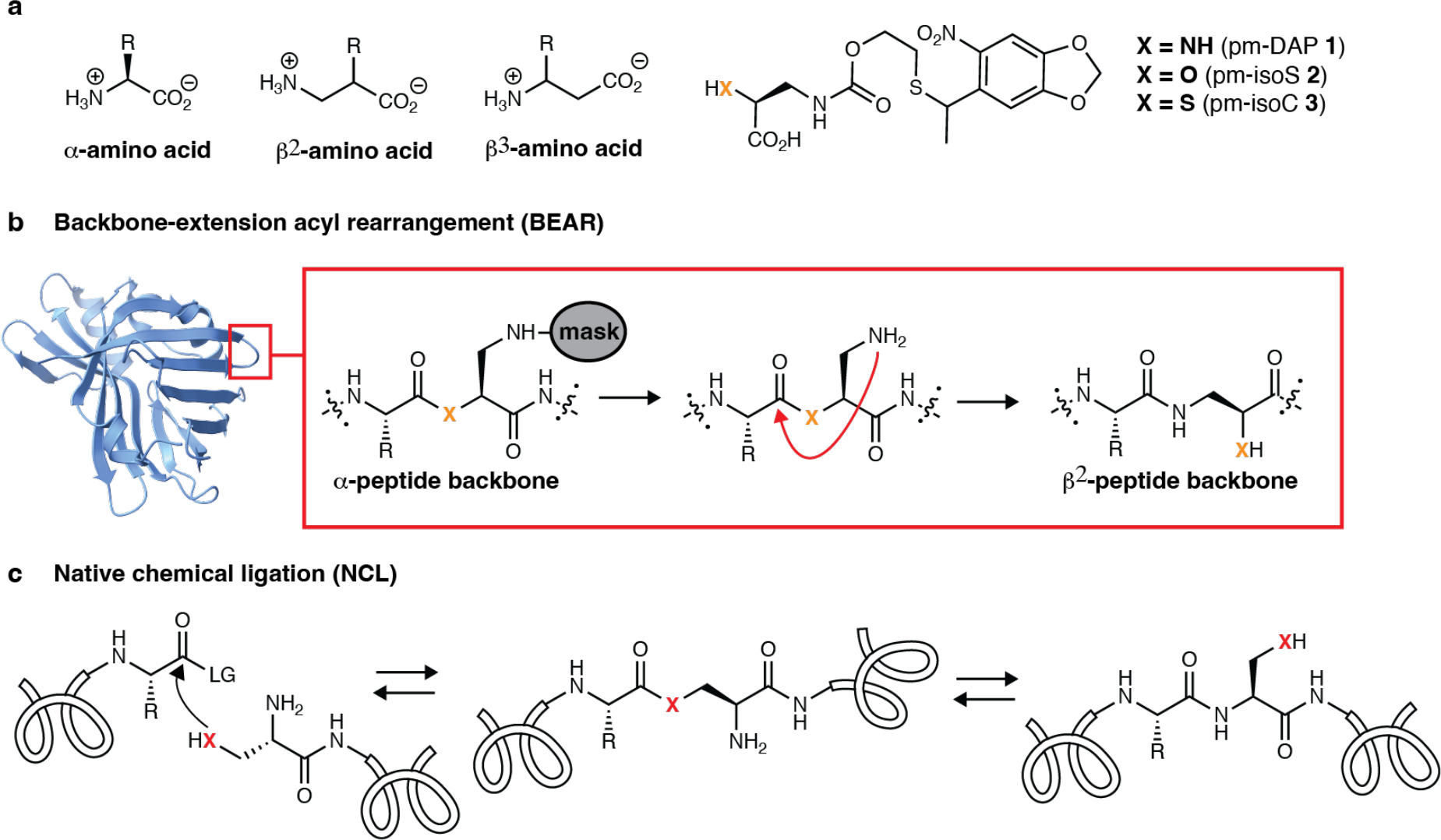
Backbone Extension Acyl Rearrangement (BEAR) reactions create extended backbones in full-length proteins expressed *in vivo*. (A) Structures of α-, β^2^, and β^3^-amino acids as well as photo-masked (pm) analogs of diaminopropionic acid (pm-DAP, **1**), isoS (pm-isoS, **2**), and isoC (pm-isoC, (**±)-3**). (B) A BEAR reaction results from the intramolecular cyclization of an unmasked nucleophile, in this case, a primary amine, on an N-terminal ester or thioester. In the case shown, breakdown of the resultant tetrahedral intermediate generates a β^2^-peptide linkage embedded within the protein chain. PDB ID 7SNS. (C) BEAR reactions are analogous to the second step of the well-known native chemical ligation (NCL) reaction^9,71^.

Despite the gain in function provided by β-linkages, there exists only a single example^63^ in which a β-amino acid has been incorporated into a protein *in vivo*. That effort required an engineered ribosome with a structurally characterized assembly defect^64^ and could not be generalized due to the absence of an orthogonal β-amino acid-selective aminoacyl-tRNA synthetase. *In vitro* translation systems containing wild type ribosomes can introduce certain β^2^- and β^3^-amino acids into polypeptides^65,66^, but this methodology is not easily scalable and relies on stoichiometric RNA acylation reagents that have not been shown to function in cells. Indeed, it has been hypothesized that complex engineering of both an orthogonal aminoacyl-tRNA synthetase and the ribosome will be required to create a programmable system for the cellular synthesis of proteins containing β-amino acid linkages^7,67,68^. Here we describe an approach to the programmed cellular synthesis of β-amino acid-containing proteins that does not require engineering of either a synthetase or the ribosome.

Rather than relying on direct reaction of a β-amino acid monomer within the ribosomal peptidyl-transferase center (PTC)^63^, the strategy reported here relies on a proximity-guided intramolecular Backbone Extension Acyl Rearrangement (BEAR) that effectively edits the protein backbone post-translationally to orthogonally and site-specifically install a β-amino acid residue into full-length proteins *in vivo*. BEAR reactions rely on the fact that α-hydroxy acids are excellent substrates^4,5,8,7^ for pyrrolysyl-tRNA synthetase (PylRS), the aminoacyl-tRNA synthetase that naturally incorporates pyrrolysine into proteins in certain archaea and methanogenic bacteria, and is also orthogonal in *E. coli*. Recent work has shown that α-thiol acids^7^ are also substrates for certain PylRS variants. Furthermore, PylRS variants accept α-amino acids with a variety of side chains^69^, including those carrying a protected nucleophile^70^. We envisioned that if an α-hydroxy or α-thiol substrate for such a PylRS variant carried a side chain bearing a masked β-amine nucleophile, that nucleophile could be unmasked post-translationally to promote a BEAR reaction (**Figure 1B**) that is analogous to the second step of the canonical and now ubiquitous native chemical ligation (NCL) reaction^9,71^ (**Figure 1C**). The result of a BEAR reaction would be an internal β^2^-linkage.

Related acyl rearrangements within short peptides are known. It was reported in 2004 that isocysteine (isoC) supports an NCL-like reaction to install a β^2^-linkage into a chemically synthesized short peptide *in vitro*^72^. More recently, it was shown that backbone rearrangements involving isoserine (isoS) can occur in short peptides prepared during small-scale *in vitro* translation reactions^12^. Inspired by these findings, we report that proteins containing a photo-masked isoS derivative can be unmasked post-translationally, either in cells or after protein purification. Unmasking reveals the β-amine nucleophile and triggers a spontaneous, intramolecular BEAR reaction that directly installs a β^2^-amino acid linkage. This work represents a new, generalizable strategy to install extended backbones into proteins *in vivo*, and relies only on established activities of known orthogonal enzymes and the wild type *E. coli* ribosome. The hetero-oligomers so-generated expand the diversity of protein-like polypeptides that may be synthesized ribosomally.

## Results and Discussion

### Computational studies confirm that BEAR reactions of isoS and isoC are thermodynamically favorable

To establish thermodynamic plausibility, we performed computational studies and model reactions to confirm that intramolecular rearrangements of isoC and isoS linkages within proteins would be favorable under physiological conditions. A previous study has used DFT to examine the energetics and mechanistic pathways of a canonical cysteine-based NCL reaction, but this study did not include the isoC, Ser, or isoS variants considered herein^73^. For our analysis, we used Density Functional Theory (DFT) calculations to evaluate the relative energetics of acyl rearrangement reactions involving Cys, Ser, isoC, and isoS nucleophiles in the context of a simplified substrate (**Figure 2A**). For each proposed species in the reaction, molecular mechanics methods with an OPLS4 force field were first used to generate and evaluate 10,000 conformers for each species. The geometries and electronic energies of all conformers with energies within 5 kcal/mol of the global minimum were then re-optimized using DFT methods (B3LYP-D3/6-31G**). The global minimum conformation for each intermediate was then subjected to single point calculations to determine the vibrational frequencies at the B3LYP-D3/6-31G** level. This analysis allowed estimation of the zero-point energy and internal entropy parameters. Finally, electronic energies were determined with greater accuracy using ωB97M-V/6-311++G(3df,3dp). These final calculations were performed in the gas phase and with a CPCM solvent model to represent aqueous solvation. Results from the solvated calculations were used to generate the final thermodynamic values reported herein (see Extended Data 1 for gas phase values).

**Figure 2.**
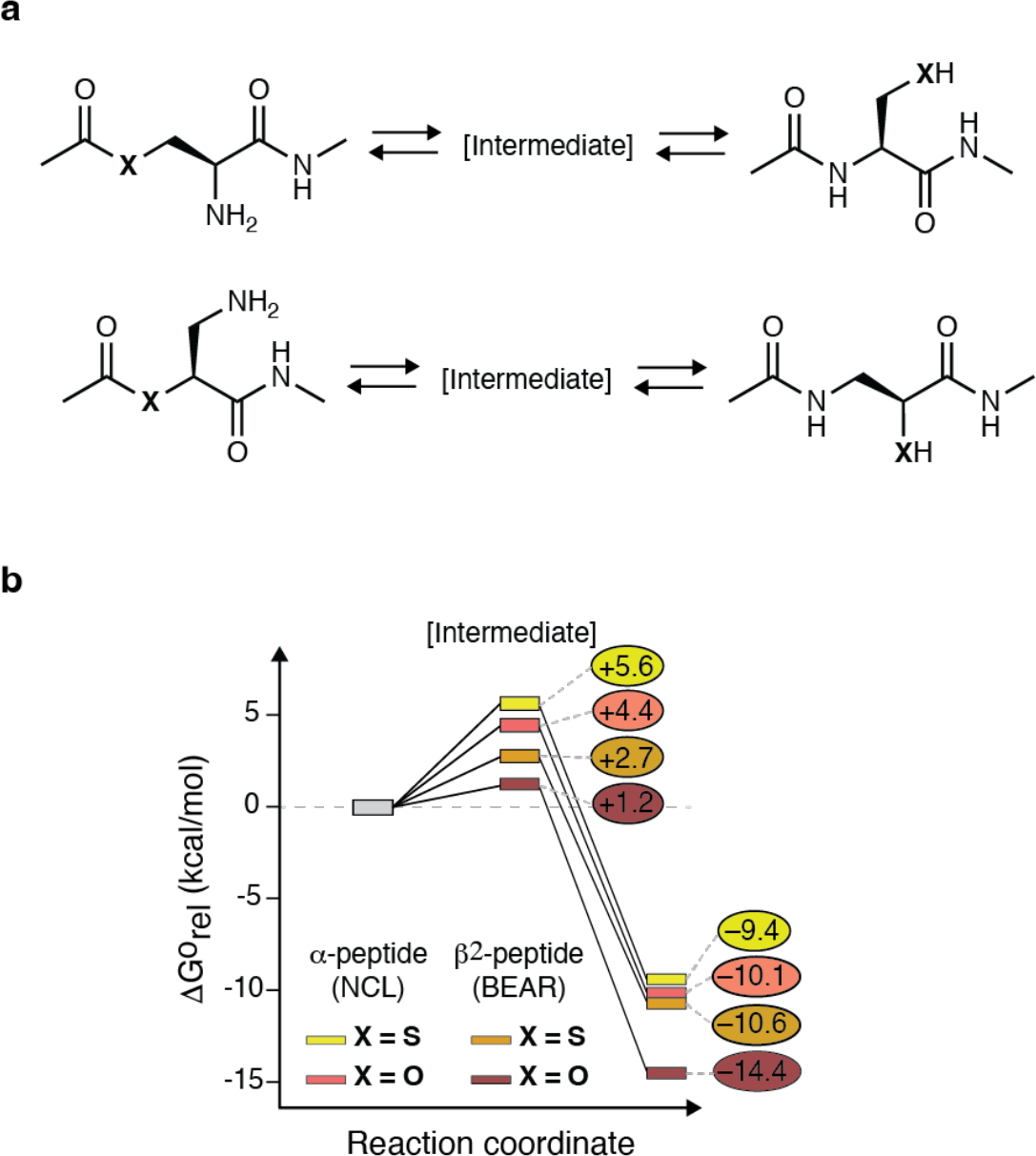
Computational analysis suggests that BEAR reactions of isoS and isoC are thermodynamically favorable. **(A)** We compared the energetics of intramolecular cyclization reactions that represent the second step of the canonical NCL reaction (top) and the single step BEAR reaction (bottom). Both were evaluated using simplified N-Me amide derivatives of Ac-Ser, Ac-Cys, Ac-isoS, and Ac-isoC. The species [Intermediate] represents the cyclic intermediate formed during each acyl-transfer step. **(B)** Relative thermodynamics (ΔGº) of the NCL and BEAR reactions established using DFT (ωB97M-V/6-311++G(3d,f)(3d,p), with a CPCM solvent model). Each reaction can proceed through two diastereomeric intermediates; only the lowest energy species is listed. All species were assumed to be uncharged (i.e., not protonated on basic sites and not deprotonated on acidic sites) for this analysis. For additional details and a full list of the computed values, see the Extended Data 1.

These calculations revealed that all four rearrangement reactions were thermodynamically favorable, as expected, but also identified unexpected differences as a function of heteroatom location and identity. Overall, the acyl rearrangement of isoS was characterized by the most favorable free energy profile, followed by isoC, Ser, and finally Cys. Free energy parameters for the cyclic intermediates formed in each case followed the same general trend (**Figure 2B**). Due to the high favorability for isoS- and isoC-based BEAR reactions, we reasoned that both could take place spontaneously in the context of an intact protein or translation product.

### Model studies support the feasibility of oxo-ester-based BEAR reactions

Next, we synthesized two model systems to experimentally establish the feasibility of *O*-to-*N* and *S*-to-*N* BEAR reactions under conditions that mimic those found in cells. We chose to photo-mask the β^2^-amine nucleophile, as photo-unmasking is compatible with cells and photo-masked diaminopropionic acid (pm-DAP) **1** (**Figure 1A**) is a known substrate for the PylRS variant DAPRS^70,74^. We hypothesized that the analogous photo-masked isoserine (pm-isoS, **2**) and photo-masked isocysteine (pm-isoC, **(±)-3**) would also act as DAPRS substrates and facilitate BEAR reactions in cells or after protein purification. In the model systems, pm-isoS **2** or pm-isoC (±**)-3** were acylated with *N*-acetylglycine to generate ester **4** or thioester **(±)-5**, respectively (**Figure 3A, Supplementary Fig. 1A, 1B**). We also synthesized the authentic products **6** and **(±)-7** that would result from the BEAR reaction of ester **4** or thioester **(±)-5**, respectively (**Figure 3A, Supplementary Fig. 1C, 1D)**.

**Figure 3.**
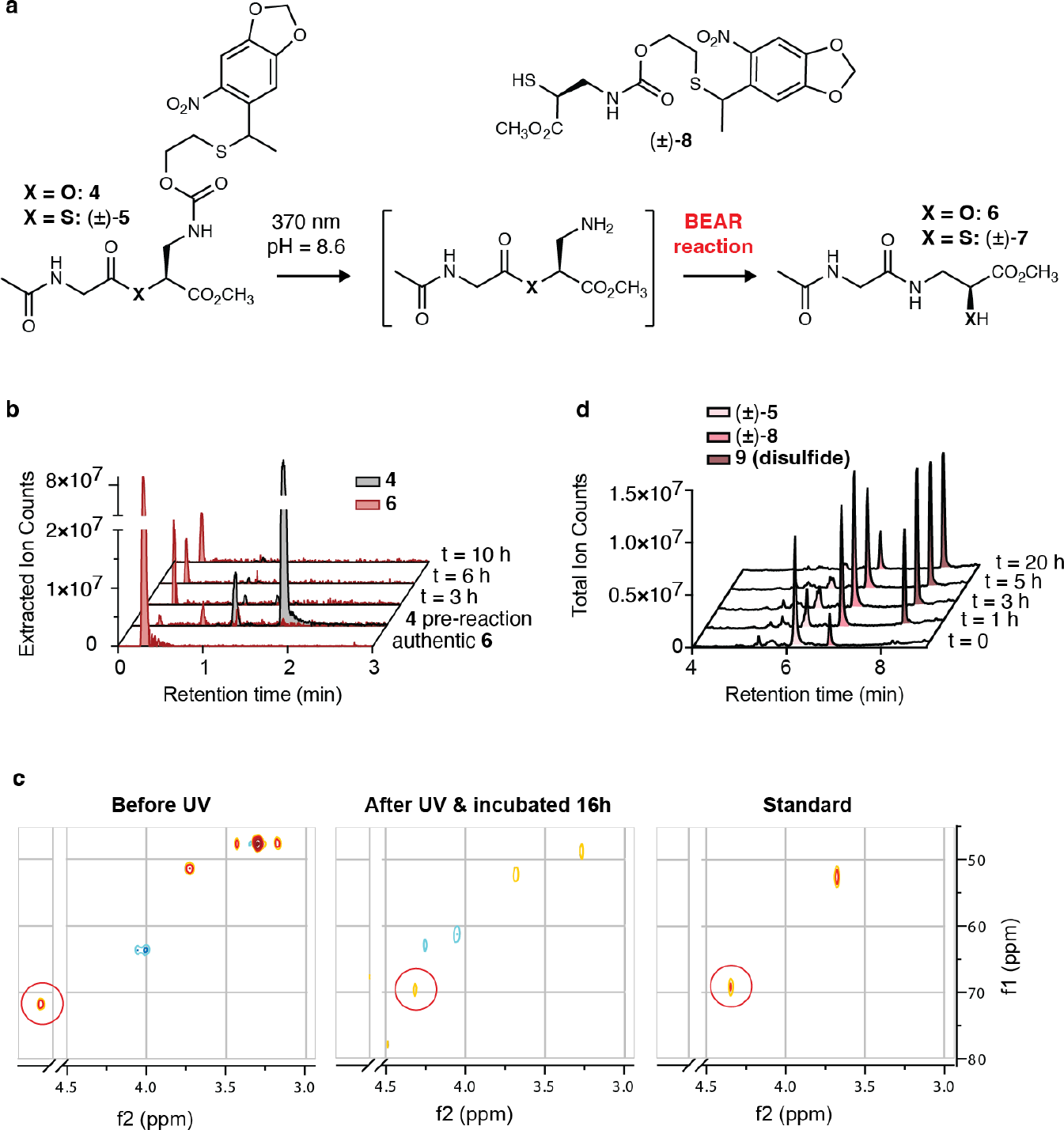
Model studies confirm the feasibility of oxo-ester-based BEAR reactions. (A) Model dipeptides **4** and **(±)-5** used to evaluate the feasibility of *in vivo* BEAR reactions to generate β^2^-linkages within intact proteins. Irradiation at 370 nm was expected to reveal a reactive amine nucleophile and initiate rearrangement into β^2^-peptides **6** or (±)-**7**. (B) Extracted ion chromatograms (EIC) corresponding to the mass of unmasked product **6** (MH+, 219.10 Da) in time-dependent LC-MS analyses following photo-unmasking of ester **4**. Reactions were performed in 10% D_2_O, 90% Reaction Buffer (50 mM sodium phosphate buffer, 1.5 mM DTT, pH 8.6) at 3 mM **4** and subjected to 370 nm irradiation for 2 min, followed by incubation at 37 ºC for the indicated time. The extracted chromatogram following BEAR reaction matches that of authentic standard **6**. (C) Multiplicity-edited Heteronuclear Single Quantum Correlation (HSQC) spectroscopy of ester **4**, authentic standard **6**, and the reaction mixture resulting from photo-unmasking and incubation of ester **4**. In the experimental spectra, the cross-peak (red circle) corresponding to the C_α_-H_α_ correlation in ester **4** shifts upfield in both dimensions, ultimately matching the authentic standard **6**. (D) Thioester **(±)-5** was dissolved in 10% D_2_O, 90% Reaction Buffer at 3 mM, but immediately began to hydrolyze to **(±)-8** with a half-life of less than 1 h. The resulting thiol dimerized into disulfide **9**, which became the major product after 5 h.

With these model systems in hand, we set out to establish the feasibility of an ester-or thioester-based BEAR reaction. For the ester-based BEAR reaction, a solution of 3 mM **4** in 50 mM phosphate buffer and 1.5 mM DTT (pH 8.6, containing 10% D_2_O) was irradiated at 370 nm (11 W) for 2 min and then incubated at 37°C for 3-16 h. The course of the reaction was monitored by LC-MS (**Figure 3B**) and product identity was confirmed by 1D- and 2D-NMR (**Figure 3C, Supplementary Fig. 2**). Photo-unmasking of ester **4** upon irradiation and subsequent incubation resulted in complete conversion to a new product within 3 h whose retention time and mass matched that of the authentic BEAR product **6** (**Figure 3B**). 1D ^1^H-NMR of the reaction mixture showed the resonance corresponding to Hα in ester **4** began to disappear at 3 h. By 16 h, the peak disappeared and was replaced by a new upfield peak whose chemical shift matched that of Hα in the authentic standard **6** (**Supplementary Fig. 2**). The identity of the BEAR product was further confirmed by 2D-NMR; multiplicity-edited HSQC spectroscopy illustrated that the Hα-Cα cross-peak of ester **4** was cleanly transformed into an upfield-shifted peak corresponding to the rearranged product **6** and matching the authentic standard (**Figure 3C**).

A different reaction course was observed when thioester **(±)-5** was incubated under identical conditions. Evaluation of the reaction components using LC-HRMS before UV irradiation revealed rapid hydrolysis of the thioester (t_1/2_ < 1 hour) to form thiol **(±)-8** and generation of the analogous disulfide **9** (**Figure 3D**). Although the hydrolytic lability of thioesters, as well as oxoesters, within folded proteins is likely sequence- and structure-dependent, the rapid hydrolysis of thioester **5** suggested that BEAR reactions of oxoesters might be more straightforward to implement than those of thioesters.

### Incorporation of photo-masked isoserine 2 into proteins *in vivo*

Given the feasibility of BEAR reactions in model systems, we next determined whether pm-isoS **2** or pm-isoC **(±)**-**3** could be incorporated into proteins *in vivo*. We synthesized photo-masked isoserine (pm-isoS, **2**) and photo-masked isocysteine (pm-isoC, **(±)-3**) and the previously reported α-amino acid, pm-DAP (**1, Supplementary Fig. 3**). We first determined whether pm-isoS **2** or pm-isoC **(±)-3** could be introduced at a single position within superfolder green fluorescent protein (sfGFP). Once incorporated, irradiation would unmask the nucleophilic amine that would promote the same intramolecular BEAR reaction observed in the model system to generate a β^2^-peptide linkage *in situ* (**Figure 4A**). We focused initially on position N150 of sfGFP, which faces outward from the folded β-barrel and has previously been shown to incorporate pm-DAP **1**^70^.

**Figure 4.**
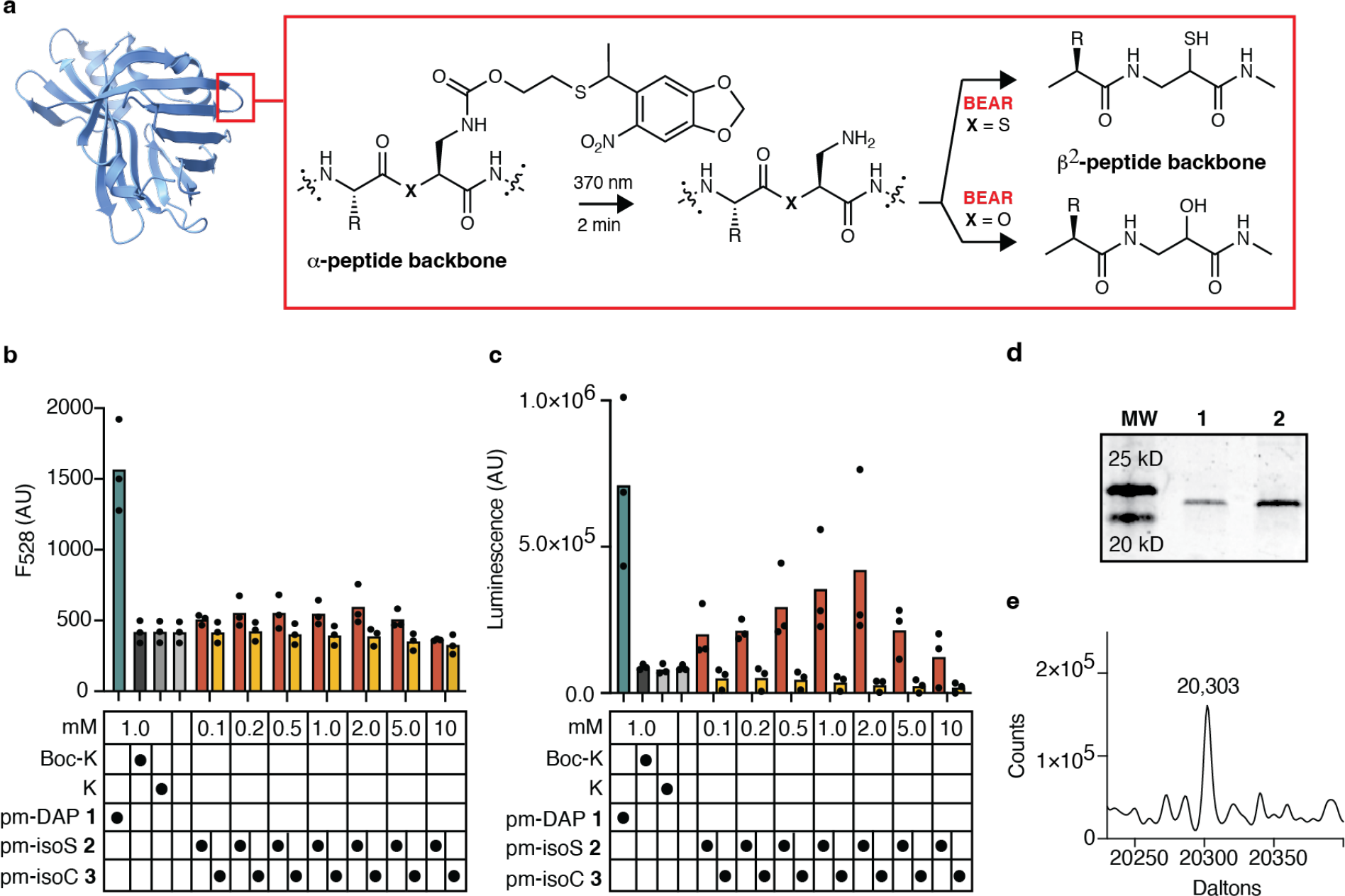
Incorporation of photo-masked isoserine 2 into proteins *in vivo*. (A) Scheme illustrating the utility of pm-isoS **2** or pm-isoC **(±)-3** for intra-protein BEAR reactions. Cellular incorporation of pm-isoS **2** or pm-isoC **(±)-3** generates a protein containing an internal ester or thioester but retains a natural peptide-like backbone. Photo-unmasking generates an intermediate capable of a BEAR reaction to generate a β^2^-peptide linkage. PDB ID 7SNS. (B) Plot showing emission at 528 nm (F_528_, emission maximum of GFP) at the 24 h time-point as a function of the concentration of pm-DAP **1**, pm-isoS **2**, or pm-isoC **(±)-3** as well as BocK, K, and in the absence of any added substrate. (C) Luminescence at the 18 h time point as a function of the concentration of pm-DAP **1**, pm-isoS **2**, or pm-isoC **(±)-3** as well as BocK, K, and in the absence of any added substrate. (D) Coomassie-stained gel showing SDS-PAGE of NanoLuc affinity-purified from growths of DH10B *E. coli* expressing DAPRS and NanoLuc with a TAG codon inserted between positions G159 and V160 and supplemented with 1 mM pm-DAP **1** or pm-isoS **2**. (E) Deconvoluted mass following LC-MS analysis of the affinity-purified NanoLuc G159-pm-isoS-V160 construct. The mass expected for NanoLuc containing pm-isoS **2** at a single position is 20,301 Da; the observed mass is 20,303.

To begin, we transformed DH10B *E. coli* cells with two plasmids, one encoding DAPRS/tRNA^Pyl^_CUA_ (pBK-DAPRS) and a second encoding the N150TAG variant of sfGFP (p15A-sfGFP-N150TAG). Cells were grown in the presence of increasing concentrations (0.1 -10 mM) of pm-isoS **2** or pm-isoC **(±)-3**. Control cultures supplemented with 1 mM pm-DAP **1** (positive control) or 1 mM Lys or Boc-Lys (negative controls) were run alongside, as well as a final control culture lacking any substrate (**Figure 4B**). All growths were performed initially at 200 μL scale at 37°C in a 96-well plate format. The change in OD_600_ and fluorescence at 528 nm (F_528_) were evaluated over 24 h after induction with 0.2% arabinose when the OD_600_ of the growths reached 0.6 (**Supplementary Fig. 4**).

Examination of the fluorescence at 528 nm of each growth at the 24 h time-point suggested robust incorporation of pm-DAP **1**, as expected, along with modest incorporation of pm-isoS **2**. There was no evidence for incorporation of pm-isoC **(±)-3** (**Figure 4B**). In the case of pm-isoS **2**, emission due to GFP increased in a concentration-dependent manner with the highest GFP expression observed at pm-isoS **2** concentrations between 0.2 mM and 2 mM. However, the overall level of sfGFP expression was, at best, only 1.4-fold over background. We concluded that a more sensitive reporter protein was needed to optimize the incorporation of pm-isoS **2** prior to evaluating how to best implement a BEAR reaction and establish the desired β^2^-peptide linkage. The lower apparent activity of pm-isoC **(±)-3** is consistent with the previously reported lower activity of α-thiol acids as substrates for PylRS variants *in vitro*^7^. A purified sample of sfGFP containing pm-DAP **1** was used to optimize conditions needed for photo-unmasking (**Supplementary Fig. 5)**.

We selected Nanoluciferase (NanoLuc)^75^ as a more sensitive reporter, as its readout (i.e., light) is generated catalytically. We prepared a series of reporter plasmids with NanoLuc in place of sfGFP and TAG codons at six different positions. Of the six plasmids, two contained either a mutated or inserted TAG codon near the NanoLuc N-terminus (G2TAG and G2-TAG-V3^76^), two contained inserted TAG codons within a loop (G159-TAG-V160^77^ and G103-TAG-V104), and two placed mutated TAG codons within a β-sheet (T130-TAG and G131-TAG). To validate the plasmids, BL21(DE3) *E. coli* were co-transformed separately with each NanoLuc reporter plasmid and pEVOL-PylRS^7^, which encodes the *Methanosarcina alvus* PylRS/tRNA^Pyl^_CUA_ pair. In each case, the cells were grown for 18 h in the presence of either 1 mM BocK or α-OH-BocK^4^ (positive controls), or Lys or no substrate (negative controls). After 18 h, luminescence was determined using the Nano-Glo® Luciferase Assay System (Promega). These experiments clearly identified the reporter encoding G159-TAG-V160 NanoLuc as possessing the largest dynamic range; the signal generated for growths containing α-OH-BocK was more than three orders of magnitude higher than growths to which no substrate had been added. All other constructs show a dynamic range of two orders of magnitude or less (**Supplementary Fig. 6**).

With an optimized detection system in hand, we next probed for *in vivo* incorporation of pm-isoS **2** or pm-isoC **(±)-3**. We transformed DH10B *E. coli* with pET15a-NanoLuc(G159-TAG-V160) as well as pBK-DAPRS encoding DAPRS/tRNA^Pyl^_CUA_, and grew the cells in the presence of 0.1 to 10 mM of pm-isoS **2** or pm-isoC **(±)-3**. Control cultures contained either 1 mM pm-DAP **1** (positive control) or 1 mM Lys, BocK, or no substrate (negative controls). The luminescence intensity of each growth was measured after 18 h as described above (**Figure 4C**). Even with this more sensitive reporter, the photo-masked pm-isoC **3** showed no evidence of incorporation, with no signal above background at any substrate concentration. However, addition of pm-isoS **2** led to robust concentration-dependent NanoLuc expression, with up to 5.2-fold enhanced signal above background at a concentration of 2 mM **2** (**Figure 4C**). LC-MS analysis of intact NanoLuc purified from a preparative-scale growth (**Figure 4D**) showed a mass corresponding to the introduction of pm-isoS **2** at a single position (**Figure 4E**).

### BEAR reaction occurs with full conversion in NanoLuc

We then subjected a purified sample of NanoLuc containing pm-isoS **2** to optimized unmasking conditions established using sfGFP containing pm-DAP **1** (**Supplementary Fig. 5**) Although intact protein LC-MS analysis provided evidence that pm-isoS **2** had been introduced into NanoLuc and unmasked (**Supplementary Fig. 7**), this technique alone could not confirm that a BEAR reaction had occurred, as the BEAR reaction is fundamentally an isomerization. To confirm that the BEAR reaction occurred, purified NanoLuc samples from growths supplemented with pm-DAP **1** or pm-isoS **2** were unmasked and evaluated by high-resolution tryptic peptide mapping (**Figure 5A**) alongside a synthetic authentic tryptic standard **10** containing the anticipated β^2^-substituted product of the BEAR reaction. Samples A and B (positive controls) were purified from growths supplemented with pm-DAP **1** (Sample A) and subsequently irradiated at 370 nm (Sample B). Samples C and D were purified from growths supplemented with pm-isoS **2** (Sample C) and subsequently irradiated (Sample D). Similarly, Sample E was isolated from a growth supplemented with pm-isoS **2**, but in this case the cells were irradiated with 370 nm light prior to cell lysis and NanoLuc purification. Sample F followed the same procedure as sample E with an additional 370 nm irradiation step after purification. Sample G contained material from a NanoLuc expression to which no substrate was added as a negative control. Finally, in an effort to increase the fraction of isolated NanoLuc containing pm-isoS **2**, we modified DAPRS to contain three mutations previously identified as beneficial for α-hydroxy acid selectivity in PylRS^8^. These mutations did not improve the incorporation of pm-isoS **2** into NanoLuc grown in either DH10B, BL21, or C321.ΔA.exp^78^ *E. coli* (**Supplementary Fig. 8**).

All samples were denatured and digested with trypsin prior to LC-MS/MS. Trypsin cleaves NanoLuc to generate a fragment containing residues V155 through R165 with an additional residue between G159 and V160 (**Figure 5B**). As anticipated, when isolated from Samples A and B, this tryptic fragment contained primarily either pm-DAP **1** (Sample A) or its unmasked product DAP (Sample B) (**Supplementary Fig. 9**). Specifically, 44.64% of the NanoLuc in Sample A contained pm-DAP **1** between G159 and V160, while 11% contained DAP indicating a low level of unmasking in ambient light. With regard to Sample B, 0.06% of the isolated NanoLuc contained pm-DAP **1**, while 57.53% contained DAP. Comparison of the compositions of Samples A and B demonstrate essentially quantitative unmasking (**Figure 5C**). Samples A and B both contained approximately 30% Gln in place of pm-DAP **1** due to misincorporation (**Supplementary Fig. 9**).

NanoLuc isolated from growths supplemented with pm-isoS **2** generated analogous tryptic fragments. Tryptic mapping of Sample C revealed that 2.09% of the isolated NanoLuc contained pm-isoS **2** between G159 and V160 while 1.89% contained the mass of isoS, again indicating some unmasking in ambient light. Tryptic mapping of Sample D, however, revealed the isolated NanoLuc contained 0.0% pm-isoS **2** and 9.5% of a product whose mass corresponded to isoS.

To determine if the tryptic NanoLuc product isolated from Sample D whose mass corresponded to isoS contained the anticipated BEAR product, we synthesized peptide **10**, which contains the appropriate β^2^-residue between G159 and V160 (**Figure 5B**). When co-injected on LC-MS/MS, peptide **10** co-eluted with the NanoLuc tryptic product isolated from Sample D whose mass corresponded to isoS (**Figure 5D**), and the two materials generated identical MS/MS spectra (**Figure 5E** and **5F**). Notably, the tryptic product from Sample D whose mass corresponded to isoS does not co-elute with a synthetic peptide containing Ser between G159 and V160 (**Supplementary Fig. 10**). We hypothesize that the lower level of pm-isoS **2** in Sample C is due to ester hydrolysis during sample preparation or LC-MS/MS analysis; photo-unmasking of Sample D provides protection against hydrolysis by creating a stable amide bond. Together, these experiments provide conclusive evidence that a protein containing pm-isoS **2** can be unmasked quantitatively and subsequently undergo a spontaneous intramolecular BEAR reaction to install a β^2^-peptide in the NanoLuc backbone.

**Figure 5.**
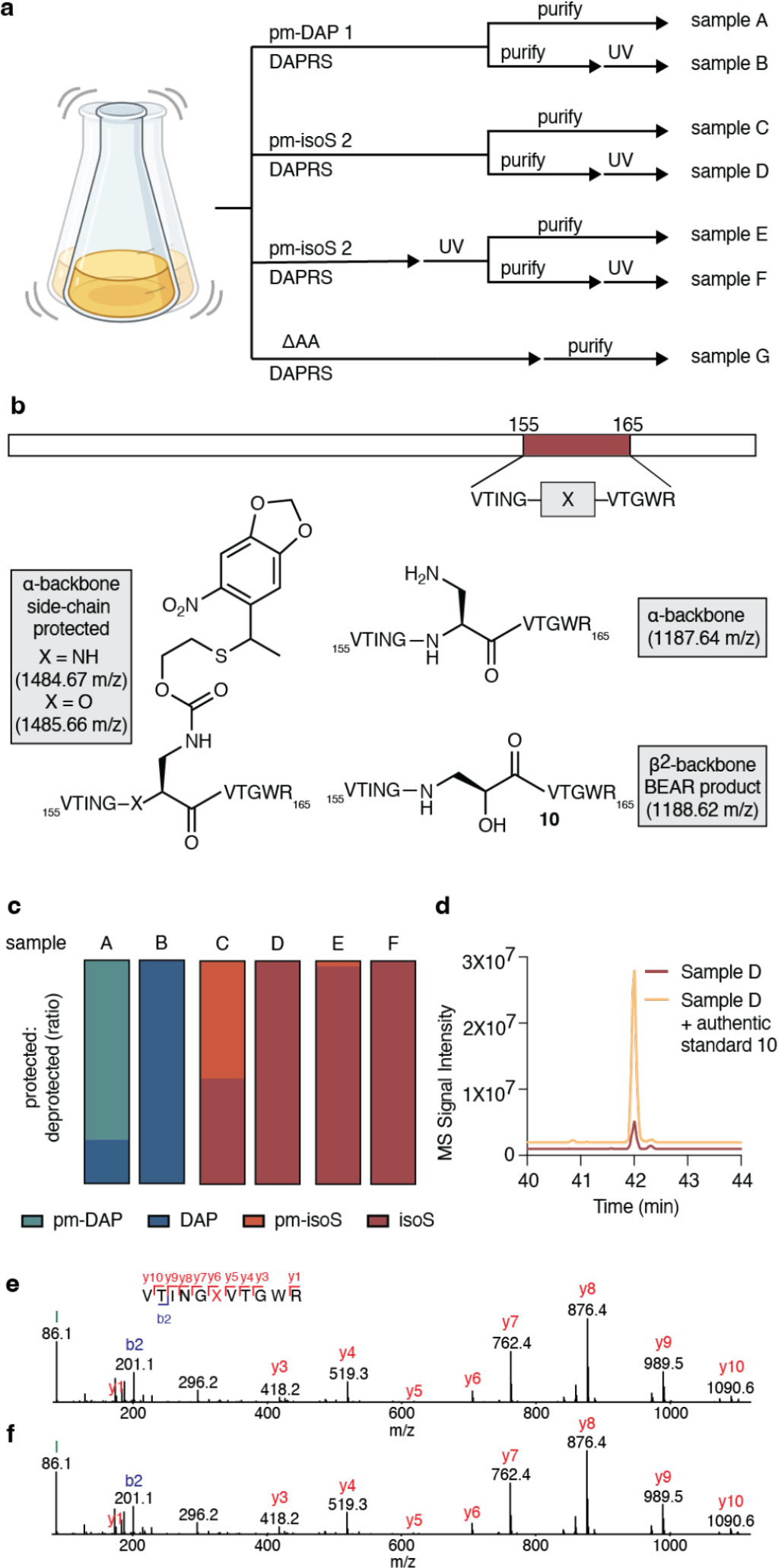
BEAR reactions proceed in quantitative yield. (A) Seven samples (Samples A-G) of NanoLuc were isolated from DH10B *E. coli* growths containing DAPRS and the monomers shown either before or after photo-unmasking. All were analyzed by high-resolution tryptic mapping. (B) The NanoLuc tryptic fragment comprising residues 155-165 (including the expected site of incorporation of pm-DAP **1** or pm-isoS **2** between native G159 and V160 was used for analysis. The mass of this fragment prior to unmasking, as well as before (α-peptide backbone) and after (β^2^-peptide backbone) BEAR cyclization, is shown. **Note that for peptide mapping, V160 in the native protein will be labeled as V161 and X will be in position 160. (C) Peptide mapping reveals partial photo-unmasking from ambient light prior to 370 nm irradiation, and full photo-unmasking upon direct exposure to 370 nm light. (D) Trace illustrating the coelution of the Sample D NanoLuc tryptic fragment comprising residues 155-165 (including the expected site of incorporation of pm-isoS **2**) alongside authentic standard **10**. The small later-eluting peak contains α-Ser in position **X** (**Supplementary Fig. 10**). (E) The MS/MS profile of Sample D after tryptic digestion is identical to that of (F) authentic standard **10**, providing conclusive evidence for a successful BEAR reaction..

Analysis of the remaining samples provides insight into an optimal BEAR workflow for establishing β^2^-linkages within intact proteins. Tryptic samples derived from Sample E, where cells were irradiated prior to NanoLuc purification, contained 0.15% pm-isoS **2** and 5.79% of the BEAR product, suggesting that higher levels of the rearranged β^2^-amino acid BEAR product are achieved with post-purification unmasking. Consistent with this notion, Sample F, which was irradiated before and after NanoLuc purification, contained 0.0% of pm-isoS **2** and 9.35% of the β^2^-amino acid BEAR product (**Figure 5C**). We note that all NanoLuc samples expressed from growths containing pm-isoS **2** contained significant levels of Ala (∼33%) and Gln (∼22%) in place of pm-isoS **2** (**Supplementary Fig. 9**). No evidence of pm-DAP **1**, DAP, pm-isoS **2**, or isoS was found in negative control Sample G (**Supplementary Fig. 9)**.

## Conclusions

There is great interest in strategies to achieve the programmed cellular synthesis of polymers whose monomers are not simply α-amino acids^1^. The availability of such molecules would provide otherwise non-existent opportunities to expand and evolve protein and polypeptide structure and function and develop improved therapeutic agents. However, despite the availability of multiple orthogonal aminoacyl-tRNA synthetase enzymes for non-canonical α-amino acids^79,80^, and multiple engineered ribosomes with finite levels of orthogonality^68^, polymers containing even the simplest non-α-amino acid—a β^2^-amino acid—have remained largely out of reach^63^. The challenge is two-fold. One challenge is the ribosome, whose ability to promote efficient bond-forming reactions *in vivo* to and from anything other than an α-amino acid or α-hydroxy acid is unknown^67^. The second challenge is the availability of ribosome substrates—acylated tRNAs. Although recent work has expanded the diversity of monomers accepted by certain archaeal aaRS enzymes^4,7^, there is still not a single efficient and orthogonal enzyme that acylates tRNA with a β-amino acid, let alone a monomer that differs from an α-amino acid by more than a CH_2_ group.

Here we bypass these challenges by reframing the problem of cellular hetero-oligomer synthesis in the language of chemistry. Rather than relying on direct reactions of β-amino acid monomers within the ribosome active site, the strategy reported here relies on light-promoted and proximity-guided intramolecular rearrangements that effectively “edit” the protein backbone post-translationally. In this case, the intramolecular rearrangement converts an α-backbone directly into a β-backbone. As far as we know, this report represents the first example in which an expanded backbone β-amino acid linkage has been introduced in an orthogonal fashion into a protein in a cell. Although we demonstrate this concept using a single photo-masked nucleophile, the strategy we describe can be expanded to alternative nucleophiles, side chain geometries, and unmasking strategies capable of installing even more diverse backbones into proteins expressed in cells and with temporal control. We anticipate that BEAR reactions, such as those reported here, will enable the straightforward cellular synthesis and ultimately the evolution of extended backbone-protein hybrids for research and development of next-generation biomaterials and protein therapeutics.

## Author contributions

Study conception and design: L.T.R., A.C., M.B.F., S.J.M., A.S.; preparation of materials: L.T.R., C.S., T.L.D., N.X.H., S.Z., D.A.D., N.W.; data collection:

L.T.R., C.S., T.L.D., B.S., N.X.H., S.Z., N.W., M.B.F.; analysis and interpretation of results:

L.T.R., C.S., T.L.D., B.S., Z.Z., M.B.F., S.J.M., A.S.; and manuscript preparation: L.T.R., C.S., T.L.D., M.B.F., S.J.M., A.S.

## Acknowledgements

We are grateful to members of the Schepartz, Miller, and Chatterjee labs for insightful comments and suggestions, and to Jason Chin (MRC) for sharing materials. We also thank Hasan Celik for assistance with NMR spectroscopy and acknowledge NIH S10OD024998 for spectrometer support. This work was supported by the NSF Center for Genetically Encoded Materials (C-GEM; CHE 2002182). L.T.R. and T.L.D. were supported by the NSF Graduate Research Fellowship Program (DGE-1752814 and DGE-2139841, respectively). C.K.S. is supported by the Miller Institute for Basic Research in Science, University of California Berkeley. A.S. is a Chan-Zuckerberg Biohub-San Francisco Investigator and an ARC Innovation Investigator. Created with BioRender.com.

## Data Availability

The data in this study are available from the corresponding authors upon request.

