## Supplementary Information for "Backbone extension acyl rearrangements enable cellular synthesis of proteins with internal β^2^-peptide linkages"

### DFT Analysis Methods

#### Methods to support Figure 2

***Computational Studies.*** For each compound that was studied, molecular mechanics methods (Macromodel, OPLS4 force field) were used to minimize a starting population of 10,000 conformers that was initially generated using the Conformational Search tool in the Schrödinger Maestro 2022-2 environment. The geometries of all unique conformers that were found to be within 5 kcal/mol of the global minimum were then optimized again using DFT at the B3LYP-D3/6-31G** level. The global minima resulting from this round of computation were subjected to a second geometry optimization and vibrational spectrum calculation in QChem v. 6.02 (B3LYP-D3/6-31G**) to determine the zero-point energies, enthalpy corrections, and the internal entropy values at 298.15 K. All compounds were confirmed to have no imaginary vibrational frequency modes. For all species, refined electronic energy calculations were performed on the optimized geometries using an expanded basis set (ωB97M-V/6-311++G(3df, 3pd)). For calculations including solvation, a CPCM model was used with Bondi van der Waals radii and a SAS solvent probe radius of 1.4 Å. In these cases, solvation was introduced only into the final electronic energy calculations. Tabulated energy and entropy values appear in **Extended Data 1** along with atomic coordinates of the optimized structural minima used for each calculation.

### Synthesis Methods

#### General synthetic details

All reactions were carried out under ambient atmosphere unless otherwise noted. Room temperature (rt or RT) is defined as 21–23 °C. All reagents were obtained from commercial sources and used without further purification, unless otherwise noted. Cyclopentanone was distilled under vacuum and stored in a freezer prior to use. *N,N*-diisopropylethylamine (DIPEA) was distilled over CaH under a nitrogen atmosphere prior to use or dispensed from a Sigma-Aldrich Sure/Seal$^{TM}$ container (99.5%, purified via redistillation) under a nitrogen atmosphere. Toluene was dried over alumina and dispensed under argon from a Glass Contour Seca Solvent Purification System. Deionized water was used for reactions, extraction solutions, and reverse phase chromatography. Acetonitrile and ethyl acetate used for chromatography were High-Performance Liquid Chromatography (HPLC) grade, while hexanes used for chromatography was certified ACS grade. Methanol used for chromatography was either certified ACS grade or electronic grade. Hexanes used for recrystallization was HPLC grade. High-resolution mass spectrometry (HRMS) was either conducted by the Chemical and Biophysical Instrumentation Center in the Chemistry Department at Yale University using a Waters Xevo Quadrupole Time-of-Flight (Q-TOF) high resolution mass spectrometer using electrospray ionization (ESI), on an LTQ FT-ICR mass spectrometer equipped with an electrospray ionization source (Finnigan LTQ FT, Thermo Fisher Scientific, Waltham, MA) operated in either positive or negative ion mode, or collected on an Agilent 6530 Accurate-Mass Q-TOF LC-MS. For crude data analysis, ultra-performance liquid chromatography-mass spectrometry (UPLC/MS) was performed with a Waters Acquity UPLC/MS instrument equipped with a reverse phase BEH C18 column (1.7 mm particle size, 2.1 x 50 mm), a dual atmospheric pressure chemical ionization (APCI)/electrospray ionization (ESI) mass spectrometry detector, and a photodiode array detector. Analytical thin-layer chromatography (TLC) was performed using 60 Å Silica Gel F254 pre-coated plates (0.25 mm thickness). TLC plates were visualized by irradiation with a UV lamp or with Hanessian's stain (cerium ammonium molybdate). Flash chromatography was performed using either a Biotage® Isolera One purification system equipped with a 10, 25, 50, 100 or 340 g SNAP Ultra (HP Sphere, 25 µm silica) cartridge for normal phase column chromatography, and 12, 30, 60, or 120 g SNAP-C18 columns for reverse phase column chromatography; or a Teledyne ISCO Combiflash NextGen 300+ equipped with a 4, 12, or 24 g RediSep columns for normal phase column chromatography. Reverse phase HPLC purification was performed on an Waters LC Prep 150 system equipped with a Waters 2998 UV photodiode array detector, a Waters 2707 autosampler,  and an Waters Fraction Collector III using a preparative reverse phase C18 column (CSH C18 19 x 150 mm OBD Column 5 µm). The mobile phase for HPLC was water with 0.1% (v/v) trifluoroacetic acid (solvent A) and acetonitrile with 0.1% (v/v) trifluoroacetic acid (solvent

B), at a flow rate of 20 mL/min. Analytes were collected based on their absorbance at 280 nm or 214 nm.

Routine $^{1}$H nuclear magnetic resonance (NMR) spectra were recorded on Agilent or Bruker 400, 500, or 600 MHz spectrometers at ambient temperature unless otherwise stated. Chloroform-*d* ($CDCl_3$, with or without 1% v/v tetramethylsilane [TMS]) and dimethylsulfoxide-$d_6$ (DMSO-$d_6$) were purchased from Cambridge Isotope Laboratories and used without further purification. $CDCl_3$ (with or without 1% v/v TMS) was stored at ambient temperature over 5Å molecular sieves; DMSO-$d_6$ ampules were used immediately upon opening. Methanol-$d_4$ was purchased from Sigma Aldrich. Spectra were processed using MestReNova 14.2.0 using the automatic phasing, polynomial baseline correction capabilities, and zero-filling. Splitting was determined using the automatic multiplet analysis function with intervention as necessary. Spectral data are reported as follows: chemical shift (multiplicity [singlet (s), broad singlet (br s), doublet (d), triplet (t), quartet (q), pentet (p), multiplet (m), doublet of doublets (dd), doublet of doublet of doublets (ddd), doublet of triplet of doublets (dtd), doublet of doublet of doublet of doublets (dddd), doublet of triplets (dt), triplet of doublets (td), etc.], coupling constant (Hz), integration). The abbreviation "app" denotes an apparent multiplicity (e.g., "app t" denotes an apparent triplet). Chemical shifts are reported in parts per million (ppm, δ), and coupling constants are reported in Hz. $^{1}$H resonances are referenced to solvent residual peaks for $CDCl_3$ (7.26 ppm) or DMSO-$d_6$ (2.50 ppm) or to TMS (0.00 ppm) or to methanol-$d_4$ (3.31 ppm)[1]. Routine $^{13}$C NMR spectra were recorded on Agilent or Bruker 400 (101), 500 (126), or 600 (151) MHz spectrometers with protons fully decoupled. $^{13}$C Resonances are reported in ppm relative to solvent residual peaks for $CDCl_3$ (77.16 ppm) or DMSO-$d_6$ (39.52 ppm) or methanol-$d_4$ (49.00 ppm)

**Abbreviations**

Ac – acetyl
Ac-Gly-OH – *N*-acetylglycine
APCI – Atmospheric Pressure Chemical Ionization
aq. – aqueous
Boc – *tert*-butoxycarbonyl
ca. – *circa* (approximately)
CaH – calcium hydride
Calcd, calcd., or calc. – calculated
$CDCl_3$ – chloroform-*d*
conc. – concentrated
COSY – Correlated Spectroscopy
CV – column volume(s)
DCM – dichloromethane (methylene chloride)
DI – deionized
DIPEA – *N,N*-diisopropylethylamine (Hünig's base)

DMF – *N,N*-dimethylformamide
DMSO-$d_6$ – dimethyl sulfoxide-$d_6$
equiv – equivalent(s)
ESI – Electrospray Ionization
$Et_2O$ – diethyl ether
EtOAc – ethyl acetate
Fmoc – fluorenylmethyloxycarbonyl
Fmoc-OSu – *N*-(*9H*-fluoren-9-ylmethoxycarbonyloxy)succinimide
h – hour(s); min – minute(s)
H-Cys(Trt)-OH – *S*-trityl-L-cysteine
HATU – hexafluorophosphate azabenzotriazole tetramethyl uronium
HBTU – hexafluorophosphate benzotriazole tetramethyl uronium
HCl – hydrochloric acid
Hex – hexanes
HPLC – High-Performance Liquid Chromatography
HRMS – High-Resolution Mass Spectrometry
HSQC – Heteronuclear Single Quantum Coherence
isoC – isocysteine (3-amino-2-mercaptopropanoic acid)
isoS – isoserine (3-amino-2-hydroxypropanoic acid)
LiCl – lithium chloride
M – molar (i.e., mol/L); N – normal
MeCN – acetonitrile
MeOH – methanol (methyl alcohol)
$MgSO_4$ – magnesium sulfate
$Na_2SO_4$ – sodium sulfate
$NaHCO_3$ – sodium bicarbonate (sodium hydrogen carbonate)
NaOAc – sodium acetate
NaOH – sodium hydroxide
$NEt_3$ – triethylamine
NMR – Nuclear Magnetic Resonance
pm – photo-masked
*p*TsOH·$H_2O$ – *p*-toluenesulfonic acid monohydrate
Q-TOF – Quadrupole Time-of-Flight
RP-HPLC – Reverse Phase High-Performance Liquid Chromatography
rt or RT – room temperature (21–23 °C)
tBu – *tert*-butyl
TFA – trifluoroacetic acid
TIPS – triisopropylsilane
TLC – Thin-Layer Chromatography
TMS – tetramethylsilane

TMSCl - chlorotrimethylsilane
Trt – triphenylmethyl (trityl)
TrtOH – triphenylmethanol
uncorr. – uncorrected
UPLC/MS – Ultra-Performance Liquid Chromatography/Mass Spectrometry
v/v – volume/volume (i.e., mL/100 mL)
w/w – weight/weight (i.e., g/100 g)

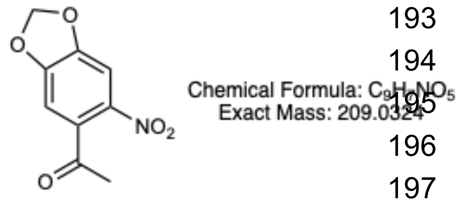

**12**

**1-(6-nitrobenzo[*d*][1,3]dioxol-5-yl)ethan-1-one (12)**

*(Following a procedure adapted from Huguenin-Dezot et al., 2019[2])*

A solution of 3',4'-(methylenedioxy)acetophenone (**11**) (3.283 g, 20 mmol) in glacial acetic acid(12.8 mL) was added dropwise to a 500 mL three-neck round-bottom flask containing conc. $HNO_3$ (27.2 mL, 70% w/w) at 0 ˚C over 1 h. The reaction mixture was maintained at 0 ˚C during the addition and for an additional 1 h with stirring under a $N_2$ atmosphere. The mixture was then warmed to 40 ˚C and stirred for an additional 3 h., the mixture was cooled to RT and poured into crushed ice in a beaker. A yellow precipitate appeared, which was stirred for 15 min and then filtered. The yellow solid was washed with water (3 × 50 mL) and dried under vacuum. The crude yellow solid was then purified by recrystallization (THF/*n*-hexane) to obtain the title compound (**12**) as yellow crystals (2.968 g, 71%): $R_f$ = 0.5 ($SiO_2$, 100% $CH_2Cl_2$). **$^1$H NMR** (500 MHz, chloroform-*d*) δ 7.54 (s, 1H), 6.75 (s, 1H), 6.18 (s, 2H), 2.49 (s, 3H); **$^{13}$C{$^1$H} NMR** (126 MHz, chloroform-*d*) δ 199.6, 153.1, 149.3, 140.6, 135.6, 106.6, 105.3, 104.0, 30.6.; **MS** (ESI) *m/z* calc. for $C_9H_7NO_5^+$ [MH]+: 210.04, found: 210.21

**1-(6-nitrobenzo[*d*][1,3]dioxol-5-yl)ethan-1-ol ((±)-13)**

*(Following a procedure adapted from Huguenin-Dezot et al., 2019[2])*

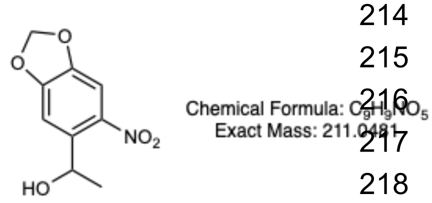

**13**

Ketone **12** (1.045 g, 5.0 mmol, 1.0 equiv) was dissolved in THF (25 mL) and $NaBH_4$ (472.87 mg, 12.5 mmol, 2.5 equiv) was added. The resultant suspension was stirred overnight at RT and then quenched with 1 N aq. HCl until gas evolution ceased. The mixture was poured into brine, the organic components were extracted with $CH_2Cl_2$ (3x), and the combined organic layers were dried over sodium sulfate and concentrated under reduced pressure. The residue was purified via flash column chromatography (100% $CH_2Cl_2$) to give the desired product as a yellow solid (1.046 g, 4.96 mmol, 99%). $R_f$ = 0.30 ($SiO_2$, 100% $CH_2Cl_2$). **$^1$H NMR** (500 MHz, chloroform-*d*) δ 7.46 (s, 1H), 7.27 (s, 1H), 6.13 – 6.10 (m, 2H), 5.46 (q, J = 6.3 Hz, 1H), 1.54 (d, J = 6.3 Hz, 3H); **$^{13}$C{$^1$H} NMR** (126 MHz, chloroform-*d*) δ 152.6, 147.1, 141.7, 139.1, 106.5, 105.3, 103.1, 65.9, 24.3; **HRMS** (ESI) *m/z* calc. For $C_9H_9NO_5$+ [MH]+: 212.0553, found: 212.0508.

**5-(1-bromoethyl)-6-nitrobenzo[*d*][1,3]dioxole ((±)-14)**

*(Following a procedure adapted from Huguenin-Dezot et al., 2019[2])*

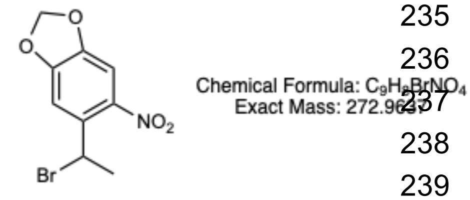

**14**

Alcohol **(±)-13** (1.03 g, 4.88 mmol, 1.0 equiv) was dissolved in dry $CH_2Cl_2$ (24.5 mL) and cooled to 0 °C under a nitrogen atmosphere in a 250 mL round-bottom flask. After 10 min, $PBr_3$ (0.185 mL, 1.95 mmol, 0.4 equiv) was added dropwise at 0 °C. Next, the mixture was brought to rt and stirred continuously for 16 h with aluminum foil wrapped around the flask. The reaction was judged to be complete by TLC analysis ($SiO_2$, TLC eluent: 100% $CH_2Cl_2$), cooled to 0 °C, quenched by the addition of 1 M aq. NaOH (2 mL), and warmed to rt to stir for 30 min under a nitrogen atmosphere. After the quenching was complete, saturated aq. $NaHCO_3$ solution was added (20 mL). The contents were loaded into a separatory funnel, the aqueous phase was discarded, and the organic phase was washed sequentially with further saturated aq. $NaHCO_3$ solution (1 × 20 mL) and brine (2 × 20 mL). The organic layer was separated, dried over anhydrous $Na_2SO_4$, and concentrated *in vacuo* to obtain a yellow solid. The crude product was purified by flash chromatography on $SiO_2$ [eluent: 100% $CH_2Cl_2$] to obtain pure alkyl halide **(±)-14** (1.33 g, 4.87 mmol, >99%) as yellow crystals. $R_f$ = 0.30 ($SiO_2$, 100% $CH_2Cl_2$). **$^1$H NMR** (500 MHz, chloroform-*d*) δ 7.34 (s, 1H), 7.26 (s, 1H), 6.12 (t, J = 1.3 Hz, 2H), 5.89 (q, J = 6.8 Hz, 1H), 2.03 (d, J = 6.9 Hz, 2H); **$^{13}$C{$^1$H} NMR** (126 MHz, chloroform-*d*) δ 152.1, 147.7, 141.7, 134.8, 108.8, 105.1, 103.3, 42.8, 27.7; **HRMS** (ESI) *m/z* calcd. for $C_9H_8BrNO_4^+$ [MH]+: 273.9709, found: 273.9664.

**2-((1-(6-nitrobenzo[*d*][1,3]dioxol-5-yl)ethyl)thio)ethan-1-ol ((±)-15)**

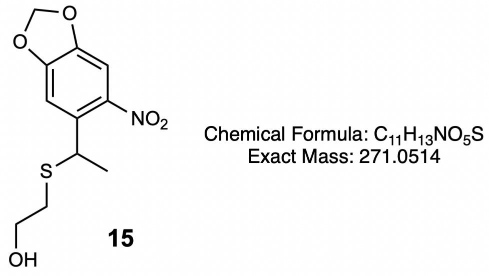

**15**

*(Following a procedure adapted from Huguenin-Dezot et al., 2019[2])*

An aqueous solution of NaOH was freshly prepared (0.5 M, 1.4 g in 7 mL of $H_2O$, 20.0 mmol, 5.8 equiv), loaded into a 100 mL round-bottom flask, and degassed by bubbling through a stream of nitrogen gas at rt. After 30 min, mercaptoethanol (0.254 mL, 3.6 mmol, 1.05 equiv) was added to the flask and degassing was continued for a further 15 min. Separately, freshly prepared alkyl halide **(±)-14** (0.941 g, 3.43 mmol, 1.0 equiv) was dissolved in 1,4-dioxane (11.5 mL) in a 100 mL round-bottom flask wrapped in aluminum foil and degassed by bubbling through a stream of nitrogen gas for 15 min. The degassed solution of **(±)-14** in 1,4-dioxane was added dropwise into the flask containing the aq. NaOH and mercaptoethanol solution, at rt under a positive pressure of nitrogen gas. The mixture was left stirring for 16 h at rt in the dark under a nitrogen atmosphere, after which time the reaction was judged to be complete by TLC. The mixture was then evaporated under reduced pressure to remove the volatile organic components. The resultant yellow aqueous mixture was then extracted with EtOAc (2 x 15 mL) and the combined organic phases were washed with a saturated $NH_4Cl$ solution (1 x 15 mL), followed by brine (3 x 15 mL). The organic layer was then separated, dried over anhydrous $Na_2SO_4$, filtered, and evaporated to dryness to obtain a yellow oil. The product was purified by flash chromatography on $SiO_2$ (eluent: EtOAc/*n*-hexane = 3:7) to obtain alcohol **(±)-15** as yellow crystals (0.920 g, 3.39 mmol, >99%). $R_f$ = 0.33 ($SiO_2$ plate, EtOAc/*n*-hexane = 3:7). **$^1$H NMR** (500 MHz, chloroform-*d*) δ 7.28 (s, 1H), 7.27 (s, 1H), 6.11 – 6.09 (m, 2H), 4.79 (q, J = 6.9 Hz, 1H), 3.69 – 3.58 (m, 2H), 2.62 – 2.49 (m, 2H), 1.56 (d, J = 6.9 Hz, 3H); **$^{13}$C{$^1$H} NMR** (126 MHz, chloroform-*d*) δ 151.8, 146.7, 143.1, 135.8, 107.8, 104.5, 102.8, 60.7, 38.1, 34.7, 22.9; **MS** (ESI) *m/z* calcd. for $C_{11}H_{13}NO_5S$+ [MH]+: 272.06, found: 272.17.

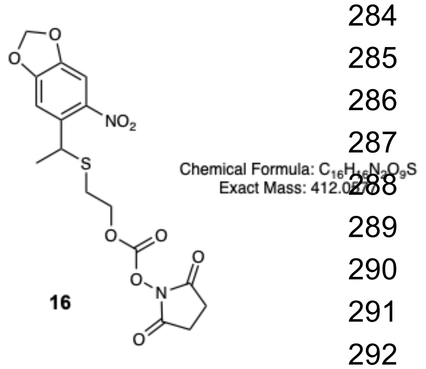

**2-((1-(6-nitrobenzo[*d*][1,3]dioxol-5-yl)ethyl)thio)ethyl 2,5-dioxopyrrolidine-1-carboxylate ((±)-16)**

*(Following a procedure adapted from Huguenin-Dezot et al., 2019[2])*

A 20 mL microwave vial was charged with alcohol **(±)-15** (0.542 g, 1.99 mmol, 1.0 equiv) dissolved in dry $CH_3CN$ (6 mL) under nitrogen atmosphere, and dry DIPEA (1.04 mL, 5.98 mmol, 3.0 equiv) was added. In a second 20 mL microwave vial *N,N'*-disuccinimidyl carbonate (0.716 g, 2.8 mmol, 1.4 equiv), was added with dry $CH_3CN$ (6 mL) under nitrogen atmosphere (the mixture did not completely dissolve). The contents of the first vial were transferred to the second vial dropwise (12 mL total) under nitrogen atmosphere in the dark. After 30 min, all components were dissolved and left to stir at rt for 16 hours as a homogenous yellow solution. The reaction was judged to be complete by TLC analysis ($SiO_2$ plate, EtOAc/*n*-hexane = 3:7) after this time. Carbonate **(±)-16** was immediately carried to the next step without further purification: $R_f$ = 0.12 ($SiO_2$ plate, EtOAc/*n*-hexane = 3:7).

**L-isoserine methyl ester (H-isoS-OMe, 17)**

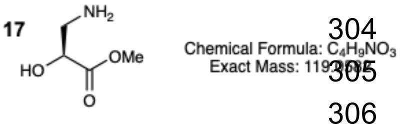

To a solution of isoserine ((*S*)-3-amino-2-hydroxypropanoic acid, 139 mg, 1.3 mmol, 1.0 equiv) in anhydrous methanol (3 mL) was added TMSCl (0.33 mL, 2.6 mmol, 2 equiv). After 24 h, the mixture was concentrated *in vacuo* to provide methyl ester **17** as a colorless solid (155 mg, 1.3 mmol, >99% yield). **$^1$H NMR** (600 MHz, methanol-$d_4$) δ = 4.44 (dd, *J* = 8.1, 4.0 Hz, 1H), 3.79 (s, 3H), 3.33 – 3.29 (m, 2H), 3.13 (dd, *J* = 13.1, 8.1 Hz, 1H); **$^{13}$C{$^1$H} NMR** (151 MHz, methanol-$d_4$) δ = 172.9, 68.3, 53.1, 43.0; **MS** (ESI) m/z calcd. for $C_4H_{10}NO_3^+$ [2MH]+: 239.12, found: 239.12

##### *N*-photo-masked-L-isoserine methyl ester (pm-isoS-OMe, 18)

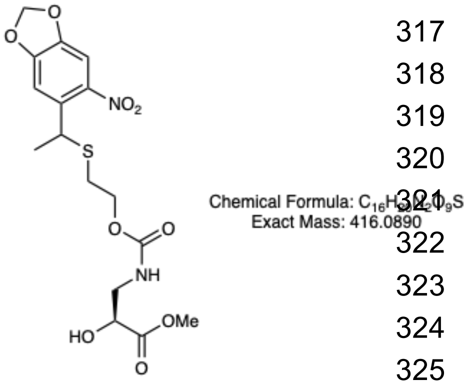

**18**

Methyl ester **17** (51 mg, 0.5 mmol, 1.1 equiv) was added in one portion to a solution of carbonate **(±)-16** (0.44 mmol, 1.0 equiv, prepared as described above) in dry $CH_3CN$ (2.2 mL) under nitrogen gas and stirred for 16 h at rt. After this time, the reaction was judged to be complete by LC-MS and the reaction mixture was diluted with water containing 0.1% TFA, filtered, and purified by RP-HPLC while shielded from light. The residue was then subjected to preparative reverse phase HPLC on a Waters Prep 150 LC System [CSH C18 19 x 150 mm OBD Column 5 µm; gradient: 5-50% $H_2O$-$CH_3CN$ (+0.1% TFA throughout) mobile phase over 16 min] to obtain the title compound. Purity of resulting fractions was determined by LC-MS and pure fractions were immediately evaporated of solvent, affording photo-masked compound **18** as a yellow oil and a mixture of diastereomers (62 mg, 34% yield). The product was immediately used in the subsequent reaction to avoid possible decay. A $^1$H NMR is provided for reference. **$^1$H NMR** (600 MHz, methanol-$d_4$) δ = 7.35 – 7.29 (m, 2H), 6.15 – 6.12 (m, 2H), 4.78 – 4.72 (m, 1H), 4.22 (t, *J* = 6.2 Hz, 1H), 4.02 (dd, *J* = 11.9, 6.1 Hz, 2H), 3.73 (s, 2H), 2.68 (s, 3H), 2.66 – 2.44 (m, 2H), 1.54 (dt, *J* = 6.8, 3.1 Hz, 3H)**;** **MS** (ESI) m/z calcd. for $C_{16}H_{21}N_2O_9S^+$ [MNa]+: 439.08 found: 439.32

##### *N*-photo-masked-*O*-(*N*-acetylglycyl)-L-isoserine methyl ester (pm-isoS(Ac-Gly)-OMe, 4)

To a solution of *N*-acetylglycine (68 mg, 0.6 mmol, 10 equiv) in dry $CH_2Cl_2$ (5 mL) was added DIC (0.09 mL, 0.6 mmol, 10 equiv) and NMI (0.05 mL, 0.6 mmol, 10 equiv). The resulting mixture was added to a flask containing photo-masked compound **18** (24 mg, 0.06 mmol, 1.0 equiv). After stirring for 0.5 h, the reaction mixture was concentrated *in vacuo*. The crude material was dissolved (95% water, 5% MeCN, 0.1% TFA), filtered through a 0.2 micron PTFE filter, and purified by reverse phase HPLC on a Waters Prep 150 LC System [CSH C18 19 x 150 mm OBD Column 5 µm; gradient: 5-50% $H_2O$-$CH_3CN$ (+0.1% TFA throughout) mobile phase over 30 min] to afford a waxy yellow solid as a mixture of diastereomers (16 mg, 53% yield). **$^1$H NMR**

**4**

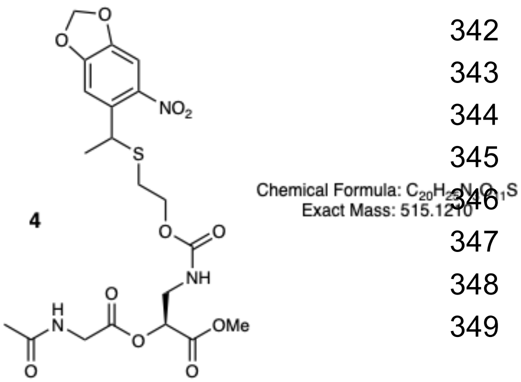

(600 MHz, methanol-$d_4$): δ 7.31 (d, *J* = 1.5 Hz, 2H), 6.14 – 6.12 (m, 2H), 5.14 – 5.09 (m, 1H), 4.75 (dd, *J* = 7.0, 3.7 Hz, 1H), 4.10 – 4.00 (m, 4H), 3.74 (s, 3H), 3.62 – 3.52 (m, 2H), 2.63 – 2.50 (m, 2H), 2.00 (s, 3H), 1.54 (d, *J* = 6.9 Hz, 3H); **$^{13}C\{^1H\}$ NMR** (151 MHz, methanol-$d_4$) δ = 173.8, 170.6, 170.1, 158.5, 153.4, 148.5, 144.8, 136.7, 108.7, 105.4, 104.7, 73.3, 65.3, 65.1, 53.1, 42.4, 41.8, 39.9, 31.4, 23.1, 22.3; **HRMS** (ESI) m/z calcd. for $C_{20}H_{26}N_3O_{10}S_2^+$ [MH]+: 516.1283, found: 516.1296

**2-(2-oxo-1-oxa-4-thiaspiro[4.4]nonan-3-yl)acetic acid ((±)-20)**

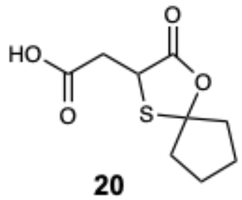

**20**

Chemical Formula: $C_9H_{12}O_4S$
Exact Mass: 216.0456

*(Following a procedure adapted from Dose & Seitz, 2004[3])*
To a two-neck round-bottom flask equipped with a stir bar was added 2-mercaptosuccinic acid **(±)-19** (7.00 g, 46.6 mmol, 1.0 equiv) and *p*-toluenesulfonic acid monohydrate (0.89 g, 4.66 mmol, 0.1 equiv). The flask was charged with benzene (100 mL), and a Dean-Stark apparatus was attached and charged with additional benzene and a small aliquot of water. The Dean-Stark apparatus was affixed with a reflux condenser, and the system was sealed with rubber septa and placed under a nitrogen atmosphere. The suspension was briefly stirred (< 2 min), and distilled cyclopentanone (6.2 mL, 69.9 mmol, 1.5 equiv) was added to the flask through a septum via syringe. The reaction was placed into a preheated oil bath and refluxed for 4.5 h, at which point the solution had turned clear. The reaction was cooled to room temperature and concentrated *in vacuo*. The crude residue was dissolved in saturated aq. $NaHCO_3$ and washed with DCM (3x). The combined organic extracts were discarded and the aqueous phase was acidified to pH 1-2 with concentrated aq. HCl and extracted with DCM (3x). The combined organic phases were then dried over $Na_2SO_4$, filtered, and concentrated *in vacuo.* The dried material was then purified by recrystallization in EtOAc/Hexanes to afford thioketal **(±)-20** as an off-white solid (3.17 g, 14.7 mmol, 31% yield). **$^1H$ NMR** (400 MHz, $CDCl_3$) δ 10.94 (br s, 1H), 4.41 (dd, *J* = 9.1, 4.1 Hz, 1H), 3.25 (dd, *J* = 17.5, 4.1 Hz, 1H), 2.87 (dd, *J* = 17.6, 9.1 Hz, 1H), 2.35 – 2.18 (m, 2H), 2.09 – 1.91 (m, 2H), 1.90 – 1.72 (m, 4H). *[In reasonable accord with the literature characterization[3]]*

***N*-photo-masked DL-isocysteine thioketal ((±)-21)**

*(Following a procedure adapted from Dose & Seitz, 2004, for a related compound[3])*
To a round-bottom flask equipped with a stir bar was added thioketal **(±)-20** (3.17 g, 14.7 mmol, 1.0 equiv) and dry toluene (59 mL). The flask was sealed with a rubber septum and placed under a nitrogen atmosphere. Triethylamine (2.45 mL, 17.6 mmol, 1.2 equiv) was added slowly at room temperature through the septum via syringe, and the solution was stirred for 30 min. The solution was then cooled to 0 °C with an ice bath, and diphenylphosphoryl azide (3.79 mL, 17.6

mmol, 1.2 equiv) was added slowly through the septum via syringe. The solution was removed from the ice bath and stirred for 3.5 h at room temperature, during which time it gradually darkened to a caramel color. The reaction mixture was then opened to air via a vent needle in the septum, warmed to 85 °C in a preheated oil bath, and stirred with a vent needle until gas evolution ceased (ca. 30 min). The flask was cooled to 60 °C, at which point alcohol **(±)-15** (3.78 g, 13.9 mmol, 0.95 equiv) was added slowly via syringe. The solution was then stirred overnight at 60 °C, cooled to room temperature, and diluted with EtOAc. The mixture was then washed sequentially with saturated aq. $NaHCO_3$ (1x), water (1x), 10% aq. citric acid (1x), and brine (1x). The organic phase was dried over $Na_2SO_4$, filtered, and concentrated *in vacuo*. The crude residue was purified twice via normal phase automated flash column chromatography (0-100% EtOAc/Hexanes to 0-10% MeOH/EtOAc), twice by reverse phase automated flash column chromatography (0-100% MeCN/$H_2O$), and lyophilized to afford photo-masked thioketal **(±)-21** as a viscous yellow oil and as a mixture of diastereomers (2.60 g, 5.37 mmol, 36% yield).

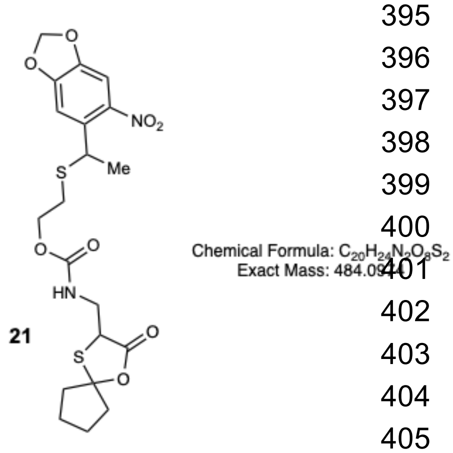

**21**

*Specific Chromatography Details*

Normal Phase 1 – Biotage® SNAP Ultra 100 g; 0% EtOAc/Hex for 2.0 CV, 0-42% over 4.2 CV, 42% for 0.8 CV, 42-100% over 5.7 CV, 100% over 14.6 CV, then 0-10% MeOH/EtOAc over 1.0 CV, 10% for 11.4 CV.

Normal Phase 2 – Biotage® SNAP Ultra 340 g; 0% EtOAc/Hex for 2.0 CV, 0-46% over 11.0 CV, 46% for 0.1 CV, 46-48% over 0.1 CV, 48% for 0.3 CV, 48-100% over 3.6 CV, 100% for 1.0 CV, then 0-10% MeOH/EtOAc over 1.0 CV, 10% for 0.9 CV.

Reverse Phase 1 – Biotage® SNAP Ultra C18 120 g; 0% MeCN/$H_2O$ for 1.0 CV, 0-1% over <0.1 CV, 1% for 0.2 CV, 1-13% over 1.8 CV, 13% for 0.7 CV, 13-48% over 5.1 CV, 48% for 0.3 CV, 48-59% over 1.6 CV, 59% for 2.8 CV, 59-100% over 6.1 CV, 100% for 1.0 CV.

Reverse Phase 2 – Biotage® SNAP Ultra C18 120 g; 0% MeCN/$H_2O$ for 1.0 CV, 0-59% over 7.1 CV, 59% for 1.7 CV, 59-100% over 4.8 CV, 100% for 1.0 CV.

**$^1$H NMR** (600 MHz, $CDCl_3$) δ 7.27 (s, 1H), 7.26 (s, 1H), 6.11 (app s, 1H), 6.09 (app s, 1H), 5.47 (app q, $J$ = 6.3 Hz, 1H), 4.93 – 4.75 (m, 1H), 4.24 (app td, $J$ = 6.1, 2.9 Hz, 1H), 4.19 – 4.03 (m, 2H), 3.65 (app t, $J$ = 6.3 Hz, 2H), 2.70 – 2.39 (m, 2H), 2.30 – 2.15 (m, 2H), 2.04 – 1.93 (m, 2H), 1.89 – 1.81 (m, 2H), 1.77 (m, 2H), 1.53 (d, $J$ = 6.9 Hz, 3H); **$^{13}$C{$^1$H} NMR** (151 MHz, $CDCl_3$) δ 173.20, 173.19, 156.0, 152.0, 146.8, 143.3, 136.1, 108.0, 104.6, 103.0, 95.8, 65.1, 48.29, 48.27, 42.64, 42.61, 42.5, 41.34, 41.32, 38.9, 30.4, 23.8, 23.6, 23.04, 23.02; $^1$H-$^1$H COSY and multiplicity-edited $^1$H-$^{13}$C HSQC spectra are also provided; **HRMS** (ESI) *m/z*: $[M+H]^+$ Calcd for $C_{20}H_{25}N_2O_8S_2^+$ 485.1047; Found 485.1045.

#### *N*-photo-masked *S*-(*N*-acetylglycyl) DL-isocysteine methyl ester (pm-$^{DL}$isoC(Ac-Gly)-OMe, (±)-5)

To a round-bottom flask equipped with a stir bar was added was added photo-masked thioketal **(±)-21** (576 mg, 1.19 mmol, 1 equiv) and MeOH (8 mL). Solid NaOH (99 mg, 2.48 mmol, 2.1 equiv) was then added to the flask in one portion. The reaction was vigorously stirred for 1 h, after which the solution was diluted with water and acidified to pH ~1 with concentrated aq. HCl. The solution was then extracted with DCM (3x), and the combined organic layers were washed with brine, dried over $Na_2SO_4$, filtered, and concentrated *in vacuo* to provide the wet intermediate thiol **(±)-8** as a viscous oil (503 mg). A portion of the crude material (460 mg, 1.06 mmol) was used in the next step without further purification. A crude $^1$H NMR spectrum of this intermediate compound is provided for reference.

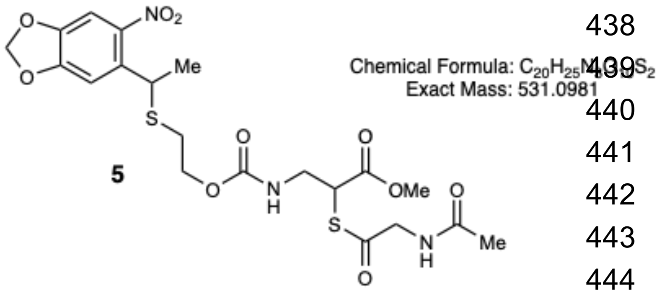

**5**

To a vial equipped with a stir bar was added *N*-acetylglycine (150 mg, 1.28 mmol, 1.2 equiv) and HATU (485 mg 1.28 mmol, 1.2 equiv). The vial was then charged with DMF (3.5 mL), and dry DIPEA (0.56 mL, 3.19 mmol, 3 equiv) was added dropwise with stirring. The solution was pre-stirred for 10 min, then transferred to a vial containing thiol **(±)-8** (460 mg, 1.06 mmol, 1 equiv) and a stir bar. The resultant solution was then stirred for 2.5 h and partitioned between EtOAc (ca. 60 mL) and 5% aq. LiCl (ca. 60 mL). The organic phase was then washed sequentially with 5% aq. LiCl (3x), 10% aq. citric acid (1x), saturated aq. $NaHCO_3$ (2x), and brine (1x). The organic phase was dried over $Na_2SO_4$, filtered, and concentrated *in vacuo*. The resultant material was then purified via normal phase automated flash column chromatography (20-100% EtOAc/Hexanes) and lyophilized to afford photo-masked thioester **(±)-5** as a yellow foam and as a mixture of diastereomers (324 mg, 0.610 mmol, 51% yield over two steps, uncorrected for diverted material).

*Specific Chromatography Details*

Normal Phase 1 – Biotage® SNAP Ultra 50 g; 20% EtOAc/Hex for 1.0 CV, 20-100% over 12.0 CV, 100% for 6.1 CV.

**$^1$H NMR** (500 MHz, $CDCl_3$) δ 7.28 (d, *J* = 2.6 Hz, 1H), 7.26 (d, *J* = 2.6 Hz, 1H), 6.27 (app d, *J* = 6.8 Hz, 1H), 6.11 (app t, *J* = 1.1 Hz, 1H), 6.09 (app d, *J* = 1.3 Hz, 1H), 5.41 – 5.25 (m, 1H), 4.93 – 4.75 (m, 1H), 4.37 (app td, *J* = 6.1, 2.1 Hz, 1H), 4.24 (app dd, *J* = 5.8, 2.9 Hz, 2H), 4.12

(app dq, $J$ = 10.9, 6.6 Hz, 2H), 3.75 (s, 3H), 3.71 – 3.61 (m, 1H), 3.55 (app dtd, $J$ = 14.5, 5.9, 2.9 Hz, 1H), 2.69 – 2.42 (m, 2H), 2.07 (s, 3H), 1.54 (d, $J$ = 6.9 Hz, 3H).

**$^{13}C\{^1H\}$ NMR** (126 MHz, $CDCl_3$) δ 195.1, 170.49, 170.40, 156.1, 152.2, 146.9, 143.4, 136.36, 136.31, 108.2, 104.71, 104.69, 103.1, 65.4, 53.2, 49.1, 45.33, 45.30, 42.0, 39.0, 30.5, 23.10, 23.08.

$^1H$-$^1H$ COSY and multiplicity-edited $^1H$-$^{13}C$ HSQC spectra are also provided.

**HRMS** (ESI) *m/z*: $[M+Na]^+$ Calcd for $C_{20}H_{25}N_3NaO_{10}S_2^+$ 554.0874; Found 554.0853.

**Ac-Gly-isoS-OH (22)**

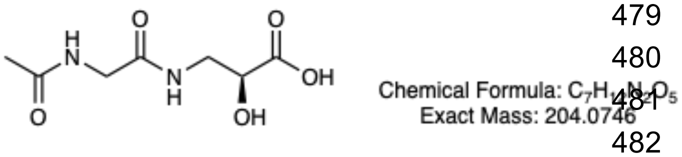

**22**

The isoserine authentic standard was prepared using solid-phase peptide synthesis (SPPS). 2-chlorotrityl chloride resin (143 mg, 1.4 mmol/g, 1.0 equiv) was transferred to a 5 mL fritted syringe and swelled with DCM for five minutes. Solvent was evacuated and replaced with a solution of (2*S*)-3-((((9*H*-fluoren-9-yl)methoxy)carbonyl)amino)-2-(*tert*-butoxy)propanoic acid (0.3 mmol, 1.5 equiv) and DIPEA (0.8 mmol, 4 equiv) in DCM and incubated for 1 hour. This solution was discarded, and the resin was washed three times with DMF. The resin was then capped with a solution of DCM:MeOH:DIPEA (17:2:1, v/v) for 30 minutes. The resin was washed three times with DMF, deprotected using 20% piperidine in DMF for 10 minutes, and washed again three times with DMF. A solution of *N*-acetylglycine (2 mmol, 10 equiv), HBTU (1.9 mmol, 9.5 equiv), and DIPEA (4 mmol, 20 equiv) in DMF was added to the syringe and incubated for 30 minutes. The resin was then washed three times with DMF and three times with DCM before being fully dried. Finally, the resin was cleaved using a solution of TFA:water (98:2) for two hours. The isolated solution was dried *in vacuo*, to yield dipeptide **22** as a colorless oil (37 mg, 93% from resin loading).

**$^1H$ NMR** (600 MHz, methanol-$d_4$) δ = 4.23 (dd, $J$ = 7.1, 4.4 Hz, 1H), 3.84 (s, 2H), 3.62 (dd, $J$ = 13.7, 4.4 Hz, 1H), 3.41 (dd, $J$ = 13.7, 7.1 Hz, 1H), 2.01 (s, 3H)

**$^{13}C\{^1H\}$ NMR** (151 MHz, methanol-$d_4$) δ = 175.6, 173.9, 172.1, 70.5, 44.0, 43.5, 22.4

**MS** (ESI) m/z calcd. for C7H13N2O5+ [MH]+: 205.08, found: 205.18

**Ac-Gly-isoS-OMe (6)**

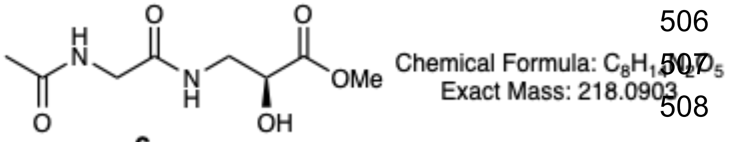

**6**

To a solution of dipeptide **22** (34 mg, 0.17 mmol, 1.0 equiv) in dry methanol (3 mL)was added TMSCl (0.04 mL, 0.33 mmol, 2.0 equiv).

After stirring for 16 h, the reaction mixture was concentrated *in vacuo* to afford the desired dipeptide methyl ester **6** as a colorless oil (36 mg, quant.) **$^{1}$H NMR** (600 MHz, methanol-*$d_4$*) δ = 4.56 (s, 1H), 4.26 (dd, *J* = 6.4, 4.8 Hz, 1H), 3.83 (d, *J* = 1.4 Hz, 2H), 3.56 (dd, *J* = 13.7, 4.8 Hz, 1H), 3.45 (dd, *J* = 13.7, 6.4 Hz, 1H), 3.35 (s, 3H), 2.00 (s, 3H). **$^{13}$C{$^{1}$H} NMR** (151 MHz, methanol-*$d_4$*) δ = 174.4, 173.9, 172.1, 70.7, 43.8, 43.5, 30.7, 22.4. **MS** (ESI) m/z calcd. for $C_8H_{15}N_2O_5^+$ [MH]+: 219.10, found: 219.10

##### *S*-Trt-DL-isocysteine (H-$^{DL}$isoC(Trt)-OH, (±)-24)

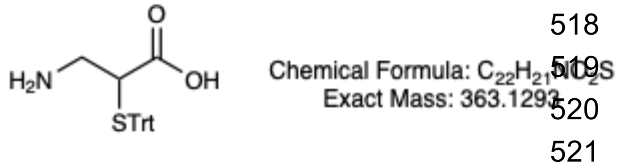

**24**

*(Following a procedure adapted from Dose & Seitz, 2004[3])* Thioketal **(±)-20**, an intermediate en route to compound **(±)-24**, was prepared as described previously (*vide supra*) with the following modifications: (i) the compound was prepared at a 59.9 mmol scale; (ii) the reaction time was shortened to 3.5 h; (iii) $MgSO_4$ was used as the drying agent, rather than $Na_2SO_4$; and (iv) the material was not subjected to purification via recrystallization. The resultant off-white solid (thioketal **(±)-20**, 1.99 g, 9.20 mmol) was used in the next step without further purification. A crude $^{1}$H NMR spectrum is provided for reference; the product peaks are in reasonable accord with the literature characterization[3].

*(Following a procedure adapted from Dose & Seitz, 2008[4])*
To a round-bottom flask equipped with a stir bar was added crude thioketal **(±)-20** (1.99 g, 9.20 mmol, 1.0 equiv) and dry toluene (40 mL). The flask was sealed with a rubber septum and placed under a nitrogen atmosphere. Triethylamine (1.5 mL, 10.8 mmol, 1.2 equiv) was added slowly through the septum via syringe at room temperature, and the solution was stirred for 30 min. The solution was then cooled to 0 °C with an ice bath, and diphenylphosphoryl azide (2.18 mL, 10.1 mmol, 1.1 equiv) was added slowly through the septum via syringe. The resultant solution was removed from the ice bath and stirred for 4 h at room temperature, during which it gradually darkened to a caramel color. The reaction mixture was then opened to air via a vent needle in the septum, warmed to 85 °C in a preheated oil bath, and stirred with a vent needle until gas evolution ceased (ca. 30 min). The flask was then removed from the oil bath and cooled to room temperature. The reaction mixture was diluted with EtOAc (40 mL), washed with saturated aq. $NaHCO_3$ (1x), and then washed with DI water (2x). The organic phase was dried over $MgSO_4$, filtered, and concentrated to provide a crude residue, which was used in the next step without further purification.

The crude residue from the previous step was taken up in 6 N aq. HCl (40 mL) and refluxed for 3 h. The reaction mixture was then cooled to room temperature and washed with $Et_2O$ (3x). The ethereal washes were discarded, and the aqueous phase was concentrated *in vacuo* to provide a dark red oil (**(±)-23**), which was used in the next step without further purification.

Trifluoroacetic acid (33 mL) was added to the crude dark red oil (**(±)-23**) from the previous step, and the solution was briefly stirred before triphenylmethanol (2.58 g, 9.9 mmol, 1.1 equiv) was added portionwise. The resultant mixture was stirred for 1 h at room temperature and concentrated *in vacuo*. The residue was then dissolved in $Et_2O$ (33 mL), and 0.2 N aq. NaOAc was added with vigorous stirring until the biphasic mixture had a pH of ~4 and a tan precipitate appeared (ca. 500 mL of 0.2 N aq. NaOAc was required). The mixture was filtered, and the precipitate was dried under vacuum. The precipitate was then collected and dissolved in acetone (33 mL) and stirred for 30 min at 40 °C. The mixture was cooled to room temperature and placed in a -20 °C freezer overnight. The mixture was filtered and the precipitate was dried under vacuum. An additional crop could be obtained by analogous recrystallization of the initial filtrate from NaOAc treatment. In total, 1.397 g (3.84 mmol) of crude trityl-protected isocysteine (**(±)-24**) was obtained from this process as an off-white solid, and a portion of the material (0.477 g, 1.31 mmol) was carried forward to the next step without further purification. A crude $^{1}H$ NMR spectrum is provided for reference; the product peaks are in reasonable accord with the literature characterization[4].

**_N_-Fmoc-_S_-Trt-DL-isocysteine (Fmoc-$^{DL}$isoC(Trt)-OH, (±)-25)**

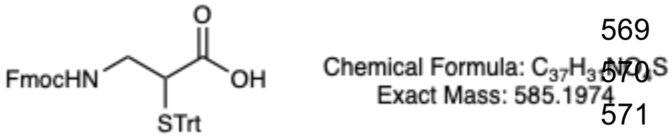

**25**

*(Following a procedure adapted from Cui et al., 2022 for the Fmoc protection of H-Cys(Trt)-OH[5])*

A round-bottom flask equipped with a stir bar was charged with crude trityl-protected isocysteine **(±)-24** (477 mg, 1.31 mmol, 1.00 equiv), 1,4-dioxane (3.3 mL), and water (1.7 mL) to afford a suspension. $NaHCO_3$ (116 mg, 1.38 mmol, 1.05 equiv) was added, and the mixture was stirred for 20 min. Fmoc-OSu (450 mg, 1.33 mmol, 1.02 equiv) was then added, and the mixture was stirred for an additional 30 min, at which point an additional portion of 1,4-dioxane (3.3 mL) and water (1.7 mL) was added. The reaction was stirred overnight at room temperature. The resultant mixture was diluted with water (10 mL), affording a cloudy white suspension. The mixture was concentrated *in vacuo* to approximately ½ of the original volume, and decanted. The decanted solid was then split into three sets and purified via reverse phase automated flash column chromatography (Set 1: 0-100% MeCN in $H_2O$; Set 2: 10-100% MeCN in $H_2O$; Set 3: 5-100% MeCN in $H_2O$ + 5% of aq. 2% formic acid buffer throughout. *Vide infra*). The residue was taken up in 10% aq. citric acid (50 mL) and vigorously stirred for 2.5 hours and extracted with EtOAc (3x30 mL). The combined organic phases were washed with brine (1x), dried over $Na_2SO_4$, filtered, and concentrated *in vacuo*. The resultant material was filtered through a syringe filter (0.2 μm pore size) with DCM/MeCN/$Et_2O$ (1:1:1, ca. 10 mL total volume), concentrated *in vacuo*, purified via reverse phase automated flash column chromatography (2-100% MeCN in $H_2O$ + 5% of aq. 2% formic acid buffer throughout), and lyophilized to afford Fmoc-$^{DL}$isoC(Trt)-OH (compound **(±)-25**, 227 mg, 0.388 mmol, 0.6% yield over five steps, uncorrected for diverted material).

*Specific Chromatography Details*

Reverse Phase Set 1 – Biotage® SNAP Ultra C18 12 g; 0-14% MeCN/$H_2O$ over 1.2 CV, 14% for 0.2 CV, 14-36% over 1.9 CV, 36% for 6.4 CV, 36-77% over 3.7 CV, 77% for 4.1 CV, 77-93% over 1.4 CV, 93% for 0.1 CV, 93-100% over 0.6 CV, 100% for 5.4 CV.
Reverse Phase Set 2 – Biotage® SNAP Ultra C18 12 g; 10-16% MeCN/$H_2O$ over 0.7 CV, 16% for 0.7 CV, 16-38% over 2.8 CV, 38% for 3.7 CV, 38-57% over 2.4 CV, 57% for 0.2 CV, 57-65% over 1.0 CV, 65% for 2.2 CV, 65-80% over 1.8 CV, 80-81% over <0.1 CV, 81% for 2.3 CV, 81-100% over 0.9 CV, 100% for 1.0 CV.
Reverse Phase Set 3 – Biotage® SNAP Ultra C18 12 g; 5-51% MeCN/$H_2O$ (+ 5% of a 2% aq. formic acid additive throughout) over 5.5 CV, 51% for 8.6 CV, 51-77% over 3.1 CV, 77% for 4.7 CV, 77-80% over 0.3 CV, 80-100% over 1.0 CV, 100% for 7.0 CV.
Final Reverse Phase – Biotage® SNAP Ultra C18 30 g; 2-80% MeCN/$H_2O$ (+ 5% of a 2% aq. formic acid additive throughout) over 9.0 CV, 80-89% over 0.4 CV, 89% for 4.9 CV, 89-100% over 0.5 CV, 100% for 2.0 CV.

**$^1$H NMR** (400 MHz, DMSO-$d_6$) δ 12.67 (s, 1H), 7.87 (d, $J$ = 7.5 Hz, 2H), 7.64 (t, $J$ = 7.8 Hz, 2H), 7.44 – 7.20 (m, 20H), 4.24 – 4.02 (m, 3H), 3.15 – 3.00 (m, 2H), 2.45 (m, 1H).
$^{13}$C NMR (151 MHz, DMSO-$d_6$) δ 172.1, 158.3, 144.0, 143.7, 140.6, 129.2, 128.0, 127.6, 127.0, 126.9, 125.3, 125.2, 120.1, 67.5, 65.5, 46.7, 46.5, 42.6, 39.9, 39.8, 39.7, 39.4, 39.2, 39.1.
A $^1$H-$^1$H COSY spectrum is provided for reference.
**HRMS** (ESI) m/z calcd. for $C_{37}H_{32}NO_4S^+$ [MNa]+: 608.1866, found: 608.1875

**Ac-Gly-$^{DL}$isoC-OH ((±)-26)**

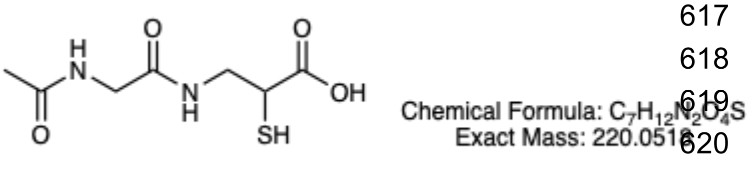

**26**

The isocysteine (isoC) authentic standard was prepared using solid-phase peptide synthesis (SPPS). 2-chlorotrityl chloride resin (36 mg, 1.4 mmol/g, 1 equiv) was transferred to a 5 mL fritted syringe and swelled with DCM for five minutes. Solvent was evacuated and replaced with a solution of Fmoc-isoC(Trt)-OH (compound **(±)-25**, 59 mg, 0.1 mmol, 2 equiv) and DIPEA (0.03 mL, 0.2 mmol, 4 equiv) in DCM and incubated for 1 hour. This solution was discarded, and the resin was washed three times with DMF. The resin was then capped with a solution of DCM:MeOH:DIPEA (17:2:1, v/v) for 30 minutes. The resin was washed three times with DMF, deprotected using 20% piperidine in DMF for 10 minutes, and washed again three times with DMF. A solution of *N*-acetylglycine (59 mg, 0.5 mmol, 10 equiv), HBTU (180 mg, 0.48 mmol, 9.5 equiv), and DIPEA (0.17 mL, 1 mmol, 20 equiv) in DMF was added to the syringe and incubated for 30 minutes. The resin was then washed three times with DMF and three times with DCM before being fully dried. Finally, the resin was cleaved using a solution of TFA:water:TIPS (95:3:2) for two hours. The isolated solution was dried *in vacuo*, and the residue was diluted with 5% acetonitrile in water, filtered through a 0.2 micron PTFE filter, and purified by reverse phase HPLC on a Waters Prep 150 LC System [CSH C18 19 x 150 mm OBD Column 5 µm; gradient: 5-50% $H_2O$-$CH_3CN$ (+0.1% TFA throughout) mobile phase over 30 min] to yield dipeptide **(±)-26** as a colorless solid (8 mg, 73% from resin loading), which was passed forward to the next step. A 1H NMR spectrum is provided for reference. **$^1$H NMR** (600 MHz, methanol-*d$_4$*) δ = 3.82 (s, 2H), 3.55 (s, 1H), 3.59 – 3.46 (m, 2H), 2.00 (s, 3H). **MS** (ESI) m/z calcd. for $C_7H_{13}N_2O_4S^+$ [MH]+: 221.06, found: 221.10

**Ac-Gly-$^{DL}$isoC-OMe ((±)-7)**

To a solution of dipeptide **(±)-26** (6 mg, 0.03 mmol, 1.0 equiv) in dry methanol (5 mL) was added TMSCl (0.01 mL, 0.05 mmol, 2.0 equiv). After stirring for 48 h, the reaction mixture was concentrated *in vacuo* to afford the desired dipeptide methyl ester **(±)-7** as a colorless solid (5.7 mg, 0.03 mmol, >99%). **$^1$H NMR** (600 MHz, methanol-*d$_4$*) δ 3.80 (s, *J* = 2.7 Hz, 2H), 3.73 (s, 3H), 3.62 – 3.59 (m, 1H), 3.57 – 3.50 (m, 2H), 2.00 (s, 3H); **$^{13}$C{$^1$H} NMR** (151 MHz, methanol-*d$_4$*) δ = 174.01, 173.89, 172.1, 53.1, 44.8, 43.5, 40.6, 22.4; **MS** (ESI) m/z calcd. for $C_8H_{15}N_2O_4S^+$ [MH]+: 235.07, found: 235.15

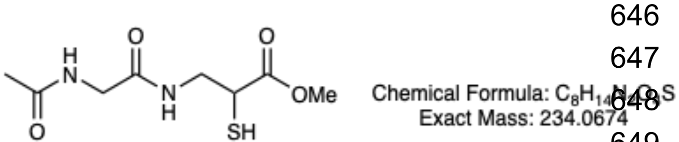

**7**

***N*$_\alpha$-Boc-*N*$_\beta$-photo-masked-L-diaminopropionic acid (27)**

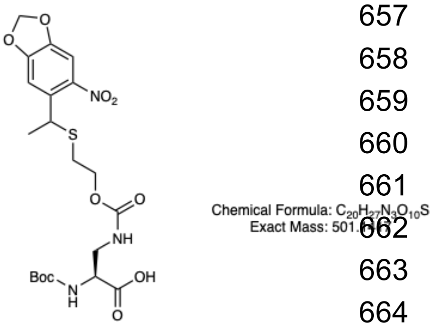

**27**

*(Following a procedure adapted from Huguenin-Dezot et al., 2019[2])*

Boc-L-Dap-OH (0.441 g, 2.16 mmol, 1.08 equiv) was added in one portion to a solution of **16** (1.99 mmol, 1.00 equiv, prepared as described above) in dry $CH_3CN$ (12 mL) under nitrogen gas and stirred for 16 h at rt. After this time, the reaction was judged to be complete by LC-MS and the reaction mixture was concentrated *in vacuo*. The residue was then subjected to preparative reverse phase HPLC on a Waters Prep 150 LC System [CSH C18 19 x 150 mm OBD Column 5 µm; gradient: 20-50% $H_2O$-$CH_3CN$ (+0.1% TFA throughout) mobile phase over 16 min] to obtain photo-masked Boc-L-Dap-OH (**27**) as a yellow solid (0.947 g, 95%) and a mixture of ~1:1 epimers. **$^1$H NMR** (500 MHz, methanol-*$d_4$*) δ 7.33 (d, J = 1.3 Hz, 1H), 7.29 (d, J = 1.2 Hz, 1H), 6.16 – 6.11 (m, 2H), 4.79 – 4.71 (m, 1H), 4.24 (t, J = 6.0 Hz, 1H), 4.11 – 3.96 (m, 1H), 3.53 (dd, J = 14.1, 4.7 Hz, 1H), 3.40 – 3.32 (m, 1H), 2.66 – 2.49 (m, 2H), 1.54 (d, J = 6.9 Hz, 3H), 1.44 (s, 9H); **$^{13}$C{$^1$H} NMR** (126 MHz, methanol-*$d_4$*) δ 173.4, 158.2, 152.8, 147.9, 144.3, 136.2, 136.2, 108.2, 104.9, 104.1, 80.2, 64.6, 54.7, 42.6, 39.4, 39.4, 30.9, 28.2, 22.6, 0.3; **MS** (ESI) *m/z* calcd. for $C_{20}H_{27}N_3O_{10}S^-$ [M]-: 500.13, found: 500.30

##### ***N*$_β$-photo-masked-L-diaminopropionic acid (pm-DAP, 1)**

*(Following a procedure adapted from Huguenin-Dezot et al., 2019[2])* Photo-masked Boc-L-Dap-OH (**27**) (0.412 g, 0.822 mmol, 1.0 equiv) was added to a dry 20 mL microwave vial and dissolved in 70% v/v TFA in $CH_2Cl_2$ (5 mL). The flask was wrapped in foil to exclude light. The yellow reaction mixture was left stirring at rt in the dark. After 2 h, the reaction was judged to be complete by LC-MS analysis. The reaction mixture was concentrated under reduced pressure to obtain a yellow gum. This residue was then subjected to preparative reverse phase HPLC on a Waters Prep 150 LC System [CSH C18 19 x 150 mm OBD Column 5 µm; gradient: 15-65% $H_2O$-$CH_3CN$ (+0.1% TFA throughout) mobile phase over 16 min] to obtain side-chain photo-masked diaminopropionic acid (pm-DAP, **1**) as a yellow solid (0.947 g, 72%) and a mixture of ~1:1 epimers. **$^1$H NMR** (500 MHz, methanol-*$d_4$*) δ 7.34 (s, 1H), 7.30 (s, 1H), 6.16 – 6.11 (m, 2H), 4.76 (q, J = 6.9 Hz, 1H), 4.08 (s, 2H), 4.10 – 4.00 (m, 1H), 3.71 (dt, J = 15.0, 4.1 Hz, 1H), 3.56 (dd, J = 15.0, 6.4 Hz, 1H), 2.63 (dt, J = 13.7, 6.8 Hz, 1H), 2.54 (dt, J = 13.7, 6.6 Hz, 1H), 1.55 (d, J = 6.9 Hz, 3H); **$^{13}$C{$^1$H} NMR** (126 MHz, methanol-*$d_4$*) δ 180.1, 158.8, 153.4, 148.4, 144.8, 136.7, 108.7, 105.4, 104.7, 64.9, 57.7, 46.7, 39.9, 31.5, 23.1; **MS** (ESI) *m/z* calcd. for $C_{15}H_{19}N_3O_8S^+$ [MH]+: 402.10, found: 402.26

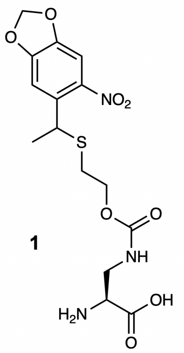

**1**

Chemical Formula: $C_{15}H_{19}N_3O_8S$
Exact Mass: 401.0893

##### ***N*-photo-masked-L-isoserine (pm-isoS, 2)**

(*Following a procedure adapted from Huguenin-Dezot et al., 2019, for a related compound*[2]*)*

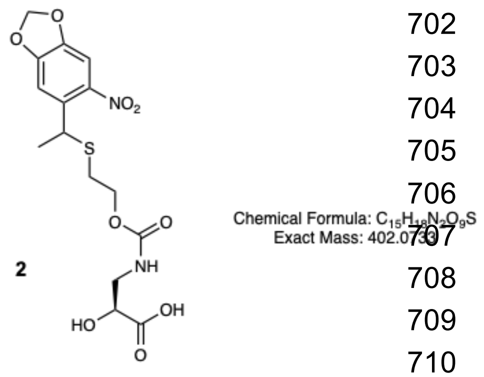

**2**

L-isoserine (0.229 g, 2.16 mmol, 1.08 equiv) was added in one portion to a solution of **16** (1.99 mmol, 1.00 equiv, prepared as described above) in dry $CH_3CN$ (12 mL) under nitrogen gas and stirred for 16 h at 60 °C. After this time the reaction was judged to be complete by LC-MS and the contents dried under reduced pressure. This was then subjected to preparative reverse phase high-performance liquid chromatography (RP-HPLC) on a Waters Prep 150 LC System [CSH $C_{18}$ 19 x 150 mm OBD Column 5 µm; gradient: 20-50% $H_2O$-$CH_3CN$ (+0.1% TFA throughout) mobile phase over 16 min] to obtain the desired photo-masked isoserine (pm-isoS, **2**) as a yellow solid (0.793 g, >99%) and a mixture of ~1:1 epimers. **$^1$H NMR** (500 MHz, methanol-*$d_4$*) δ 7.33 (s, 1H), 7.29 (s, 1H), 6.13 (dd, J = 6.9, 1.1 Hz, 2H), 4.75 (q, J = 6.9 Hz, 1H), 4.20 (dd, J = 6.9, 4.3 Hz, 1H), 4.11 – 3.95 (m, 2H), 3.50 (dt, J = 13.8, 4.2 Hz, 1H), 3.32 (d, J = 8.0 Hz, 1H), 2.58 (dp, J = 27.5, 6.8 Hz, 2H), 1.54 (d, J = 7.0 Hz, 3H); **$^{13}$C{$^1$H} NMR** (126 MHz, methanol-*$d_4$*) δ 175.7, 158.6, 153.4, 148.4, 144.8, 136.7, 108.7, 105.4, 104.7, 70.9, 65.1, 45.5, 39.9, 31.4, 23.1; **MS** (ESI) *m/z* calcd. for $C_{15}H_{18}N_2O_9S$- [M]-: 401.06, found: 401.39.

Note: pm-isoS (**2**) was found to degrade within 2 weeks when kept at -20 °C and was therefore freshly synthesized for each expression.

***N*-photo-masked-DL-isocysteine (pm-isoC, (±)-3)**

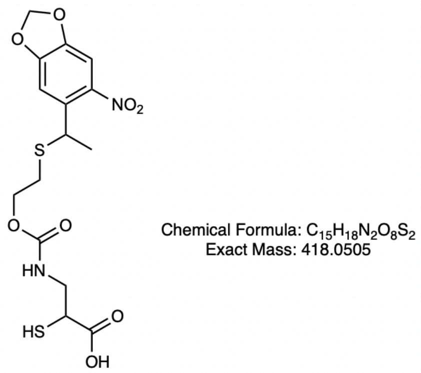

**3**

Thioketal **(±)-20** (530 mg, 1.1 mmol, 1 equiv) was dissolved in 2 mL THF and 1 M LiOH (2 mL) was added dropwise at 0 °C. The mixture was warmed to room temperature and stirred vigorously for 3 h with aluminum foil wrapped around the vial. The reaction was judged to be complete by LC-MS analysis. The reaction was then acidified to pH 5 with 1 N aq. HCl, extracted with EtOAc (3 x 5 mL), and the organic phase was dried over sodium sulfate and concentrated under reduced pressure. The residue was then subjected to preparative reverse phase HPLC on a Waters Prep 150 LC System [CSH C18 19 x 150 mm OBD Column 5 µm; gradient: 5-95% $H_2O$-$CH_3CN$ (+0.1% TFA throughout) mobile phase over 16 min] to obtain the desired photo-masked isocysteine (pm-isoC, **(±)-3**) as a yellow gum (0.456 g, >99%) and a mixture of ~1:1 epimers

$^{1}$**H NMR** (400 MHz, methanol-$d_4$) δ 7.34 (s, 1H), 7.30 (d, J = 0.8 Hz, 1H), 6.17 – 6.11 (m, 2H), 4.75 (q, J = 7.0 Hz, 1H), 4.11 – 3.97 (m, 2H), 3.53 – 3.31 (m, 3H), 2.66 – 2.47 (m, 2H), 1.54 (d, J = 6.9 Hz, 3H); $^{13}$**C{$^{1}$H} NMR** (126 MHz, methanol-$d_4$) δ 158.3, 153.4, 148.4, 144.7, 136.7, 127.0, 108.7, 105.4, 104.7, 64.8, 62.3, 39.9, 34.8, 31.4, 23.1; **MS** (ESI) *m/z* calcd. for $C_{15}H_{18}N_2O_8S_2^-$ [M]-: 417.04, found: 417.34

### NMR and LC-MS analysis of model BEAR reactions

#### Methods to support Figure 3

**LC-MS experiments**

The ester- and thioester-containing model peptides **4** and **(±)-5** were each dissolved in 10% $D_2O$, 90% Protein Buffer (50 mM sodium phosphate, 1.5 mM DTT, pH 8.6) to 3 mM. The thioester linkage within peptide **(±)-5** was unstable under these conditions. We incubated the solution at RT and 37 °C and monitored the hydrolysis and resulting oxidation by LC-HRMS (Agilent) at 0, 1, 3, 5, and 20 h. Hydrolysis of the thioester bond was complete within 3 h, and the resulting free thiol **(±)-8** oxidized to form disulfide **9** within 5 h. The ester-containing peptide **4** was stable under these conditions, and was subjected to photolysis to reveal the $\beta^2$-amine side chain. The solution of **4** described above was exposed to 370 nm light for 2 min at a distance of 0.5 meters, followed by incubation at 37 °C, with the reaction progress monitored via LC-MS (Waters) before UV exposure, and after 3, 6, and 10 h of incubation post-photolysis. The peak due to **4** disappeared within 3 h, and a peak whose mass corresponded to the fully deprotected peptide was observed. In addition, we analyzed a solution of the authentic rearranged standard (**6**) by the same LC-MS method, and found this standard to have the same mass and retention time as the peak found in our experimental samples.

**NMR experiments**

NMR experiments were necessary to confirm that the mass observed corresponded to the expected BEAR product. The ester-containing peptide **4** was dissolved in 10% $D_2O$, 90% Protein Buffer to 3 mM and evaluated using both one-dimensional $^{1}$H NMR (with solvent suppression) as well as multiplicity-edited Heteronuclear Single Quantum Coherence (HSQC) spectroscopy. All NMR spectra were collected using a Bruker 600 MHz NMR spectrometer. Briefly, the solution of **4** was irradiated at 370 nm for 2 min at a distance of 0.5 meters, incubated at 37 °C, and $^{1}$H NMR spectra were taken before irradiation, immediately following irradiation, and after 3 and 16 h of incubation. Many peaks shifted immediately following UV exposure, as expected for the multi-step pathway associated with full side-chain deprotection. We focused primarily on the peak corresponding to $H_\alpha$ of isoserine, which we expected to shift most dramatically following rearrangement (**Supplementary Fig. 2A**). In ester peptide **4**, this $H_\alpha$ peak appears at

5.2 ppm; it begins to disappear immediately after UV irradiation, is significantly reduced after 3 h, and is completely gone after 16 h. Over the same time period, a new peak at 4.3 ppm appeared in the $^{1}H$ NMR spectrum. This peak corresponds to $H_{\alpha}$ in authentic standard **6** which contains a $\beta^{2}$-peptide linkage (**Supplementary Fig. 2B**). Multiplicity-edited HSQC spectroscopy clearly shows the movement of this cross-peak correlation (**Figure 3D**). By 16 hours, the full deprotection and rearrangement to generate **6** is observed. Of note, a slight (0.03 ppm) shift is observed between the authentic standard **6** and the peak observed in the experimental sample. We attribute this minor difference in the experimental sample to 1:1 association formation between the rearranged product (**6**) and a compound arising from the removed protecting group (likely 1-(6-nitrosobenzo[*d*][1,3]dioxol-5-yl)ethan-1-one or a downstream product thereof) as there is no conceivable alteration to the molecule that would result in such a minor ppm difference in the buffered solution.

### General Biological Methods

**Bacterial Strains**

Electrocompetent MegaX DH10B™ T1R Electrocomp™ *E. coli* cells (Invitrogen) were purchased from Thermo Fisher Scientific (catalog number: C640003). NEB® 5-alpha Competent *E. coli* (DH5α) were purchased from New England BioLabs (catalog number: C2987H). BL21(DE3) Competent *E. coli* were purchased from New England BioLabs (catalog number: C2527H). C321.ΔA.exp *E. coli* were gifts from George Church (Addgene bacterial strain #49018).

**Amino Acids**

BocK (SKU 349661), α-OH-BocK (SKU AMBH93D58DB6-100MG), and Lys (SKU L5626) were purchased from Sigma-Aldrich.

**Antibiotics**

The following working antibiotic concentrations were used: Tetracycline, 10 µg/mL; Kanamycin 25 µg/mL; Carbenicillin, 100 µg/mL

**Plasmids**

pBK-DAPRS (encoding DAPRS/$tRNA^{Pyl}_{CUA}$) and p15A-sfGFP150TAG (encoding sfGFP N150TAG reporter) were gifted from the Jason Chin lab. pEVOL-*Ma*PylRS (encoding MaPylRS/$tRNA^{Pyl}_{CUA}$) was previously cloned[6]. pET32a-NanoLuc-6xHis (WT), pET32a-NanoLuc(M1-TAG-2V insertion)-6xHis, pET32a-NanoLuc(M1-G-TAG-2V insertion)-6xHis, pET32a-NanoLuc(G159-TAG-V160 insertion)-6xHis, pET32a-NanoLuc(G103-TAG-V104 insertion)-6xHis, pET32a-NanoLuc(T130TAG)-6xHis, and pET32a-NanoLuc(G131TAG)-6xHis were generated as described below. pET15a-NanoLuc(G159-TAG-V160 insertion)-6xHis was generated as described below. pBK-DAPRS-M265S, pBK-DAPRS-A267H, pBK-DAPRS-M309L, and pBK-DAPRS-M265S-A267S-M309L were generated as described below.

**Transformation protocols**

MegaX DH10B T1R Electrocomp™ Cells (ThermoFisher C640003) were transformed in accordance with manufacturer protocols with some modifications as follows. Frozen stocks were thawed on ice. Upon thawing, 100 ng of each relevant plasmid was added. After a 30 min incubation on ice, cells were electroporated using a Bio-Rad Laboratories, Inc. MicroPulser Electroporator (Boulder, CO) on the 'Bacteria 1' preset. 1 mL of Super Optimal broth with Catabolite repression (S.O.C.) was immediately added and cells were recovered at 37 °C shaking for 1 h before plating 100 µL on LB agar plates containing tetracycline and kanamycin. Plates were incubated overnight at 37 °C.

DH5α *E. coli* (NEB, catalog # C2987H) were transformed in accordance with manufacturer protocols with some modifications as follows. Frozen stocks were thawed on ice. Upon thawing, 100 ng of each relevant plasmid was added. After a 30 min incubation on ice, cells were heat-shocked for 30 sec at 42 °C and allowed to recover on ice for 2 min. Following, 350 μL of Super Optimal broth with Catabolite repression (S.O.C.) was immediately added and cells were recovered at 37 °C shaking for 1 hour before plating 50 μL on LB agar plates containing appropriate antibiotics. Plates were incubated overnight at 37 °C.

BL21(DE3) *E. coli* (NEB, catalog # C2527H) were transformed in accordance with manufacturer protocols with some modifications as follows. Frozen stocks were thawed on ice. Upon thawing, 100 ng of each relevant plasmid was added. After a 30 min incubation on ice, cells were heat-shocked for 30 sec at 42 °C and allowed to recover on ice for 2 min. Following, 350 μL of Super Optimal broth with Catabolite repression (S.O.C.) was immediately added and cells were recovered at 37 °C shaking for 1 hour before plating 50 μL on LB agar plates containing appropriate antibiotics. Plates were incubated overnight at 37 °C.

C321.ΔA.exp *E. coli* frozen stocks were thawed on ice. Upon thawing, 100 ng of each relevant plasmid was added as well as 20 μL of 5x KCM (500 mM KCl, 150 mM $CaCl_2$, 250 mM $MgCl_2$) and 180 μL nuclease-free water. After a 30 min incubation on ice, cells were heat-shocked for 90 sec at 42 °C and allowed to recover on ice for 2 min. Following, 800 μL of Super Optimal broth with Catabolite repression (S.O.C.) was immediately added and cells were recovered at 37 °C shaking for 1 hour before plating 100 μL on LB agar plates containing appropriate antibiotics. Plates were incubated overnight at 37 °C.

**Protein LC-MS Methods**

LC-MS analysis of all protein samples were performed on an Agilent 1290 Infinity II HPLC connected to an Agilent 6530B QTOF AJS-ESI. The mobile phase for LC-MS was water and acetonitrile with 0.1% (v/v) formic acid at a flow rate of 0.4 mL/min. Each protein sample was injected onto an Poroshell 300SB-C8 column (2.1 x 75 mm, 5 μM, room temp, Agilent) and separated using a linear gradient from 5% acetonitrile for 0 to 2 min and ramping to 95% acetonitrile over 7.5 min, and then washing with 95% acetonitrile for 2 min. The following parameters were used during acquisition: Fragmentor voltage 225 V, gas temperature 300 ºC, drying gas flow 10 L/min, sheath gas temperature 350 ºC, sheath gas flow 11 L/min, nebulizer pressure 35 psi, skimmer voltage 65 V, Vcap 5000 V, 1 spectra/s.

### Methods to support Figure 4B

**Transformations** pBK-DAPRS and p15A-sfGFP150TAG were transformed into MegaX DH10B T1R Electrocomp™ Cells using the above protocol.

**Expression assays of 150TAG sfGFP**
Starter *E. coli* cultures were grown overnight in 10 mL of LB Miller (AmericanBio, Catalog #AB01201) in 15 mL culture tubes supplemented with antibiotics at 37 °C. Prior to cultures reaching OD, the appropriate amount of each monomer (stored as 200 mM stocks) were allotted into the wells of a black, clear bottom, 96-well plate (1 mM final concentration unless otherwise indicated). Once cultures reached an $OD_{600}$ of 0.6 (roughly 18 h), protein expression was induced by addition of 0.2% arabinose, and the culture was transferred into the wells of the plate to ensure a total volume of 200 µL in each well. A Breathe-Easy® sealing membrane (Sigma Z380059) was placed over the 96-well plate and the plate was loaded into a BioTek Synergy HTX microplate reader with no lid. $OD_{600}$ and $F_{528}$ values ($\lambda_{ex}$ = 485 nm) were measured every 10 min for 24 h. The plate was maintained at 37 °C and was shaken during this time (**Supplementary Fig. 4**). Points represented in Figure 4 are biological replicates where three random colonies were picked on a plate from single transformation.

**Expression and Purification of N150TAG sfGFP**
The expression and purification of N150pcDAP sfGFP followed prior established protocols[2], and the purified protein was used to optimize photo-demasking conditions.

**Photo-mask deprotection optimization**
Purified sfGFP containing pm-DAP **1** at residue 150 was dissolved in Reaction Buffer (50 mM sodium phosphate buffer, 1.5 mM DTT) at 0.1 mg/mL and distributed among 4 samples in glass LCMS vials. Two samples were at a pH of 6.9 and two were at a pH of 8.6. Each sample was then irradiated at 370 nm using a Kessil PR160L-370-G2 lamp set to 25% maximum wattage (11 W) for 2 min at a distance of 0.5 meters from the lamp. Following irradiation, samples at pH 6.9 or 8.6 were incubated at either RT or 37 °C (total of 4 different conditions). LC-MS time points were taken prior to deprotection and at time = 0 h, 1 h, 2 h, and 6 h. We find that the fastest deprotection condition occurs at pH 8.6 at 37 °C with the majority of the deprotection completed by 2 h (**Supplementary Fig. 5**).

### Methods to support Figure 4C

**Design of NanoLuc variants**

All plasmids were designed to encode proteins bearing a C-terminal 6xHis tag to enable affinity purification. The sequences of the encoded proteins are as follows, with * identifying the location encoded by the TAG codon. The name of the final NanoLuc expression plasmid is given in parentheses.

NanoLuc-6xHis (WT) (pET32a-NanoLuc-6xHis (WT))
MVFTLEDFVGDWRQTAGYNLDQVLEQGGVSSLFQNLGVSVTPIQRIVLSGENGLKIDIH
VIIPYEGLSGDQMGQIEKIFKVVYPVDDHHFKVILHYGTLVIDGVTPNMIDYFGRPYEGIA
VFDGKKITVTGTLWNGNKIIDERLINPDGSLLFRVTINGVTGWRLCERILA**HHHHHH***

NanoLuc(M1-TAG-2V insertion)-6xHis (pET32a-NanoLuc(M1-TAG-2V insertion)-6xHis)
M*VFTLEDFVGDWRQTAGYNLDQVLEQGGVSSLFQNLGVSVTPIQRIVLSGENGLKIDI
HVIIPYEGLSGDQMGQIEKIFKVVYPVDDHHFKVILHYGTLVIDGVTPNMIDYFGRPYEGI
AVFDGKKITVTGTLWNGNKIIDERLINPDGSLLFRVTINGVTGWRLCERILA**HHHHHH***

NanoLuc(M1-G-TAG-2V insertion)-6xHis (pET32a-NanoLuc(M1-G-TAG-2V insertion)-6xHis)
M**G***VFTLEDFVGDWRQTAGYNLDQVLEQGGVSSLFQNLGVSVTPIQRIVLSGENGLKID
IHVIIPYEGLSGDQMGQIEKIFKVVYPVDDHHFKVILHYGTLVIDGVTPNMIDYFGRPYEG
IAVFDGKKITVTGTLWNGNKIIDERLINPDGSLLFRVTINGVTGWRLCERILA**HHHHHH***

NanoLuc(G159-TAG-V160 insertion)-6xHis (pET32a-NanoLuc(G159-TAG-V160 insertion)-6xHis and pET15a-NanoLuc(G159-TAG-V160 insertion)-6xHis)
MVFTLEDFVGDWRQTAGYNLDQVLEQGGVSSLFQNLGVSVTPIQRIVLSGENGLKIDIH
VIIPYEGLSGDQMGQIEKIFKVVYPVDDHHFKVILHYGTLVIDGVTPNMIDYFGRPYEGIA
VFDGKKITVTGTLWNGNKIIDERLINPDGSLLFRVTING*VTGWRLCERILA**HHHHHH***

NanoLuc(G103-TAG-V104 insertion)-6xHis (pET32a-NanoLuc(G103-TAG-V104 insertion)-6xHis)
MVFTLEDFVGDWRQTAGYNLDQVLEQGGVSSLFQNLGVSVTPIQRIVLSGENGLKIDIH
VIIPYEGLSGDQMGQIEKIFKVVYPVDDHHFKVILHYGTLVIDG*VTPNMIDYFGRPYEGI
AVFDGKKITVTGTLWNGNKIIDERLINPDGSLLFRVTINGVTGWRLCERILA**HHHHHH***

NanoLuc(T130TAG)-6xHis (pET32a-NanoLuc(T130TAG)-6xHis)
MVFTLEDFVGDWRQTAGYNLDQVLEQGGVSSLFQNLGVSVTPIQRIVLSGENGLKIDIH
VIIPYEGLSGDQMGQIEKIFKVVYPVDDHHFKVILHYGTLVIDGVTPNMIDYFGRPYEGIA
VFDGKKITV*GTLWNGNKIIDERLINPDGSLLFRVTINGVTGWRLCERILA**HHHHHH***

NanoLuc(G131TAG)-6xHis (pET32a-NanoLuc(G131TAG)-6xHis)
MVFTLEDFVGDWRQTAGYNLDQVLEQGGVSSLFQNLGVSVTPIQRIVLSGENGLKIDIH
VIIPYEGLSGDQMGQIEKIFKVVYPVDDHHFKVILHYGTLVIDGVTPNMIDYFGRPYEGIA
VFDGKKITVT*TLWNGNKIIDERLINPDGSLLFRVTINGVTGWRLCERILA**HHHHHH***

**Cloning of pET32a-NanoLuc expression plasmids (see Supplementary Fig. 6)**

The sequences encoding NanoLuc-6xHis (WT), NanoLuc(M1-TAG-2V insertion)-6xHis, NanoLuc(M1-G-TAG-2V insertion)-6xHis, NanoLuc(G159-TAG-V160 insertion)-6xHis, NanoLuc(G103-TAG-V104 insertion)-6xHis, NanoLuc(T130TAG)-6xHis, and NanoLuc(G131TAG)-6xHis were cloned into pET-32a(+) as follows. Circular pET-32a(+) (1 µg, Millipore Sigma, catalog # 69015-3) was incubated with 1 µL each of restrictions enzymes XbaI (New England Biosciences, catalog # R0145S) and StyI-HF (New England Biosciences, catalog # R3189S) in cutSmart Buffer (New England Biolabs, catalog #R3500S) at 37 °C for 1 h. The entire restriction digest reaction was subject to a PCR clean up with the QIAquick PCR Purification Kit (Qiagen, catalog # 28104). The concentration of purified, linearized pET-32(a)+ was determined by measuring the absorbance at 260 nm using a NanoDrop ND-1000 Spectrophotometer. Next, 33.3 ng of purified, linearized pET-32(a)+ and 100 ng of the gBlock DNA (Integrated DNA Technologies, Coralville, IA) encoding each desired NanoLuc sequence were combined in a 10 µL Gibson Assembly reaction containing HiFI DNA Assembly Master Mix (NEB, catalog #E2621L) and incubated at 50 °C for 1 h to generate circular pET-32(a)+ vectors encoding the desired NanoLuc sequence.

Circularized plasmids from the previous step were transformed into NEB 5-alpha competent *E. coli* (NEB, catalog # C2987H) as follows. Frozen stocks of cells were thawed on ice for 10 min. Upon thawing, the entire Gibson Assembly reaction was added to cells and incubated on ice for 30 minutes. Cells incubated with plasmid were then subjected to heat shock at 42 °C for 30 seconds and placed on ice for 2 minutes. 350 µL of Super Optimal broth with Catabolite repression (S.O.C.) outgrowth medium (NEB, catalog # B9020S) was added to cells and cells were incubated at 37 °C for 1 hour with shaking at 220 rpm. Agar plates containing carbenicillin were inoculated with 50 µL of transformed cells and grown overnight at 37 °C. 3 single colonies per construct were picked and inoculated into liquid cultures containing 10 mL LB + carbenicillin and grown for 16 hours at 37 °C. Pure plasmid was isolated from 10 mL cultures using Qiaprep Spin Miniprep Kit (Qiagen, catalog # 27106) and sequences were confirmed by Sanger sequencing at the UC Berkeley DNA Sequencing Facility and whole plasmid sequencing with Primordium Labs. Plasmids containing each NanoLuc construct were double transformed with pEVOL-MaPylRS into chemically competent BL21 *E. coli* following transformation protocol detailed in General Methods for expression assays.

*NanoLuc-6xHis (WT) gBlock Sequence*
TGTGAGCGGATAACAATTCCCCTCTAGAAATAATTTTGTTTAACTTTAAGAAGGAGA
TATACAatggtgtttaccctggaagattttgtgggcgattggcgccagaccgcgggctataacctggatcaggtgctggaacagggcg
gcgtgagcagcctgtttcagaacctgggcgtgagcgtgaccccgattcagcgcattgtgctgagcggcgaaaacggcctgaaaattgatat
tcatgtgattattccgtatgaaggcctgagcggcgatcagatgggccagattgaaaaaatttttaaagtggtgtatccggtggatgatcatcatt
ttaaagtgattctgcattatggcaccctggtgattgatggcgtgaccccgaacatgattgattattttggccgcccgtatgaaggcattgcggtg
tttgatggcaaaaaaattaccgtgaccggcaccctgtggaacggcaacaaaattattgatgaacgcctgattaacccggatggcagcctgct
gtttcgcgtgaccattaacggcgtgaccggctggcgcctgtgcgaacgcattctggcgcatcaccatcaccatcactaaGGCCGCAC
TCGAGCACCACCACCACCACCACTGAGATCCGGCTGCTAACAAAGCCCGAAAGGAA
GCTGAGTTGGCTGCTGCCACCGCTGAGCAATAACTAGCATAACCCCTTGGGGCCTCT
AAACGGGT

*NanoLuc(M1-TAG-2V insertion)-6xHis gBlock Sequence*
TGTGAGCGGATAACAATTCCCCTCTAGAAATAATTTTGTTTAACTTTAAGAAGGAGA
TATACAatgtaggtgtttaccctggaagattttgtgggcgattggcgccagaccgcgggctataacctggatcaggtgctggaacagg
gcggcgtgagcagcctgtttcagaacctgggcgtgagcgtgaccccgattcagcgcattgtgctgagcggcgaaaacggcctgaaaattg
atattcatgtgattattccgtatgaaggcctgagcggcgatcagatgggccagattgaaaaaatttttaaagtggtgtatccggtggatgatcat
cattttaaagtgattctgcattatggcaccctggtgattgatggcgtgaccccgaacatgattgattattttggccgcccgtatgaaggcattgc
ggtgtttgatggcaaaaaaattaccgtgaccggcaccctgtggaacggcaacaaaattattgatgaacgcctgattaacccggatggcagc
ctgctgtttcgcgtgaccattaacggcgtgaccggctggcgcctgtgcgaacgcattctggcgcatcaccatcaccatcactaaGGCCG
CACTCGAGCACCACCACCACCACCACTGAGATCCGGCTGCTAACAAAGCCCGAAAG
GAAGCTGAGTTGGCTGCTGCCACCGCTGAGCAATAACTAGCATAACCCCTTGGGGC
CTCTAAACGGGT

*NanoLuc(M1-G-TAG-2V insertion)-6xHis gBlock Sequence*
TGTGAGCGGATAACAATTCCCCTCTAGAAATAATTTTGTTTAACTTTAAGAAGGAGA
TATACAatgGGCtaggtgtttaccctggaagattttgtgggcgattggcgccagaccgcgggctataacctggatcaggtgctggaa
cagggcggcgtgagcagcctgtttcagaacctgggcgtgagcgtgaccccgattcagcgcattgtgctgagcggcgaaaacggcctgaa
aattgatattcatgtgattattccgtatgaaggcctgagcggcgatcagatgggccagattgaaaaaatttttaaagtggtgtatccggtggatg
atcatcattttaaagtgattctgcattatggcaccctggtgattgatggcgtgaccccgaacatgattgattattttggccgcccgtatgaaggca
ttgcggtgtttgatggcaaaaaaattaccgtgaccggcaccctgtggaacggcaacaaaattattgatgaacgcctgattaacccggatggc
agcctgctgtttcgcgtgaccattaacggcgtgaccggctggcgcctgtgcgaacgcattctggcgcatcaccatcaccatcactaaGGC
CGCACTCGAGCACCACCACCACCACCACTGAGATCCGGCTGCTAACAAAGCCCGAA
AGGAAGCTGAGTTGGCTGCTGCCACCGCTGAGCAATAACTAGCATAACCCCTTGGG
GCCTCTAAACGGGT

*NanoLuc(G159-TAG-V160 insertion)-6xHis gBlock Sequence*
TGTGAGCGGATAACAATTCCCCTCTAGAAATAATTTTGTTTAACTTTAAGAAGGAGA
TATACAatggtgtttaccctggaagattttgtgggcgattggcgccagaccgcgggctataacctggatcaggtgctggaacagggcg
gcgtgagcagcctgtttcagaacctgggcgtgagcgtgaccccgattcagcgcattgtgctgagcggcgaaaacggcctgaaaattgatat

tcatgtgattattccgtatgaaggcctgagcggcgatcagatgggccagattgaaaaaatttttaaagtggtgtatccggtggatgatcatcattttaaagtgattctgcattatggcaccctggtgattgatggcgtgaccccgaacatgattgattattttggccgcccgtatgaaggcattgcggtgtttgatggcaaaaaaattaccgtgaccggcaccctgtggaacggcaacaaaattattgatgaacgcctgattaacccggatggcagcctgctgtttcgcgtgaccattaacggctaggtgaccggctggcgcctgtgcgaacgcattctggcgcatcaccatcaccatcactaaGGCCGCACTCGAGCACCACCACCACCACCACTGAGATCCGGCTGCTAACAAAGCCCGAAAGGAAGCTGAGTTGGCTGCTGCCACCGCTGAGCAATAACTAGCATAACCCCTTGGGGCCTCTAAACGGGT

*NanoLuc(G103-TAG-V104 insertion)-6xHis gBlock Sequence*

TGTGAGCGGATAACAATTCCCCTCTAGAAATAATTTTGTTTAACTTTAAGAAGGAGATATACAatggtgtttaccctggaagattttgtgggcgattggcgccagaccgcgggctataacctggatcaggtgctggaacagggcggcgtgagcagcctgtttcagaacctgggcgtgagcgtgaccccgattcagcgcattgtgctgagcggcgaaaacggcctgaaaattgatattcatgtgattattccgtatgaaggcctgagcggcgatcagatgggccagattgaaaaaatttttaaagtggtgtatccggtggatgatcatcattttaaagtgattctgcattatggcaccctggtgattgatggctaggtgaccccgaacatgattgattattttggccgcccgtatgaaggcattgcggtgtttgatggcaaaaaaattaccgtgaccggcaccctgtggaacggcaacaaaattattgatgaacgcctgattaacccggatggcagcctgctgtttcgcgtgaccattaacggcgtgaccggctggcgcctgtgcgaacgcattctggcgcatcaccatcaccatcactaaGGCCGCACTCGAGCACCACCACCACCACCACTGAGATCCGGCTGCTAACAAAGCCCGAAAGGAAGCTGAGTTGGCTGCTGCCACCGCTGAGCAATAACTAGCATAACCCCTTGGGGCCTCTAAACGGGT

*NanoLuc(T130TAG)-6xHis gBlock Sequence*

TGTGAGCGGATAACAATTCCCCTCTAGAAATAATTTTGTTTAACTTTAAGAAGGAGATATACAatggtgtttaccctggaagattttgtgggcgattggcgccagaccgcgggctataacctggatcaggtgctggaacagggcggcgtgagcagcctgtttcagaacctgggcgtgagcgtgaccccgattcagcgcattgtgctgagcggcgaaaacggcctgaaaattgatattcatgtgattattccgtatgaaggcctgagcggcgatcagatgggccagattgaaaaaatttttaaagtggtgtatccggtggatgatcatcattttaaagtgattctgcattatggcaccctggtgattgatggcgtgaccccgaacatgattgattattttggccgcccgtatgaaggcattgcggtgtttgatggcaaaaaaattaccgtgTAGggcaccctgtggaacggcaacaaaattattgatgaacgcctgattaacccggatggcagcctgctgtttcgcgtgaccattaacggcgtgaccggctggcgcctgtgcgaacgcattctggcgcatcaccatcaccatcactaaGGCCGCACTCGAGCACCACCACCACCACCACTGAGATCCGGCTGCTAACAAAGCCCGAAAGGAAGCTGAGTTGGCTGCTGCCACCGCTGAGCAATAACTAGCATAACCCCTTGGGGCCTCTAAACGGGT

*NanoLuc(G131TAG)-6xHis gBlock Sequence*

TGTGAGCGGATAACAATTCCCCTCTAGAAATAATTTTGTTTAACTTTAAGAAGGAGATATACAatggtgtttaccctggaagattttgtgggcgattggcgccagaccgcgggctataacctggatcaggtgctggaacagggcggcgtgagcagcctgtttcagaacctgggcgtgagcgtgaccccgattcagcgcattgtgctgagcggcgaaaacggcctgaaaattgatattcatgtgattattccgtatgaaggcctgagcggcgatcagatgggccagattgaaaaaatttttaaagtggtgtatccggtggatgatcatcattttaaagtgattctgcattatggcaccctggtgattgatggcgtgaccccgaacatgattgattattttggccgcccgtatgaaggcattgcggtgtttgatggcaaaaaaattaccgtgaccTAGaccctgtggaacggcaacaaaattattgatgaacgcctgattaacccggatggcagcctgctgtttcgcgtgaccattaacggcgtgaccggctggcgcctgtgcgaacgcattctggcgcatcaccatcaccatcactaaGGCCGCA

CTCGAGCACCACCACCACCACCACTGAGATCCGGCTGCTAACAAAGCCCGAAAGGA
AGCTGAGTTGGCTGCTGCCACCGCTGAGCAATAACTAGCATAACCCCTTGGGGCCTC
TAAACGGGT

**Cloning of pET15a-NanoLuc(G159-TAG-V160 insertion)-6xHis**

The sequence encoding NanoLuc(G159-TAG-V160 insertion)-6xHis were cloned into a pET15a vector gifted from the Chin lab as follows. Circular p15A-sfGFP150TAG vector was linearized using oligonucleotides LTR1 and LTR2 (**Supplementary Table 1**). NanoLuc(G159-TAG-V160 insertion)-6xHis was PCR amplified out of pET32a-NanoLuc(G159-TAG-V160 insertion)-6xHis using oligonucleotides LTR3 and LTR4 (**Supplementary Table 1**). PCR using the Q5® High-Fidelity 2X Master Mix (NEB M0492S) was performed according to manufacturer protocols for 30 cycles. The PCR reaction was subject to a PCR clean up with the QIAquick PCR Purification Kit (Qiagen, catalog # 28104). The concentration of purified, linearized products was determined by absorbance at 260 nm using a NanoDrop ND-1000 Spectrophotometer. Next, 33.3 ng of purified, linearized pET15a and 100 ng of linearized NanoLuc(G159-TAG-V160 insertion)-6xHis were combined in a 10 µL Gibson Assembly reaction containing HiFi DNA Assembly Master Mix (NEB, catalog #E2621L) and incubated at 50 °C for 1 hour to generate a circular pET15a vector containing the coding sequence for NanoLuc(G159-TAG-V160 insertion)-6xHis. The circularized plasmid from the previous step was transformed into NEB 5-alpha competent *E. coli* (NEB, catalog # C2987H) as follows. Frozen stocks of cells were thawed on ice for 10 minutes. Upon thawing the entirety of the previous Gibson Assembly reaction was added to cells and incubated on ice for 30 minutes. Cells incubated with plasmid were then subjected to heat shock at 42 °C for 30 seconds and placed on ice for 2 minutes. 350 µL of Super Optimal broth with Catabolite repression (S.O.C.) outgrowth medium (NEB, catalog # B9020S) was added to cells and cells were incubated at 37 °C for 1 hour with shaking at 220 rpm. Agar plates containing tetracycline were inoculated with 50 µL of transformed cells and grown overnight at 37 °C. 3 single colonies per construct were picked and inoculated into liquid cultures containing 10 mL LB + carbenicillin and grown for 16 hours at 37 °C. Pure plasmid was isolated from 10 mL cultures using Qiaprep Spin Miniprep Kit (Qiagen, catalog # 27106) and sequences were confirmed by Sanger sequencing at the UC Berkeley DNA Sequencing Facility whole plasmid sequencing with Primordium Labs. The new pET15a-NanoLuc(G159-TAG-V160 insertion)-6xHis was double transformed with a pBK-DAPRS into MegaX DH10B T1R Electrocomp™ Cells following transformation protocol detailed in General Methods for expression assays.

**Expression of NanoLuc from pET15a-NanoLuc(G159-TAG-V160 insertion)-6xHis (Figure 4C and Supplementary Fig. 6 and 8)**

Starter cultures of E. coli picked from a single colony were grown overnight in 10 mL of LB Miller (AmericanBio, Catalog #AB01201) in a 15 mL culture tube supplemented with antibiotics at 37 °C. Saturated overnight cultures were used to induce a 50 mL culture (1:10) of each construct and grown in 37 °C with shaking at 220 rpm, and OD was monitored over the next several hours. Prior to cultures reaching the desired OD, appropriate amounts of monomers (stored as 200 mM stocks) were transferred into a black, clear bottom, 96-well plate (1 mM final concentration unless otherwise indicated). Once cultures reached an $OD_{600}$ of 0.6, protein expression was induced by addition of 1 mM IPTG or 0.2% arabinose (for pET32a and pET15a respectively), and culture was transferred into the plate for a total volume of 200 µL in each well. A Breathe-Easy® sealing membrane (Sigma Z380059) was placed over the 96-well plate and placed into a 37 °C shaker while shielding the plate from light. After 16 h, 50 µL samples from each condition were transferred onto a white-walled 96-well plate and the Nano-Glo® Luciferase Assay System (Promega N1110) was added according to manufacturer instructions. The luminescence was measured in BioTek Synergy H1 microplate reader as an endpoint assay. Points represented in Figure 4 are biological replicates where three random colonies were picked on a plate from single transformation. Points represented on Supplementary Fig. 6 and 8 are technical replicates where a random single colony was picked and the same experiment was performed three times.

#### Methods to support Figure 4D

**Large Scale Expression and Purification NanoLuc G159-pm-DAP-V160 and G159-pm-isoS-V160**

Starter cultures of 10 mL of Miller's LB Broth (AmericanBio, catalog # AB01201) supplemented with tetracycline and kanamycin were inoculated with a single colony of *E. coli* MegaX DH10B T1R Electrocomp™ cells harboring pBK-DAPRS and pET15a-NanoLuc(G159-TAG-V160 insertion)-6xHis and grown for 24 h at 37 °C with shaking at 220 rpm until the culture was saturated. The starter culture (10 mL) was used to inoculate a 200 mL expression culture of Miller's LB Broth supplemented with 1 mM pm-DAP **1** or pm-isoS **2** and also tetracycline and kanamycin. The expression culture was grown at 37 °C with shaking at 220 rpm to an OD600 of 0.6 at which point it was induced with 0.2% arabinose and grown for 16 hours under the same conditions. The expression culture was harvested by centrifugation at 4,300 x g at 4 °C for 30 minutes. The resulting cell pellet was suspended in 10 mL of Lysis Buffer (20 mM HEPES, 50 mM KCl, 10% glycerol, 10 mM imidazole pH 8.6) containing 1 tablet of cOmplete, mini EDTA-free ULTRA protease inhibitor cocktail (Sigma-Aldrich, St. Louis, MO). The cell suspension was disrupted by sonication on ice (Branson Sonifier 250, 5 cycles of 30 second pulse at 50% duty cycle and microtip limit of 5 followed by 30 second pause). The cell lysate was cleared by centrifugation at 23,000xg at 4 °C for 20 minutes. TALON® Metal Affinity Resin (2 mL) (Takara Biosciences, catalog # 635504) was equilibrated with Lysis Buffer, added to the cleared cell lysate, and incubated on a rotisserie at 4 °C for 1 hour. The TALON® resin-lysate mixture was then passed through a gravity flow Poly-Prep Chromatography Column (Bio-Rad Laboratories, Hercules, CA). Non-specifically bound proteins were removed by washing the TALON® resin with 10 mL of Lysis Buffer. The 6xHis-tagged protein was eluted by washing the TALON® resin with 2 mL of Elution Buffer (20 mM HEPES, 50 mM KCl, 10% glycerol, 1 M imidazole pH 8.6). The purified protein was loaded on to a PD-10 column (Cytiva Life Sciences, catalog # 17085101) and exchanged into 3.5 mL of Storage Buffer (20 mM HEPES, 50 mM KCl, 10% glycerol, pH 8.6) according to manufacturer's instructions. The purified protein was quantified using absorbance at 280 nm, snap frozen as single-use aliquots, and stored at -80 °C. The typical expression yield of NanoLuc constructs using the above protocol was 0.4 mg/L of *E. coli* culture.

### Methods to support Supplementary Fig. 8 - DAPRS variants

**Design of DAPRS variants**

Expression plasmids encoding four DAPRS variants were prepared. Each one contains one or all three mutations reported to improve activity of PylRS for α-hydroxy acid substrates[7]. The sequences encoded by the four plasmids, as well as the sequence of DAPRS as reported, are as follows:

DAPRS (pBK-DAPRS)
MDKKPLDVLISATGLWMSRTGTLHKIKHHEVSRSKIYIEMACGDHLVVNNSRSCRTAR
AFRHHKYRKTCKRCRVSDEDINNFLTRSTESKNSVKVRVVSAPKVKKAMPKSVSRAPK
PLENSVSAKASTNTSRSVPSPAKSTPNSSVPASAPAPSLTRSQLDRVEALLSPEDKISLNM
AKPFRELEPELVTRRKNDFQRLYTNDREDYLGKLERDITKFFVDRGFLEIKSPILIPAEYV
ERMGINNDTELSKQIFRVDKNLCLRPMLAPTLCNYLRKLDRILPGPIKIFEVGPCYRKESD
GKEHLEEFTMVQFCQMGSGCTRENLEALIKEFLDYLEIDFEIVGDSCMVFGDTLDIMHG
DLELSSACVGPVSLDREWGIDKPWIGAGFGLERLLKVMHGFKNIKRASRSESYYNGIST
NL*

DAPRS-M265S (pBK-DAPRS-M265S)
MDKKPLDVLISATGLWMSRTGTLHKIKHHEVSRSKIYIEMACGDHLVVNNSRSCRTAR
AFRHHKYRKTCKRCRVSDEDINNFLTRSTESKNSVKVRVVSAPKVKKAMPKSVSRAPK
PLENSVSAKASTNTSRSVPSPAKSTPNSSVPASAPAPSLTRSQLDRVEALLSPEDKISLNM
AKPFRELEPELVTRRKNDFQRLYTNDREDYLGKLERDITKFFVDRGFLEIKSPILIPAEYV
ERMGINNDTELSKQIFRVDKNLCLRP**S**LAPTLCNYLRKLDRILPGPIKIFEVGPCYRKESD
GKEHLEEFTMVQFCQMGSGCTRENLEALIKEFLDYLEIDFEIVGDSCMVFGDTLDIMHG
DLELSSACVGPVSLDREWGIDKPWIGAGFGLERLLKVMHGFKNIKRASRSESYYNGIST
NL*

DAPRS-A267H (pBK-DAPRS-A267H)
MDKKPLDVLISATGLWMSRTGTLHKIKHHEVSRSKIYIEMACGDHLVVNNSRSCRTAR
AFRHHKYRKTCKRCRVSDEDINNFLTRSTESKNSVKVRVVSAPKVKKAMPKSVSRAPK
PLENSVSAKASTNTSRSVPSPAKSTPNSSVPASAPAPSLTRSQLDRVEALLSPEDKISLNM
AKPFRELEPELVTRRKNDFQRLYTNDREDYLGKLERDITKFFVDRGFLEIKSPILIPAEYV
ERMGINNDTELSKQIFRVDKNLCLRPML**H**PTLCNYLRKLDRILPGPIKIFEVGPCYRKESD
GKEHLEEFTMVQFCQMGSGCTRENLEALIKEFLDYLEIDFEIVGDSCMVFGDTLDIMHG
DLELSSACVGPVSLDREWGIDKPWIGAGFGLERLLKVMHGFKNIKRASRSESYYNGIST
NL*

DAPRS-M309L (pBK-DAPRS-M309L)
MDKKPLDVLISATGLWMSRTGTLHKIKHHEVSRSKIYIEMACGDHLVVNNSRSCRTAR
AFRHHKYRKTCKRCRVSDEDINNFLTRSTESKNSVKVRVVSAPKVKKAMPKSVSRAPK

PLENSVSAKASTNTSRSVPSPAKSTPNSSVPASAPAPSLTRSQLDRVEALLSPEDKISLNM
AKPFRELEPELVTRRKNDFQRLYTNDREDYLGKLERDITKFFVDRGFLEIKSPILIPAEYV
ERMGINNDTELSKQIFRVDKNLCLRPMLAPTLCNYLRKLDRILPGPIKIFEVGPCYRKESD
GKEHLEEFT**L**VQFCQMGSGCTRENLEALIKEFLDYLEIDFEIVGDSCMVFGDTLDIMHGD
LELSSACVGPVSLDREWGIDKPWIGAGFGLERLLKVMHGFKNIKRASRSESYYNGISTNL
*

DAPRS-M265S-A267S-M309L (pBK-DAPRS-M265S-A267S-M309L)
MDKKPLDVLISATGLWMSRTGTLHKIKHHEVSRSKIYIEMACGDHLVVNNSRSCRTAR
AFRHHKYRKTCKRCRVSDEDINNFLTRSTESKNSVKVRVVSAPKVKKAMPKSVSRAPK
PLENSVSAKASTNTSRSVPSPAKSTPNSSVPASAPAPSLTRSQLDRVEALLSPEDKISLNM
AKPFRELEPELVTRRKNDFQRLYTNDREDYLGKLERDITKFFVDRGFLEIKSPILIPAEYV
ERMGINNDTELSKQIFRVDKNLCLRP**S**L**H**PTLCNYLRKLDRILPGPIKIFEVGPCYRKESD
GKEHLEEFT**L**VQFCQMGSGCTRENLEALIKEFLDYLEIDFEIVGDSCMVFGDTLDIMHGD
LELSSACVGPVSLDREWGIDKPWIGAGFGLERLLKVMHGFKNIKRASRSESYYNGISTNL
*

**Construction of plasmids encoding each DAPRS variant into pBK vector**

pBK plasmids encoding DAPRS-M265S, DAPRS-A267H, and DAPRS-M309L were generated by site-directed mutagenesis of pBK-DAPRS[2], a gift from Jason Chin (MRC). Point mutations were introduced using the Q5® Site-Directed Mutagenesis Kit (NEB, catalog #E0554S) according to manufacturer instructions and using the following primers: DAPRS-M265S used oligonucleotides LTR5 and LTR6. DAPRS-A267H used oligonucleotides LTR7 and LTR8. DAPRS-M309L used oligonucleotides LTR9 and LTR10 (**Supplementary Table 1**). A pBK plasmid encoding DAPRS-M265S-A267S-M309L was also generated from pBK-DAPRS. The circular pBK-DAPRS was linearized by PCR using oligonucleotides LTR11 and LTR12 (**Supplementary Table 1**). PCR was performed using the Q5® High-Fidelity 2X Master Mix (NEB M0492S) and according to the manufacturer protocols for 30 cycles. The products of the PCR reaction were purified using a QIAquick PCR Purification Kit (Qiagen, catalog # 28104). The concentration of purified, linearized pBK was determined by absorbance at 260 nm using a NanoDrop ND-1000 Spectrophotometer. Next, 33.3 ng of purified, linearized pBK and 100 ng of a gBlock DNA fragment (Integrated DNA Technologies, Coralville, IA) encoding DAPRS-M265S-A267S-M309L were combined in a 10 µL Gibson Assembly reaction containing HiFI DNA Assembly Master Mix (NEB, catalog #E2621L) and incubated at 50 °C for 1 h to generate a circular pBK vector encoding DAPRS-M265S-A267S-M309L. Circularized plasmids from the previous step were transformed into NEB 5-alpha competent *E. coli* (NEB, catalog # C2987H) as follows. Frozen stocks of cells were thawed on ice for 10 minutes. Upon thawing the entirety of the previous Gibson Assembly reaction was added to cells and incubated on ice for 30 minutes. Cells incubated with plasmid were then subjected to heat shock at 42 °C for 30 seconds and placed on ice for 2 minutes. 350 µL of Super Optimal broth with Catabolite repression (S.O.C.)

outgrowth medium (NEB, catalog # B9020S) was added to cells and cells were incubated at 37 °C for 1 hour with shaking at 220 rpm. Agar plates containing kanamycin were inoculated with 50 µL of transformed cells and grown overnight at 37 °C.

3 single colonies per DAPRS construct were picked and inoculated into liquid cultures containing 10 mL LB + kanamycin and grown for 16 hours at 37 °C. Pure plasmid was isolated from 10 mL cultures using Qiaprep Spin Miniprep Kit (Qiagen, catalog # 27106) and sequences were confirmed by Sanger sequencing at the UC Berkeley DNA Sequencing Facility and whole plasmid sequencing with Primordium Labs. Plasmids containing each DAPRS construct were double transformed with a pET15a-NanoLuc(G159-TAG-V160 insertion)-6xHis into chemically MegaX DH10B T1R Electrocomp™ Cells, BL21(DE3) cells, and C321.ΔA.exp cells following transformation protocol detailed in General Methods for expression assays.

*DAPRS-M265S-A267S-M309L gBlock Sequence*

attatctgggcaaactggaacgtgatatcaccaaatttttgtggatcgcggctttctggaaattaaaagcccgattctgattccggcggaatat
gtggaacgtatgggcattaacaacgacaccgaactgagcaaacaaattttccgcgtggataaaaacctgtgcctgcgtccgAGCctgC
ATccgaccctgtgtaactatctgcgtaaactggatcgtattctgccgggtccgatcaaaatttttgaagtgggcccgtgctatcgcaaagaaa
gcgatggcaaagaacacctggaagaattcaccCTGgttcagttttgccaaatgggcagcggctgcacc

### Peptide Mapping and MS/MS

#### Methods to support Figure 5B-C

A total of nine NanoLuc samples were isolated from DH10B *E. coli* growths containing DAPRS and the monomers indicated below (and in **Fig. 5a**) either before or after photo-unmasking. All were analyzed by high-resolution tryptic mapping.

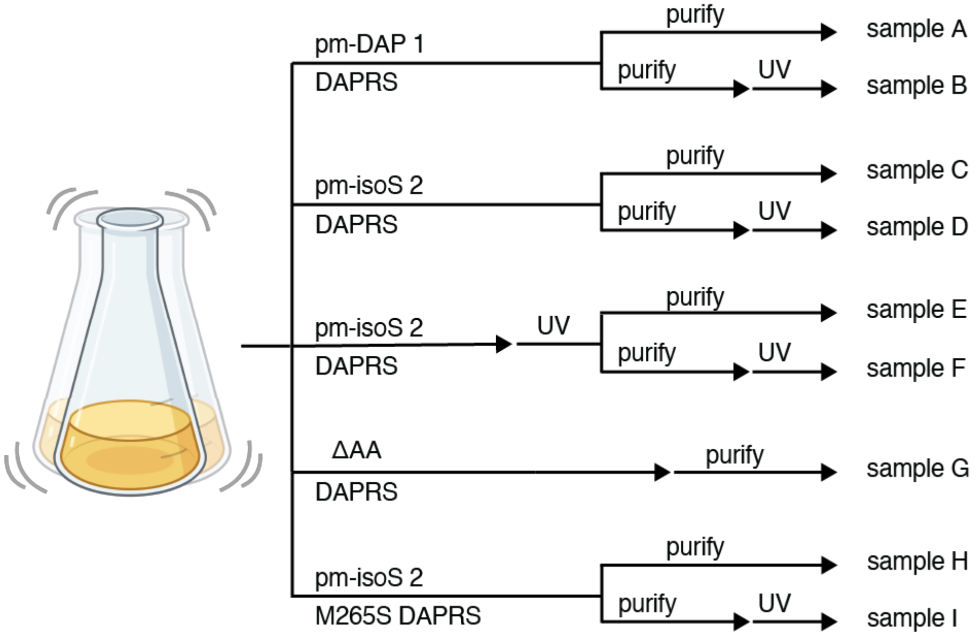

**Trypsin digestion**

The initial concentration of the nine NanoLuc samples varied from 0.1 to 0.8 mg/mL in Storage Buffer (20 mM HEPES, 50 mM KCl, 10% glycerol, pH 8.6). Each sample was concentrated, if necessary, to ~0.8 mg/mL using a microcon 10K filter. 50 µL of each sample (~40 µg) was then added to 150 µL of Denaturing Buffer (8 M guanidine HCl, 0.2 M Tris, pH 7.5) to achieve a final concentration of 6 M guanidine and 0.15 M Tris. Disulfides were reduced by adding 400 mM dithiothreitol (DTT) stock solution in deionized water to achieve a final DTT concentration of 8 mM, followed by incubating at 37 °C for 30 min. The reduced and denatured product was then alkylated by adding 400 mM iodoacetamide to achieve a final concentration of 4 mM, and incubating at 25 °C for 25 min, followed by the addition of DTT to a final concentration of 6 mM to quench the reaction. The reduced/alkylated protein was exchanged into ~45 µL of 0.1 M Tris buffer at pH 7.5 using a Microcon 10-kDa membrane, followed by addition of 4 µg trypsin (in a 1 µg/µL solution) directly to the membrane to achieve an enzyme-to-substrate ratio of at least 1:10. After 2 h at 37 °C, the digestion was quenched with an equal volume of 0.25 M acetate buffer (pH 4.8) containing 6 M guanidine. Peptide fragments were collected by spinning down through the membrane and then subjected to LC-MS/MS analysis.

**Data Collection**
LC-MS/MS analysis was performed on an Agilent 1290-II HPLC directly connected to a Thermo Fisher Q Exactive HF high-resolution mass spectrometer. Peptides were separated on a Waters HSS T3 reverse phase column (2.1 × 150 mm) at 50 °C with a 70-min acetonitrile gradient (0.5% to 35%) containing 0.1% formic acid in the mobile phase, and a total flow rate of 0.25 mL/min. The MS data were collected at 120k resolution setting, followed by data-dependent higher-energy collision dissociation (HCD) MS/MS at a normalized collision energy of 25%.

**Data Analysis**
Proteolytic peptides were identified and quantified on MassAnalyzer, an in-house developed program (available in Biopharma Finder™ from Thermo Fisher). The program performs feature extraction, peptide identification, retention time alignment, and peak integration in an automated fashion. Searched modifications include those shown in **Supplementary Table 2**, as well as amino acid substitutions.

**Selection of peptides for quantitation**
Trypsin digestion of each NanoLuc sample produced a peptide spanning residues 155-165 with an additional residue (X) between the native G159 and V160: VTING(X)VTGWR (**Supplementary Fig. 9A**). Preliminary data analysis revealed that a glutamine (Q) residue was present at site X in all nine samples. Therefore, for data analysis purposes, glutamine was considered as the native residue at site X, and all other residues present at site X were considered as modified forms of glutamine.

****Note that in the data and discussion that follows, V160 in the native protein will be labeled as V161 and X will be in position 160.**

**Quantitation**
It was found that each peptide of interest had one abundant charge state and all other charge states were of much lower intensity. We therefore use the most abundant charge state for quantification. To calculate the relative abundance of each variant in each sample (i.e. which residue is present in place of X), we determined the peak area of each species containing the variant (**Extended Data 2**), and divided it by the total peak area of all the species. Relative percentages showing how the identity of X varied across all NanoLuc Samples (A-I) is available in **Supplementary Fig. 9C**.

#### Methods to support Figure 5D

Three peptide standards were synthesized using solid phase methods and purified to aid in the interpretation of tryptic mapping data and confirm the product of a BEAR cyclization.

**Peptide A**: Val-Thr-Ile-Asn-Gly-**isoS**-Val-Thr-Gly-Trp-Arg

**Peptide B**: Val-Thr-Ile-Asn-Gly-**Ser**-Val-Thr-Gly-Trp-Arg
**Peptide C**: Val-Thr-Ile-Asn-Gly-**Ala**-Val-Thr-Gly-Trp-Arg

**Reagents**. All purchased reagents were used without further purification. Fmoc-Arg-Wang resin and standard fluorenylmethyloxycarbonyl (Fmoc)-protected amino acids were purchased from Novabiochem (San Diego, CA). Trifluoroacetic acid (TFA), piperidine, benzotriazol-1-yloxytripyrrolidinophosphonium hexafluorophosphate (PyBOP), and 1-hydroxybenzotriazole (HOBt) were purchased from Sigma Aldrich (St. Louis, MO). 2-(1H-benzotriazol-1-yl)-1,1,3,3-tetramethyluronium hexafluorophosphate (HBTU) was purchased from Chem-Impex International Inc. (Wood Dale, IL). Dichloromethane (DCM), *N,N*-dimethylformamide (DMF), and 2 M *N,N*-diisopropylethylamine (DIPEA) in *N*-methylpyrrolidone (NMP) were purchased from Fisher Scientific (Hampton, NH). Triisopropylsilane (TIPS) was purchased from Acros Organics (Carlsbad, CA). (2*S*)-2-(*tert*-butoxy)-3-({[(9*H*-fluoren-9-yl)methoxy]carbonyl}amino)propanoic acid (Fmoc-isoS(tBu)-OH) was purchased from Enamine LLC (Monmouth Junction, New Jersey).

**Solid phase peptide synthesis**. Peptides A and B were synthesized on a 50 μmol scale using Fmoc chemistry and Fmoc-Arg(Pbf)-Wang resin (0.35 mmol/g substitution, 100 - 200 mesh) on a PurePep® Chorus from Gyros Protein Technologies AB (Tucson, AZ). Resin was first swollen in 6 mL of DMF for 20 min at RT. The Fmoc protecting group was removed by treating the resin twice (3 min each, at RT) with 5 mL of Deprotection Solution (20% (v/v) piperidine in DMF). Excess Deprotection Solution was removed by washing the resin 4 times with 6 mL DMF. Coupling reactions were performed by adding 5 equiv. amino acid, HOBt, and PyBOP (from 200 mM stocks in DMF) and 10 equiv. DIPEA (from a 2 M stock in NMP) to the deprotected resin for a final resin-bound peptide concentration of 12.5 mM. In general, each amino acid coupling reaction was performed at 70 °C for 5 min. For Peptide A, the coupling reaction that added isoserine (isoS) was modified to contain only 1 equiv. Fmoc-isoS(tBu)-OH. Excess coupling reagents were removed by washing 4 times with 6 mL of DMF. Once the synthesis was complete, the resin-bound peptide was dried under nitrogen and removed from the resin by treatment with 2 mL of Cleavage Cocktail (95% TFA, 2.5% water, 2.5% TIPS) stirred for 1 h at RT. Cleaved peptide was dried under reduced pressure before purification via HPLC. Peptide C was synthesized as described above for Peptides A and B with the following modifications. Coupling reactions were performed by adding 5 equiv. amino acid, HOBt, and HBTU (from 200 mM stocks in DMF) and 10 equiv. of DIPEA (from a 2 M stock in NMP) to the deprotected resin for a final resin-bound peptide concentration of 12.5 mM.

**HPLC purification**. HPLC purification was performed on an Waters LC Prep 150 system equipped with a Waters 2998 UV photodiode array detector, a Waters 2707 autosampler, and a Waters Fraction Collector III using a preparative reverse-phase C18 column (CSH C18 19 x 150 mm OBD Column 5 μm). The mobile phase for HPLC was water with 0.1% (v/v) trifluoroacetic acid (solvent A) and acetonitrile with 0.1% (v/v) trifluoroacetic acid (solvent B). Peptides were eluted at a flow rate of 20 mL/min using a linear solvent gradient from 5 - 50% acetonitrile in water over 30 minutes. Peptides were collected based on their absorbance at 280 nm.

**Mass spectrometry of purified peptides**. LC-MS analysis of each peptide was performed on an Agilent 1290 Infinity II HPLC connected to an Agilent 6530B QTOF AJS-ESI. The mobile phase for LC-MS was water and acetonitrile with 0.1% (v/v) formic acid at a flow rate of 0.7 mL/min. Each peptide was injected onto an Eclipse XDB C18 column (2.1 x 50 mm, 1.8-Micron, room temperature, Agilent) and separated using a linear gradient from 5 to 95% acetonitrile over 4.5 minutes after an initial hold at 5% acetonitrile for 0.5 minutes. The following parameters were used during acquisition: Fragmentor voltage 175 V, gas temperature 300 ºC, gas flow 8 L/min, sheath gas temperature 350 ºC, sheath gas flow 11 L/min, nebulizer pressure 35 psi, skimmer voltage 65 V, Vcap 3500 V, 1 spectra/s.

#### Methods to support Figure 5E-F

Samples D, F, and G were digested with trypsin as described earlier. Synthetic peptide standards were each spiked into the digests. Un-spiked and spiked digests, as well as synthetic peptides, were each analyzed as described in ‘Data Collection’ above.

****Note that in the data and discussion that follows, V160 in the native protein will be labeled as V161 and X will be in position 160.**

***The late-eluting serine isomer has serine at position X***

The intended $\beta^2$-amino acid and its isomeric ester precursor have the same elemental composition as serine and therefore are considered serine isomers. Comparing the elution profiles of serine isomer-containing peptides to VTINGSVTGWR (Peptide B) reveals that the synthetic Ser-containing peptide has a similar retention time as the serine isomer observed in the negative control (Sample G), as well as the late-eluting species observed in Sample D (**Supplementary Fig. 10B**).

To further confirm that the synthetic Peptide B indeed coelutes with the corresponding isomeric species in the Sample G and Sample D, a sample of Peptide B was spiked into the Sample G and Sample D digests, then analyzed using the same method. Observing one single peak in the spiked sample demonstrates the coelution of the synthetic peptide with the isomeric species in the Sample G (**Supplementary Fig. 10C**). The “Sample D + Ser peptide” sample was prepared and analyzed, showing the serine-containing peptide indeed co-elutes with the late-eluting species in Sample D (**Supplementary Fig. 10D**).

**Supplementary Fig. 10E-H** show the MS/MS spectra of different isomeric species. The exact match between the MS/MS spectra of synthetic Peptide B and that of the relevant tryptic peptide from Sample G indicates that they are the same species. In a similar way, the exact match between the MS/MS spectra of synthetic Peptide B and that of the late-eluting species in the tryptic digest from Sample D indicates that they are the same species.

In summary, taking into account both retention time and the equivalence of the MS/MS spectra, we are confident that the isomeric species in Sample G as well the late-eluting species in Sample D have serine at location X corresponding to the sequence of Peptide B.

***The amino acid at location X in the detected “Q160A” species is alanine***

Alanine-containing Peptide C was used to determine that the high abundance of Ala in samples expressed with pm-isoS **2** is indeed due to the incorporation of Ala and not the result of an unexpected side reaction of pm-isoS. A comparison of the elution profiles of alanine-containing Peptide C (VTINGAVTGWR, **Supplementary Fig. 11A**) reveals that the Peptide C coelutes with the detected “Q160A" species, indicating that the “Q160A" species contains alanine at location X. The “Q160A" species also has the same MS/MS spectrum as the synthetic peptide standard (**Supplementary Fig. 11B and 11C**).

### Supplementary Figures

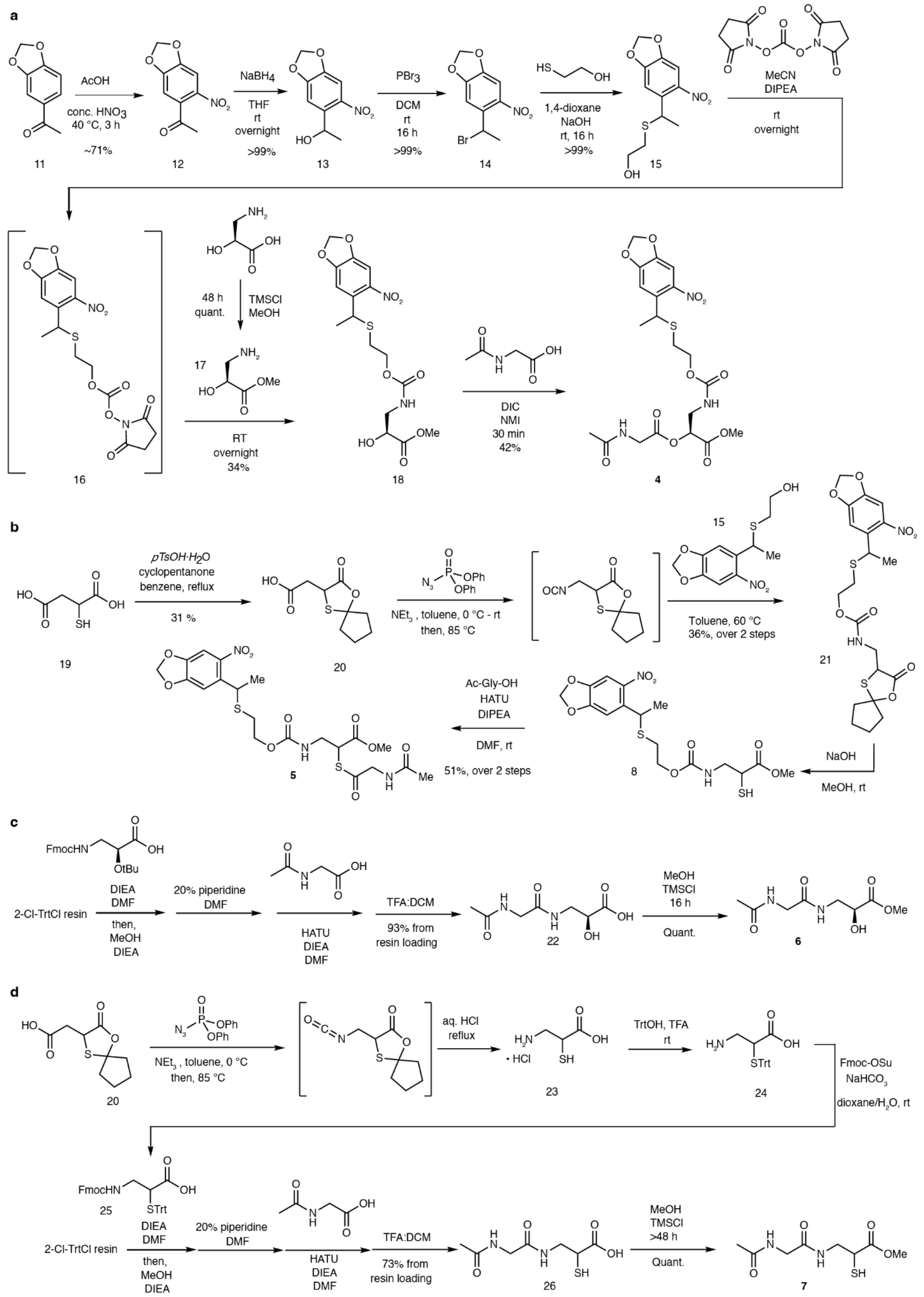

**Supplementary Fig. 1. Synthetic routes to model peptides 4 and (±)-5 and authentic standards 6 and (±)-7.** (A) Solution-phase preparation of the photo-masked *N*-acetylglycyl-isoserine methyl ester (pm-isoS(Ac-Gly)-OMe, **4**). (B) Solution-phase preparation of the photo-masked *N*-acetylglycyl-isocysteine methyl ester (pm-isoC(Ac-Gly)-OMe, **(±)-5**). (C) On-resin preparation of Ac-Gly-isoS-OMe (**6**). (D) Solution-phase preparation of Fmoc-isoC(Trt)-OH and on-resin preparation of Ac-Gly-isoC-OMe (**(±)-7**).

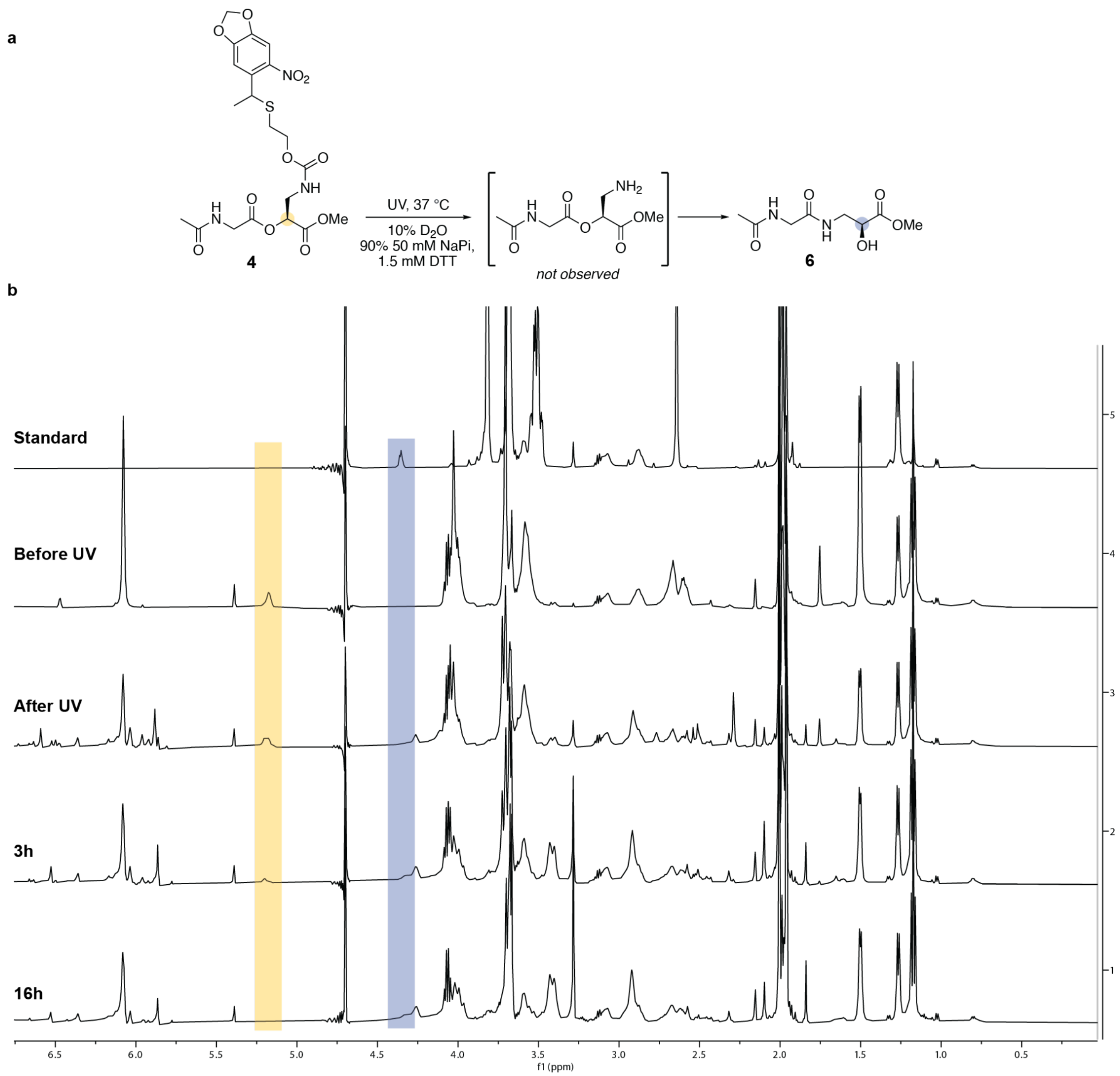

**Supplementary Fig. 2. One-dimensional $^{1}$H NMR spectroscopy provides evidence for BEAR reaction of ester 4 to generate amide 6.** (A) Ester-containing peptide **4** was dissolved in 10% $D_2O$, 90% Reaction Buffer (50 mM Na-phosphate, 1.5 mM DTT, pH 8.6) at 3 mM and subjected to 370 nm irradiation for 2 min, followed by incubation at 37 ºC for the indicated time. NMR spectroscopic analyses focused on the Hα-Cα which is expected to shift most dramatically following rearrangement from an α-ester (yellow) to a $\beta^2$-amide (blue). (B) 1D $^{1}$H NMR spectra are shown comparing the NMR spectra of the authentic standard (**6**) to that of **4** before UV irradiation, immediately after irradiation, and after 3 and 16 h of incubation at 37 ºC following irradiation. The Hα from the substrate ester **4** (yellow) disappears after 3 h while a new peak (blue, left shoulder) appears upfield that matches the authentic standard, confirming a successful BEAR reaction to generate the $\beta^2$-amide. Note that integrations are not reported here because experiments were performed with solvent suppression, resulting non-quantitative integrations.

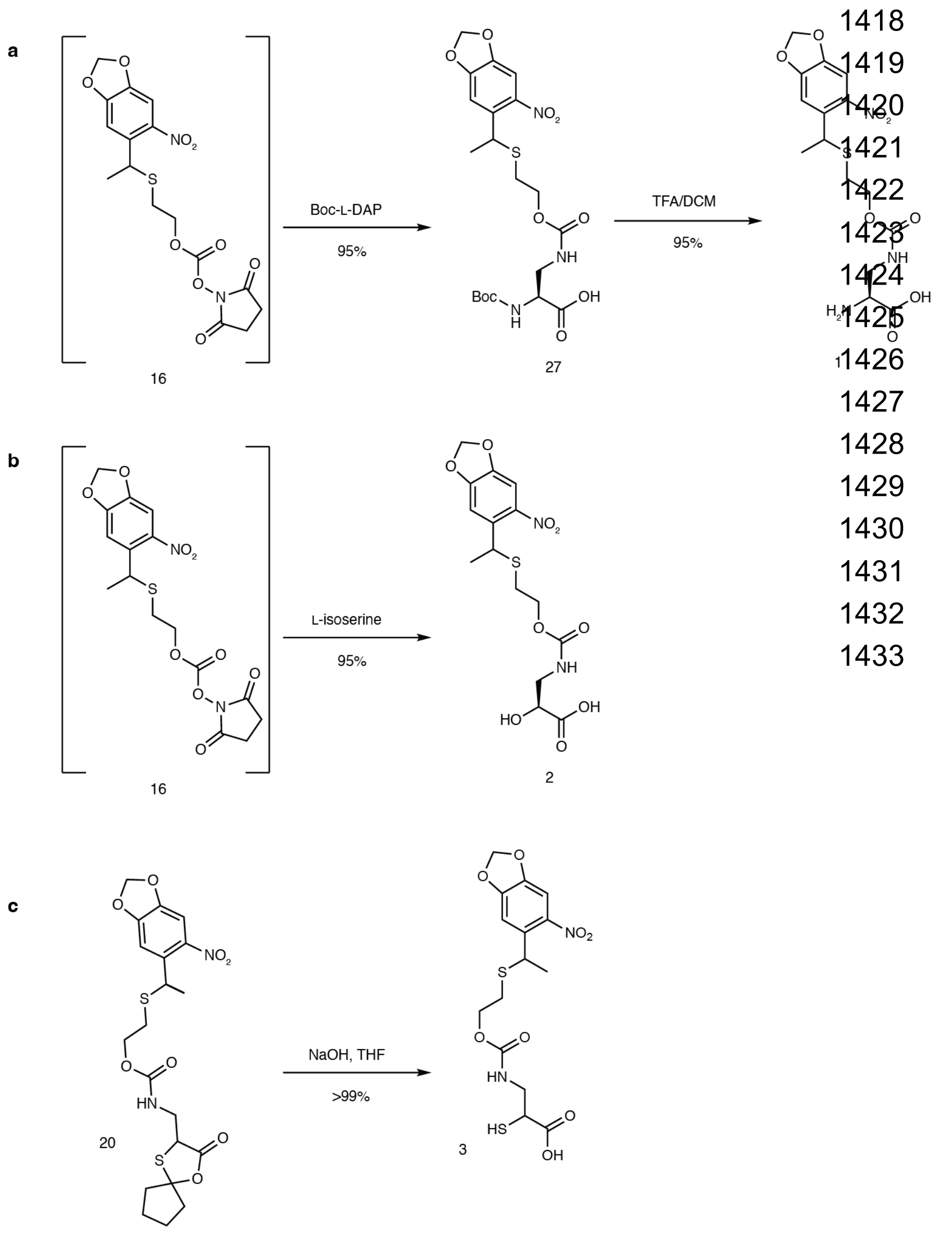

**Supplementary Fig. 3. Synthetic routes to BEAR *in vivo* substrates pm-DAP (1), pm-isoS (2), and pm-isoC ((±)-3).** (A) Solution-phase preparation of pm-DAP **1**. (B) Solution-phase preparation of pm-isoS **2**. (C) Solution-phase preparation of pm-isoC, **(±)-3**.

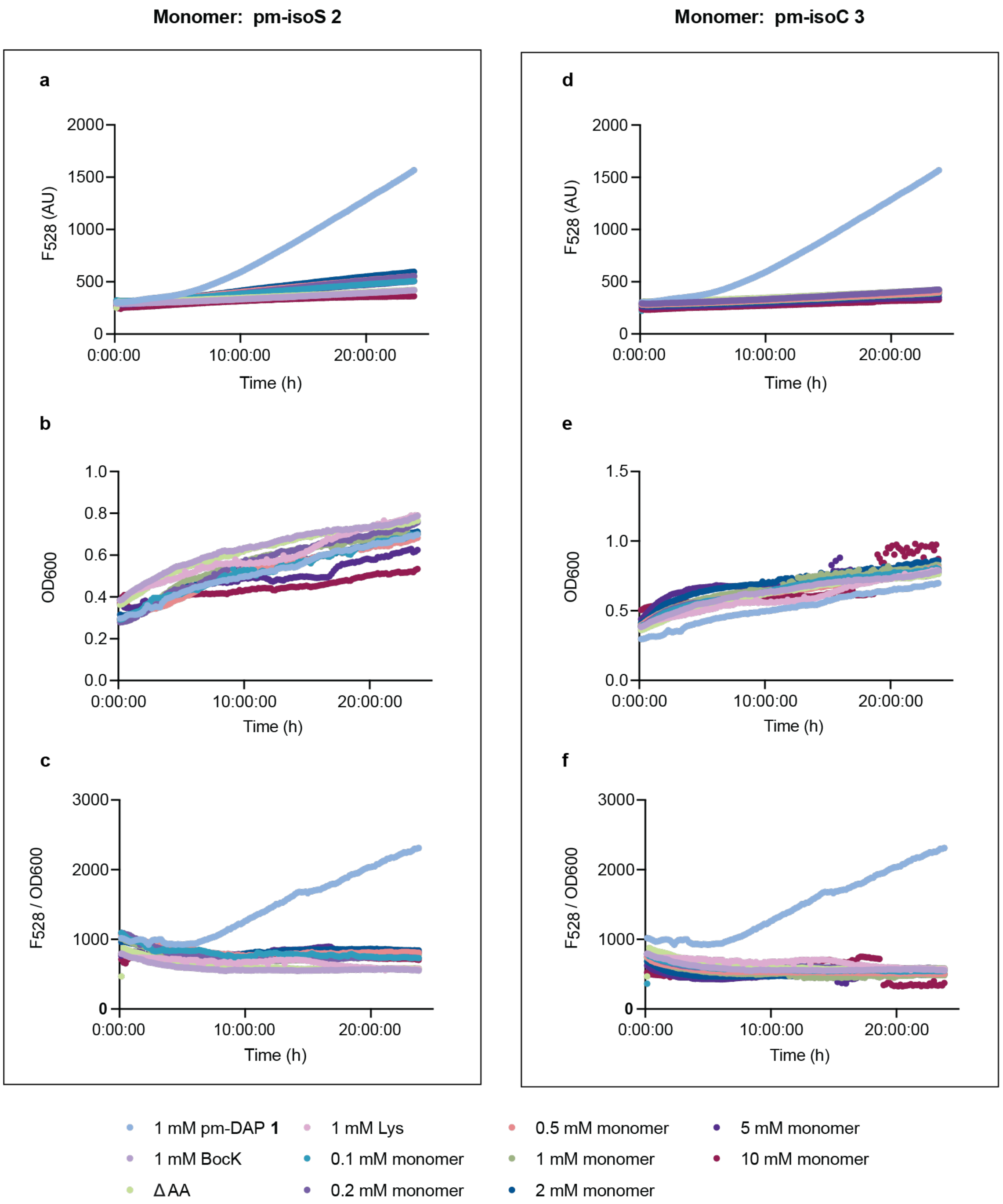

**Supplementary Fig. 4. $F_{528}$ and $OD_{600}$ of DH10B *E. coli* growths transformed with pBK-DAPRS and pET15A-sfGFP150TAG.** Plots showing (A) fluorescence at the emission maximum of GFP, $F_{528}$, (B) $OD_{600}$ , or (C) $F_{528}/OD_{600}$ over 24 h as a function of the concentration of pm-DAP **1** or pm-isoS **2** as well as BocK, Lys, and in the absence of any added substrate (ΔAA). Plots showing (D) fluorescence at the emission maximum of GFP, $F_{528}$, (E) $OD_{600}$ , or (F) $F_{528}/OD_{600}$ over 24 h as a function of the concentration of pm-DAP **1** or pm-isoC **(±)-3** as well as BocK, Lys, and in the absence of any added substrate (ΔAA).

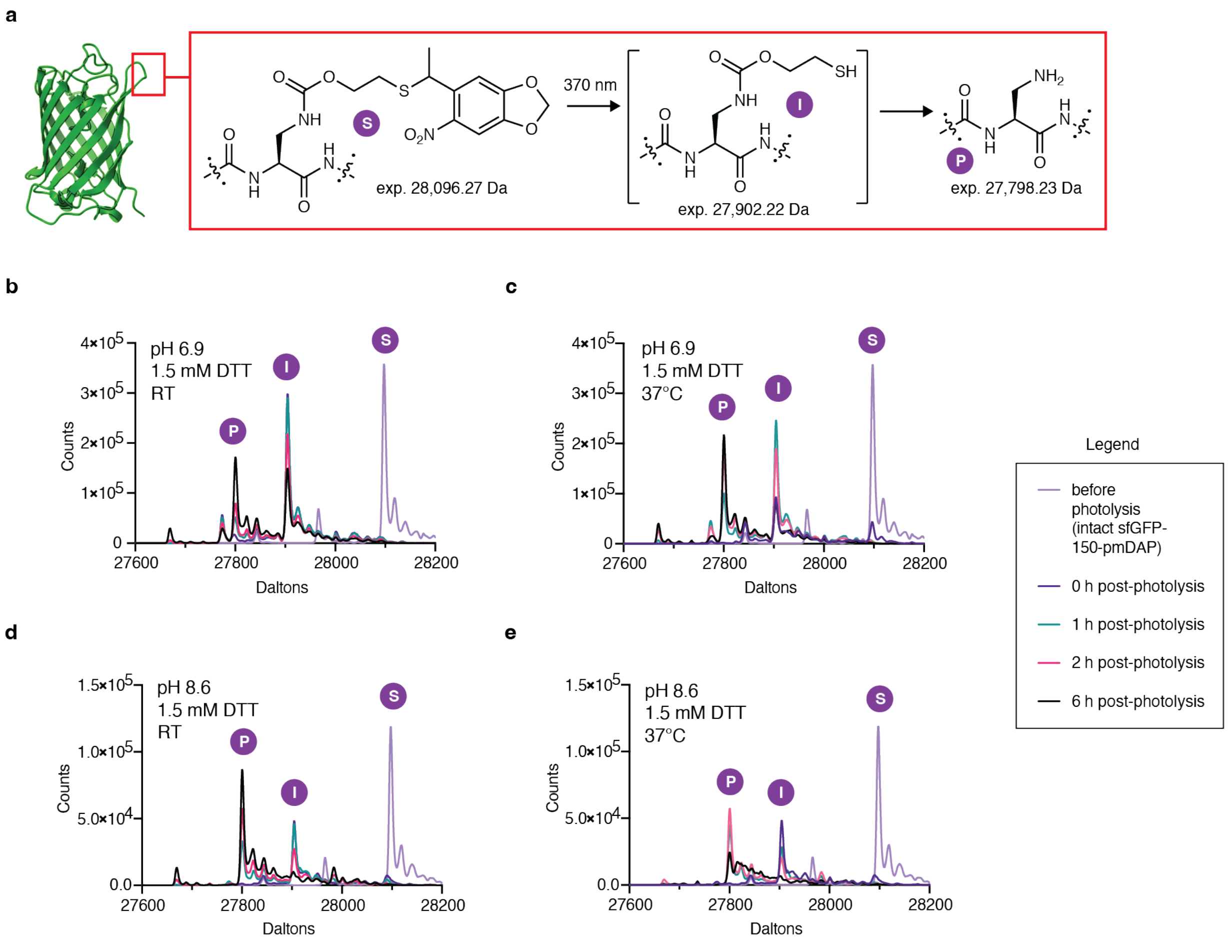

**Supplementary Fig. 5. Optimizing conditions for photo-demasking of pm-DAP 1 incorporated into sfGFP at position 150.** (A) Scheme illustrating the changes in mass expected of sfGFP containing pm-DAP at position 150 (**S**), is converted first into intermediate **(I)** and then into the completely unmasked product (**P)** containing diaminopropionic acid (DAP). Samples were treated as described in Methods. PDB ID 2B3P. (B-E) LC-MS time-course analysis of reactions incubated under the indicated conditions with the substrate (**S**), intermediate (**I**), and product (**P**) identified.

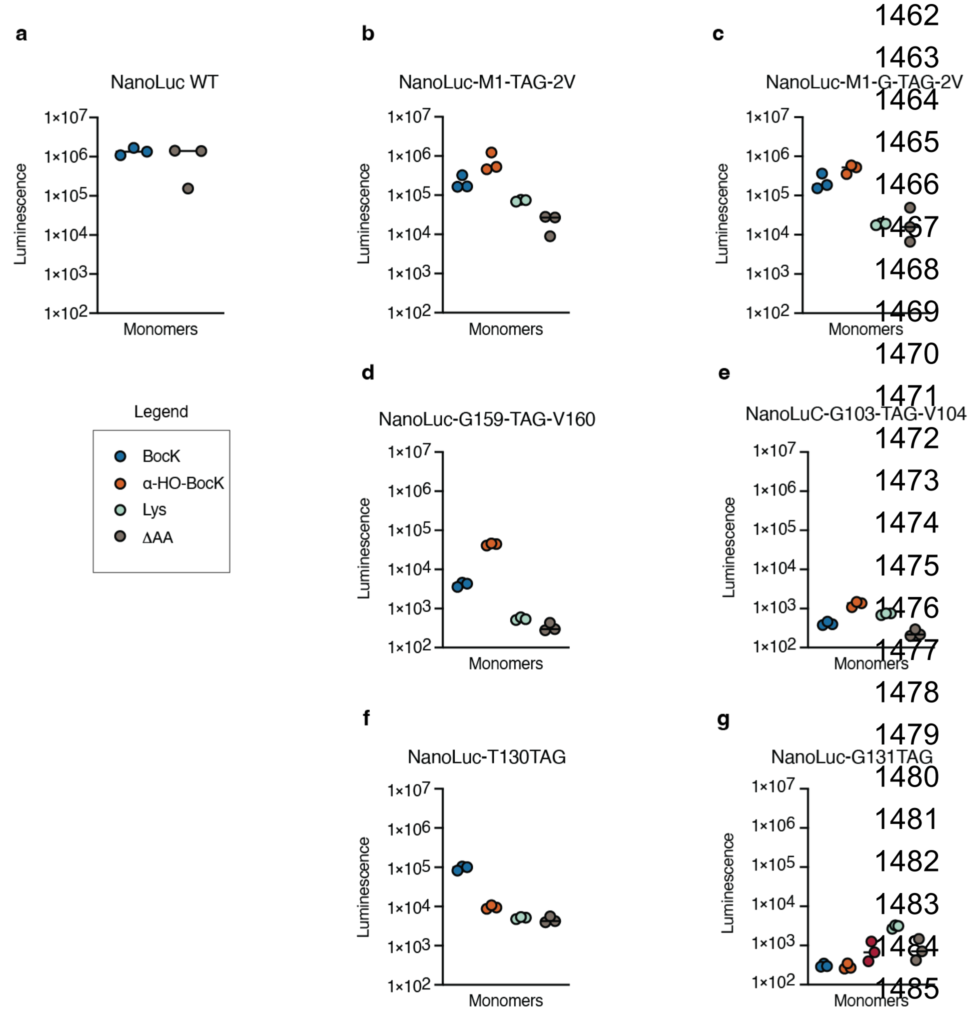

**Supplementary Fig. 6. Comparison of NanoLuc reporter plasmids.** Six reporter plasmids were prepared (see Methods) encoding NanoLuc in place of sfGFP and with a single TAG codon at various positions throughout the structure. Two plasmids encoded NanoLuc with a TAG codon substituted for or inserted after a residue near the N-terminus (G2TAG and G2-TAG-V3), two contained an inserted TAG codon within a loop (G159-TAG-V160 and G103-TAG-V104), and two positioned the TAG codon within a β-sheet (T130-TAG and G131-TAG). To validate the plasmids, BL21(DE3) *E. coli* were co-transformed separately with each NanoLuc reporter plasmid and pEVOL-*Ma*PylRS, which encodes the *Methanosarcina alvus* PylRS/$tRNA^{Pyl}_{CUA}$ pair. In each case, the cells were grown for 18 h. After 18 h, luminescence was determined using the Nano-Glo® Luciferase Assay System (Promega). (A) Wildtype (WT) NanoLuc expressed in the presence of BocK or α-OH-BocK as a positive control. (B-G) The NanoLuc variant specified was expressed in the presence of BocK or α-OH-BocK (positive controls) or Lys or no substrate ΔAA **(**negative controls).

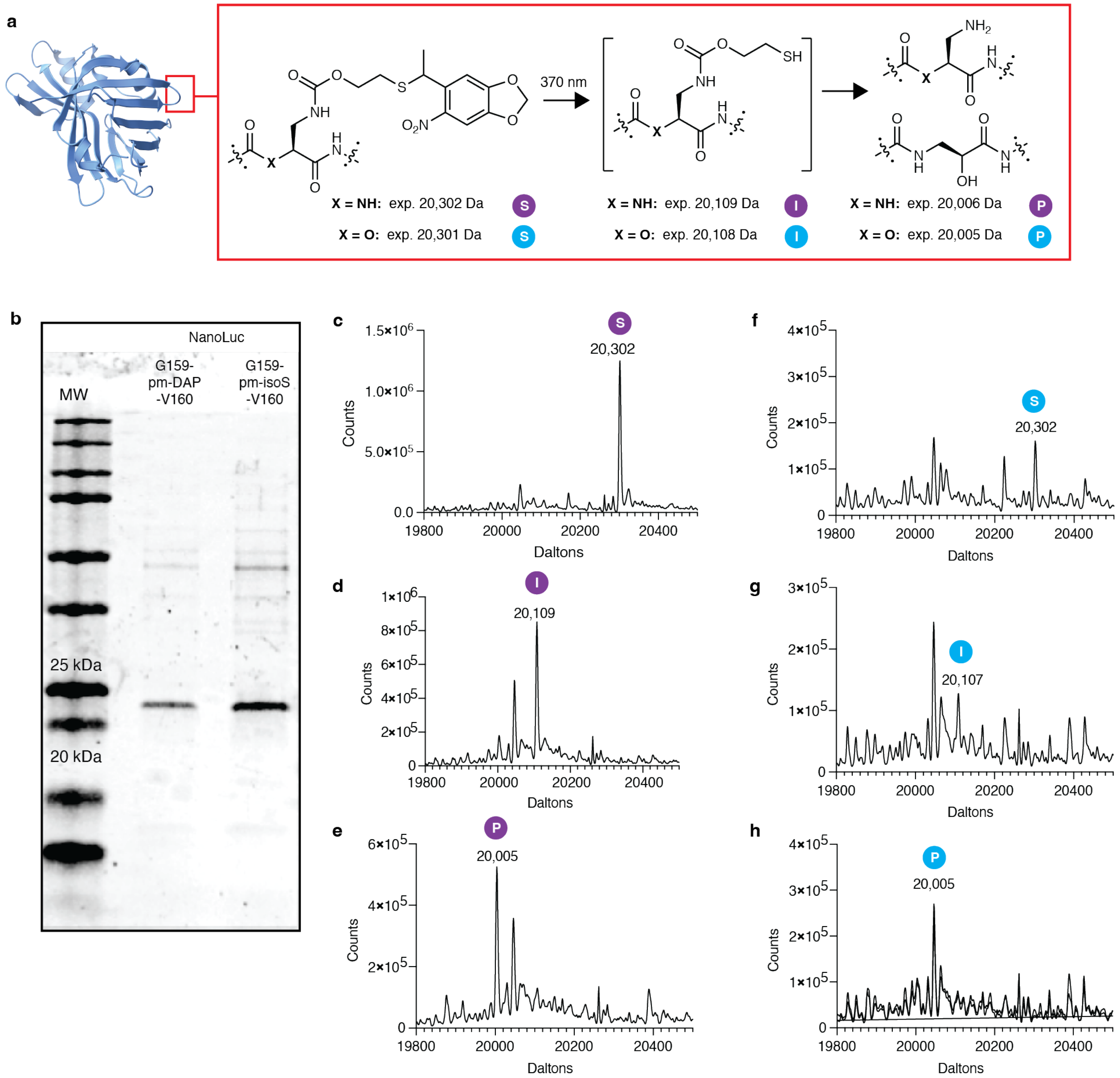

**Supplementary Fig. 7. Incorporation of pm-DAP and pm-isoS into NanoLuc G159-TAG-V160 and LC-MS analysis of photo-unmasking.** (A) General scheme illustrating the mass changes expected upon photo-unmasking of NanoLuc containing either pm-DAP or pm-isoS between residues G159 and V160. PDB ID 7SNS. (B) SDS-PAGE analysis of purified NanoLuc-G159-pm-DAP-V160 and NanoLuc-G159-pm-isoS-V160. (C) Deconvoluted mass spectrum of affinity-purified NanoLuc G159-pm-DAP-V160 prior to photolysis; (D) immediately after photolysis at 370 nm irradiation; and (E) 18 h after photolysis. (F) Deconvoluted mass spectrum of affinity-purified NanoLuc G159-pm-isoS-V160 prior to photolysis; (D) immediately after photolysis at 370 nm irradiation; and (E) 18 h after photolysis.

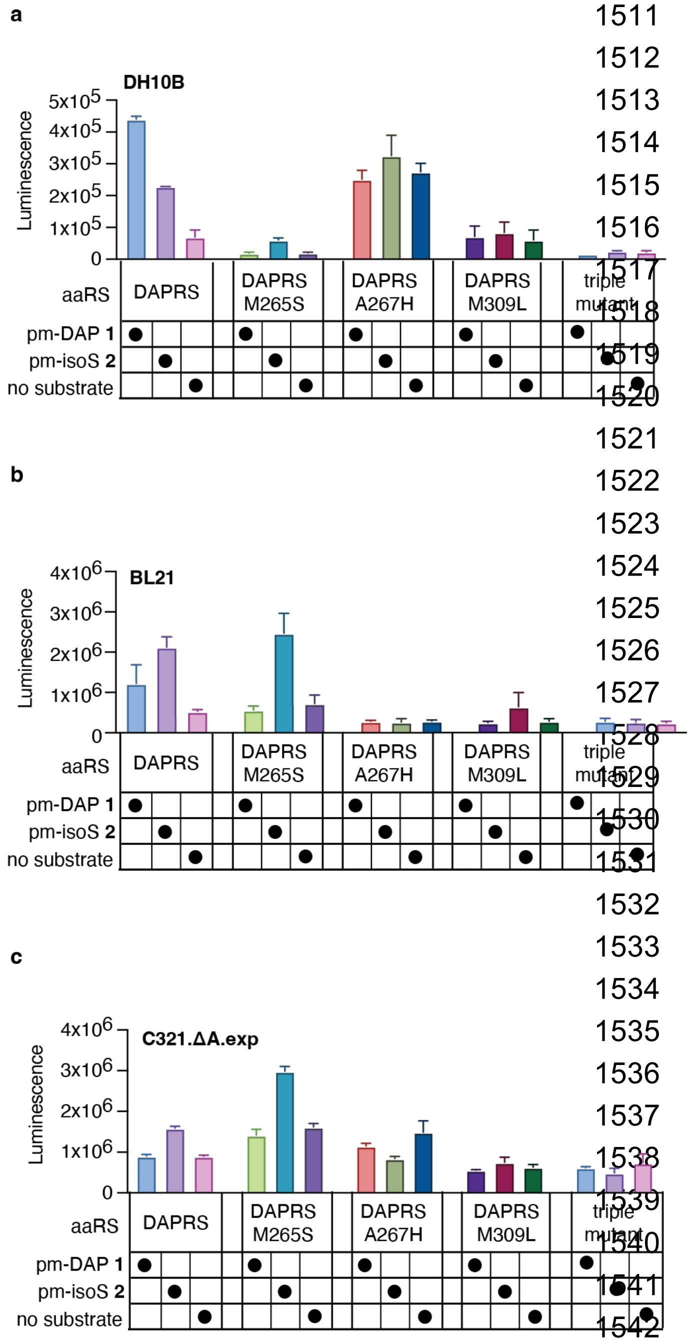

**Supplementary Fig. 8**. **Evaluation of NanoLuc activity in cells expressing select DAPRS mutants and supplemented with pm-DAP 1 or pm-isoS 2.** All plots show luminescence of (A) DH10B *E. coli*; (B) BL21 *E. coli*; or (C) C321.ΔA.exp *E. coli* at the 18 h time point for growths expressing DAPRS variants encoded by either pBK-DAPRS-M265S, pBK-DAPRS-A267H, pBK-DAPRS-M309L, or pBK-DAPRS-M265S-A267H-M309L and supplemented with either 1 mM pm-DAP **1**, 1 mM pm-isoS **2**, or in the absence of any amino or hydroxy acid (ΔAA). NanoLuc was isolated from C321.ΔA.exp cells expressing DAPRS-M265S, purified, and subject to peptide mapping analysis (See Supplementary Fig. 9). This analysis revealed extremely low, but detectable incorporation of pm-isoS **2** in samples grown in C321.ΔA.exp *E. coli.*

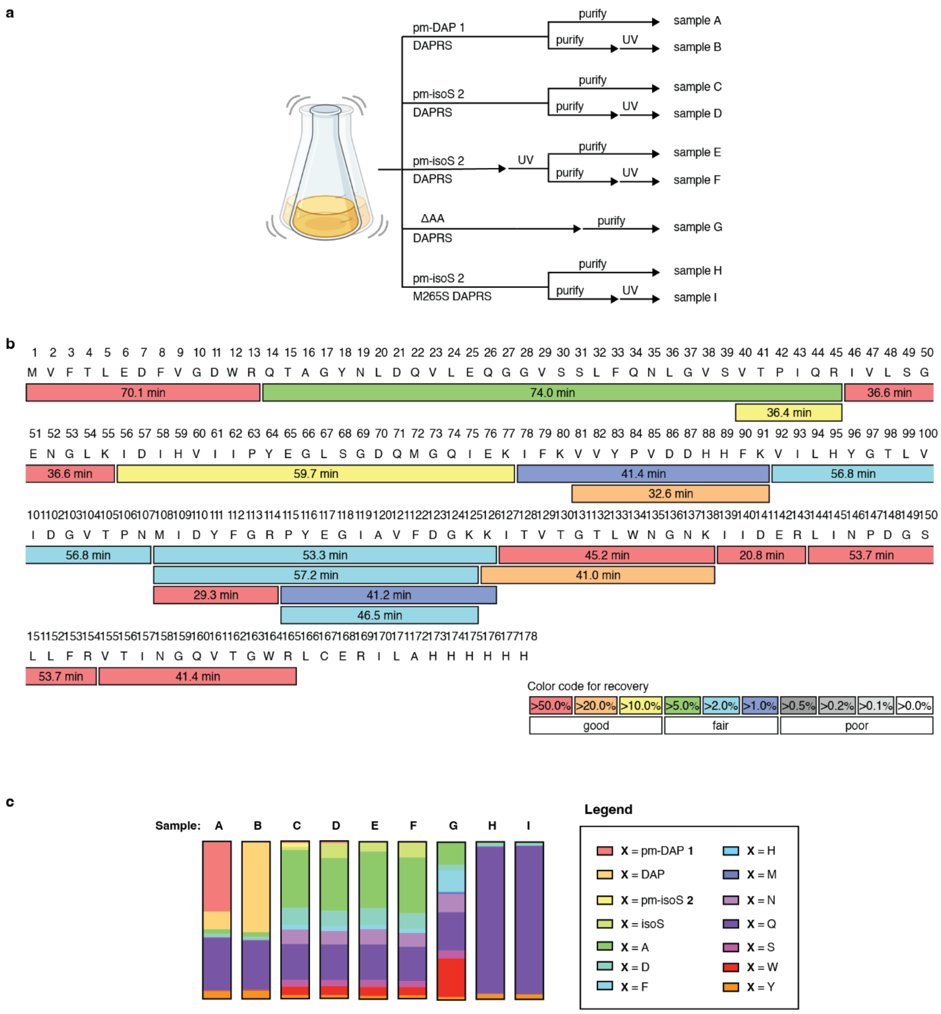

d

| Sample: | A | B | C | D | E | F | G | H | I |
|---|---|---|---|---|---|---|---|---|---|
| X = pm-DAP **1** | 44.64% | 0.06% | 1.12% | 0.00% | 0.00% | 0.00% | 0.00% | 0.00% | 0.00% |
| X = DAP | 11.00% | 57.53% | 0.57% | 1.40% | 0.35% | 0.67% | 0.00% | 0.00% | 0.10% |
| X = pm-isoS **2** | 0.15% | 0.00% | 2.09% | 0.00% | 0.15% | 0.00% | 0.00% | 0.00% | 0.00% |
| X = isoS | 0.00% | 0.00% | 1.89% | 9.50% | 5.79% | 9.35% | 0.00% | 0.54% | 0.51% |
| X = A | 2.88% | 2.85% | 36.60% | 33.43% | 35.94% | 35.53% | 14.32% | 0.41% | 0.00% |
| X = D | 1.89% | 1.76% | 10.82% | 10.00% | 10.74% | 10.02% | 3.98% | 1.82% | 1.56% |
| X = F | 0.11% | 0.10% | 2.79% | 2.66% | 2.34% | 2.57% | 12.54% | 0.00% | 0.24% |
| X = H | 0.05% | 0.04% | 0.32% | 0.28% | 0.35% | 0.13% | 0.71% | 0.00% | 0.00% |
| X = M | 0.01% | 0.00% | 0.13% | 0.09% | 0.17% | 0.12% | 1.41% | 0.23% | 0.00% |
| X = N | 0.49% | 0.48% | 8.98% | 8.39% | 9.50% | 8.62% | 11.45% | 0.18% | 0.00% |
| X = Q | 33.31% | 31.35% | 22.94% | 22.60% | 22.62% | 21.82% | 24.28% | 93.29% | 94.36% |
| X = S | 0.19% | 0.15% | 4.08% | 3.91% | 4.30% | 3.89% | 5.02% | 0.27% | 0.00% |
| X = W | 0.39% | 0.46% | 5.57% | 5.76% | 5.70% | 5.25% | 24.51% | 0.00% | 0.13% |
| X = Y | 4.90% | 5.22% | 2.09% | 1.98% | 2.06% | 2.02% | 1.79% | 3.26% | 3.10% |

**Supplementary Fig. 9. Quantitative analysis of incorporation of pm-DAP 1 and pm-isoS 2 and subsequent BEAR reaction.** (A) Nine samples (A-I) were analyzed using quantitative tryptic peptide mapping. (B) Sequence coverage map for peptide mapping of NanoLuc G159-TAG-V160. Colored sections indicate expected peptide fragments and found retention times during LC-MS/MS analysis as well as the location of the tryptic product containing residues 155-165 and an additional residue **X** between residues G159 and V160. (C) Plots illustrating the identity of **X** for Samples A-I as described in **Figure 5**. (D) Data corresponding to the plots in panel B.

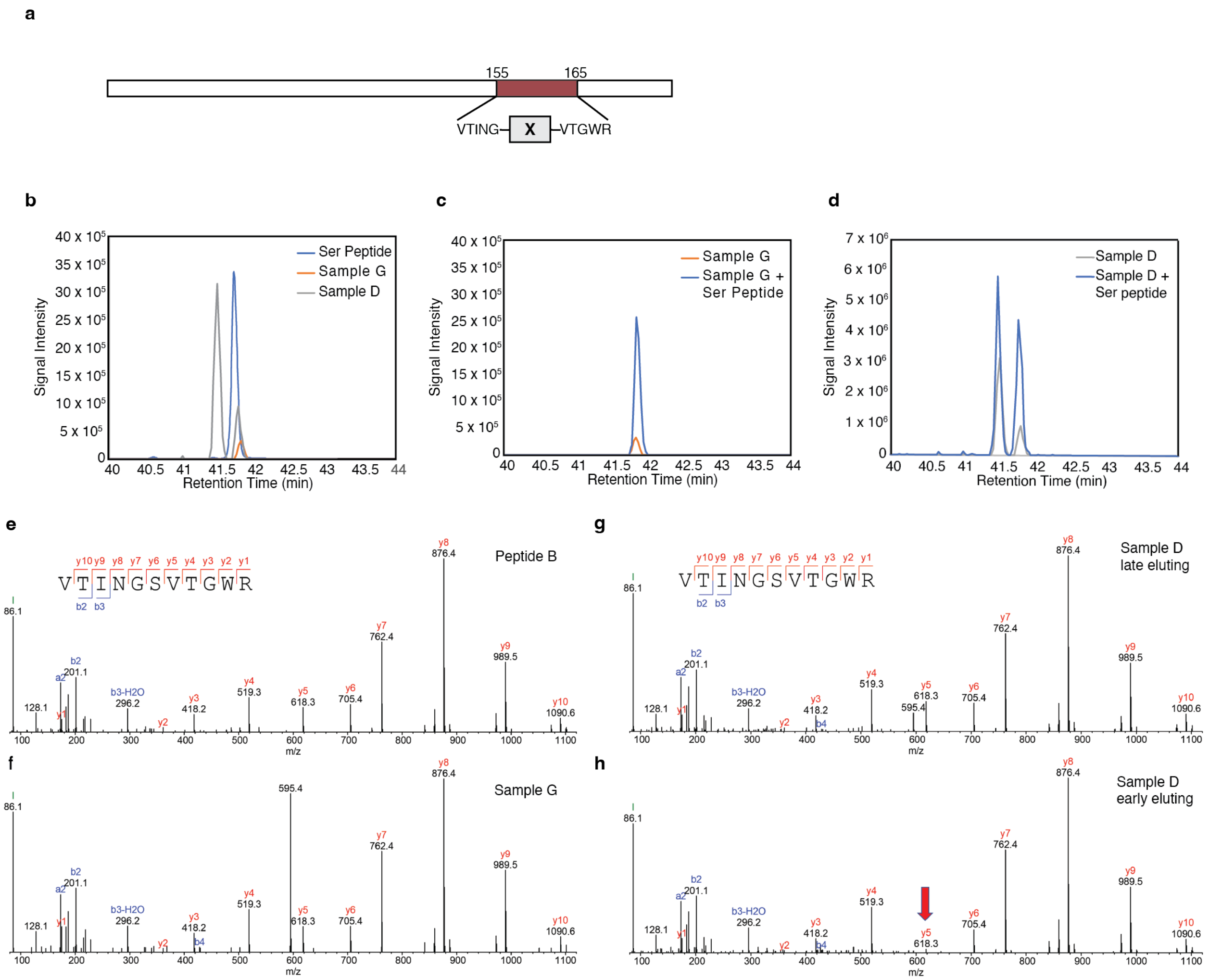

**Supplementary Fig. 10**. **The tryptic product from Sample D does not co-elute with a synthetic peptide containing Ser between G159 and V160.** (A) Digestion of NanoLuc with trypsin generates a tryptic fragment comprising residues 155-165 (including the expected site of incorporation of monomer **X** between native G159 and V160). (B) Comparison of the retention times of the tryptic products present in Samples D and G to that of synthetic Peptide B whose sequence corresponds to the tryptic product where **X** = Ser. The retention time of Peptide B (blue) matches that of the late-eluting species in Sample D (grey) (also shown in **Figure 5D**), and the peak observed in negative control Sample G (orange). (C) Comparison of the tryptic product in Sample G (orange) to that of a co-injection of Sample G and Peptide B (blue). (D) Comparison of the tryptic product in Sample D (gray) to that of a co-injection of Sample D and Peptide B. These spiking experiments demonstrate that the late-eluting species in Sample D is identical to Peptide B, where **X** = Ser. (E) The MS/MS spectrum of synthetic Peptide B is identical to that of (F) the tryptic fragment from Sample G and (G) the late-eluting isomeric peptide in Sample D. (H) The early-eluting isomeric peptide in Sample D has a different MS/MS (lower y5 indicated by a red arrow). Note that the presence of an ion at m/z 595.4 in some samples is due to co-isolation of an interfering ion at that m/z (precursor ion m/z of the peptide is 595.3).

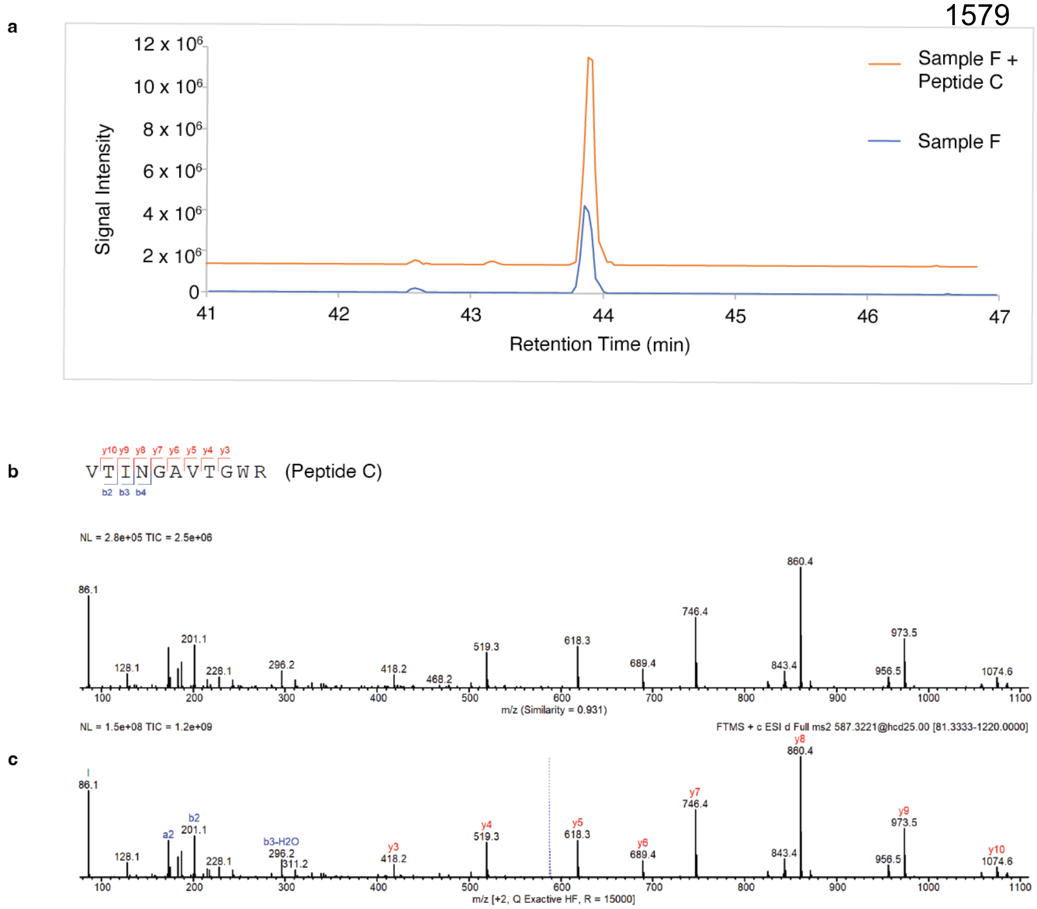

**Supplementary Fig. 11. The amino acid at location 160 in the detected "Q160A" tryptic peptide is alanine.** (A) Extracted ion chromatograms of the tryptic peptide comprising NanoLuc residues V155 through R165 from Sample F containing (orange) or without (blue) the synthetic Ala-containing Peptide C. Addition of Peptide C to Sample F leads to an increase in the intensity of the Sample F tryptic peptide without peak splitting. This observation indicates that the detected Q160A species in the Sample F tryptic digest has the same retention time as the synthetic Ala-containing Peptide C, and confirms that the amino acid at location X is indeed alanine. (B) The MS/MS spectrum of the tryptic peptide comprising NanoLuc residues V155 through R165 obtained in the Sample F tryptic digest is identical to the MS/MS spectrum of (C) the synthetic Ala-containing Peptide C.

### Supplementary Tables

**Supplementary Table 1. Oligonucleotides used in this study.**

| Name | Sequence (5' to 3') |
|---|---|
| LTR1 | ggttaattcctcctgttagcccaaaaaacg |
| LTR2 | agctcgagcgaagcttggg |
| LTR3 | gctaacaggaggaattaaccatggtgtttaccctggaagattttgtgg |
| LTR4 | gcccaagcttcgctcgagctttagtgatggtgatggtgatgcgc |
| LTR5 | tcggggccagGCTcggacgcagg |
| LTR6 | ccctgtgtaactatctgcgtaaac |
| LTR7 | acagggtcggATGcagcatcgga |
| LTR8 | gtaactatctgcgtaaactg |
| LTR9 | aaaactgaacCAGggtgaattcttccag |
| LTR10 | gccaaatgggcagcggct |
| LTR11 | caaatgggcagcggctgcacc |
| LTR12 | gtgatatcacgttccagtttgcccagataatcttcacg |

**Supplementary Table 2. Masses searched during peptide mapping.** Masses for peptide fragments containing pm-DAP **1**, DAP, pm-isoS **2**, or isoS were calculated as variations to the mass expected if Gln was present between NanoLuc residues G159 and V160.

| Residue | Formula | Mass | ΔFormula | ΔMass |
|---|---|---|---|---|
| Glutamine (Q) | $C_5H_8N_2O_2$ | 128.05858 | | |
| pm-DAP **1** | $C_{15}H_{17}N_3O_7S$ | 383.07872 | $Q+C_{10}H_9NO_5S$ | Q+255.02014 |
| DAP | $C_3H_6N_2O$ | 86.04801 | $Q-C_2H_2O$ | Q-42.0106 |
| pm-isoS **2** | $C_{15}H_{16}N_2O_8S$ | 384.06274 | $Q+C_{10}H_8O_6S$ | Q+256.00416 |
| isoS | $C_3H_5NO_2$ | 87.03203 | $Q-C_2H_3N$ | Q-41.0265 |

### Processed NMR Spectra

Standard View for $^{1}H$ NMR is defined as 10 to -1 ppm.

Extended View for $^{1}H$ NMR is defined as the full collection range (14 to -2 ppm or 16 to -4 ppm).

Standard View for $^{13}C$ NMR is defined as 210 to -10 ppm.

Extended View for $^{13}C$ NMR is defined as the full collection range (235 to -15 ppm or 220 to -20 ppm).

$^1$H NMR (500 MHz, $CDCl_3$) – Standard view **(12)**

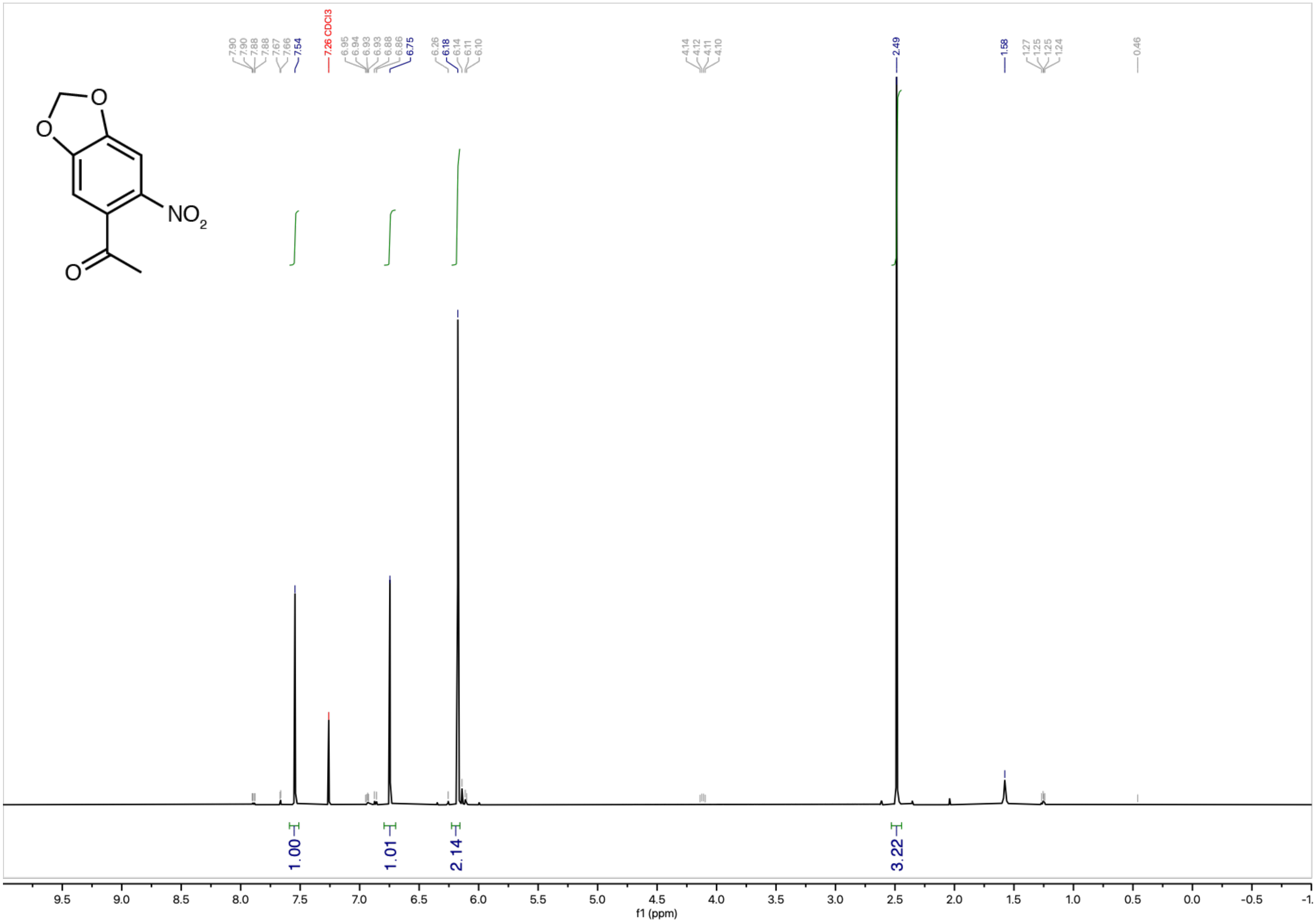

$^{13}C\{^{1}H\}$ NMR (126 MHz, $CDCl_3$) – Standard view **(12)**

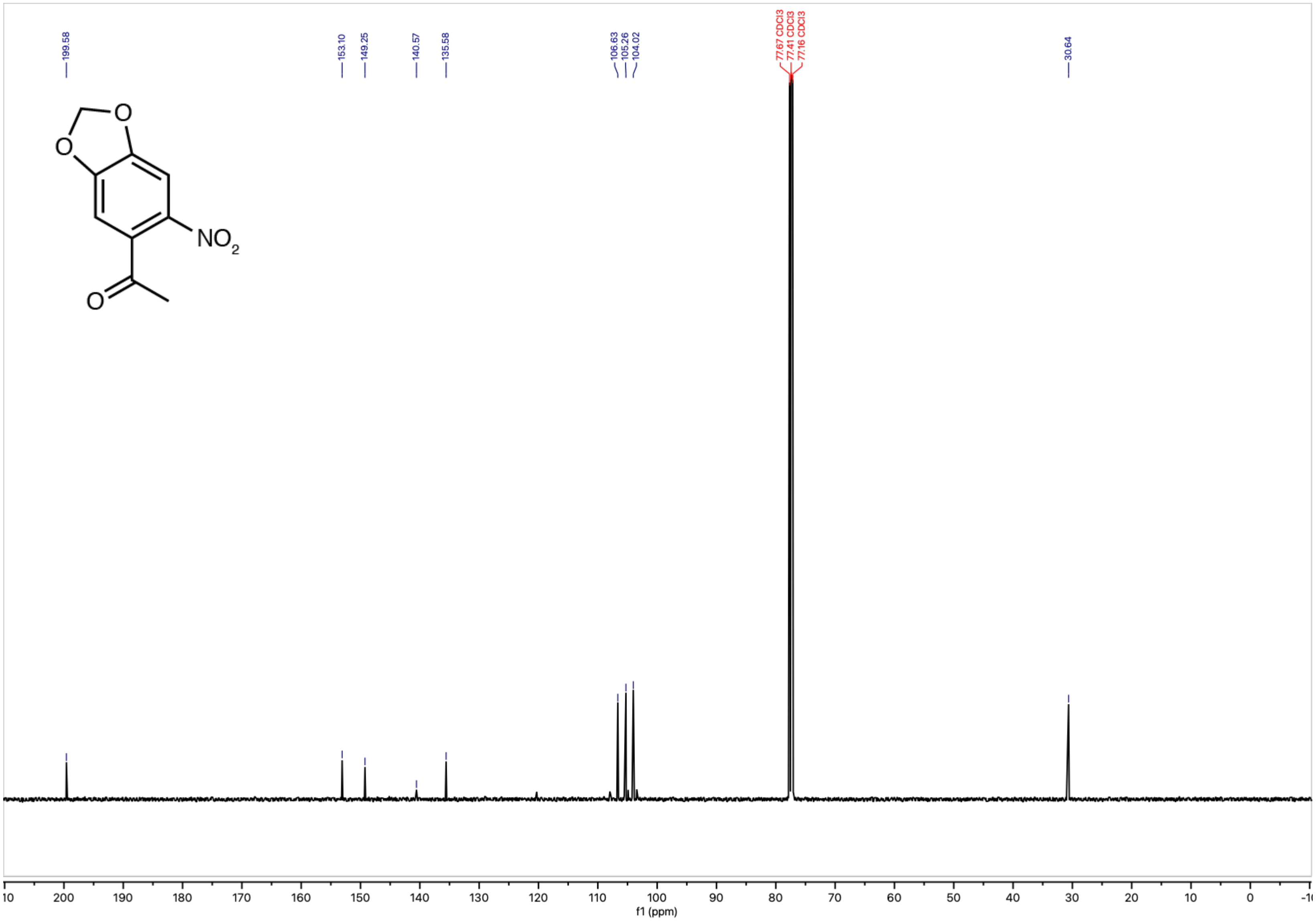

$^1$H NMR (500 MHz, $CDCl_3$) – Standard view **((±)-13)**

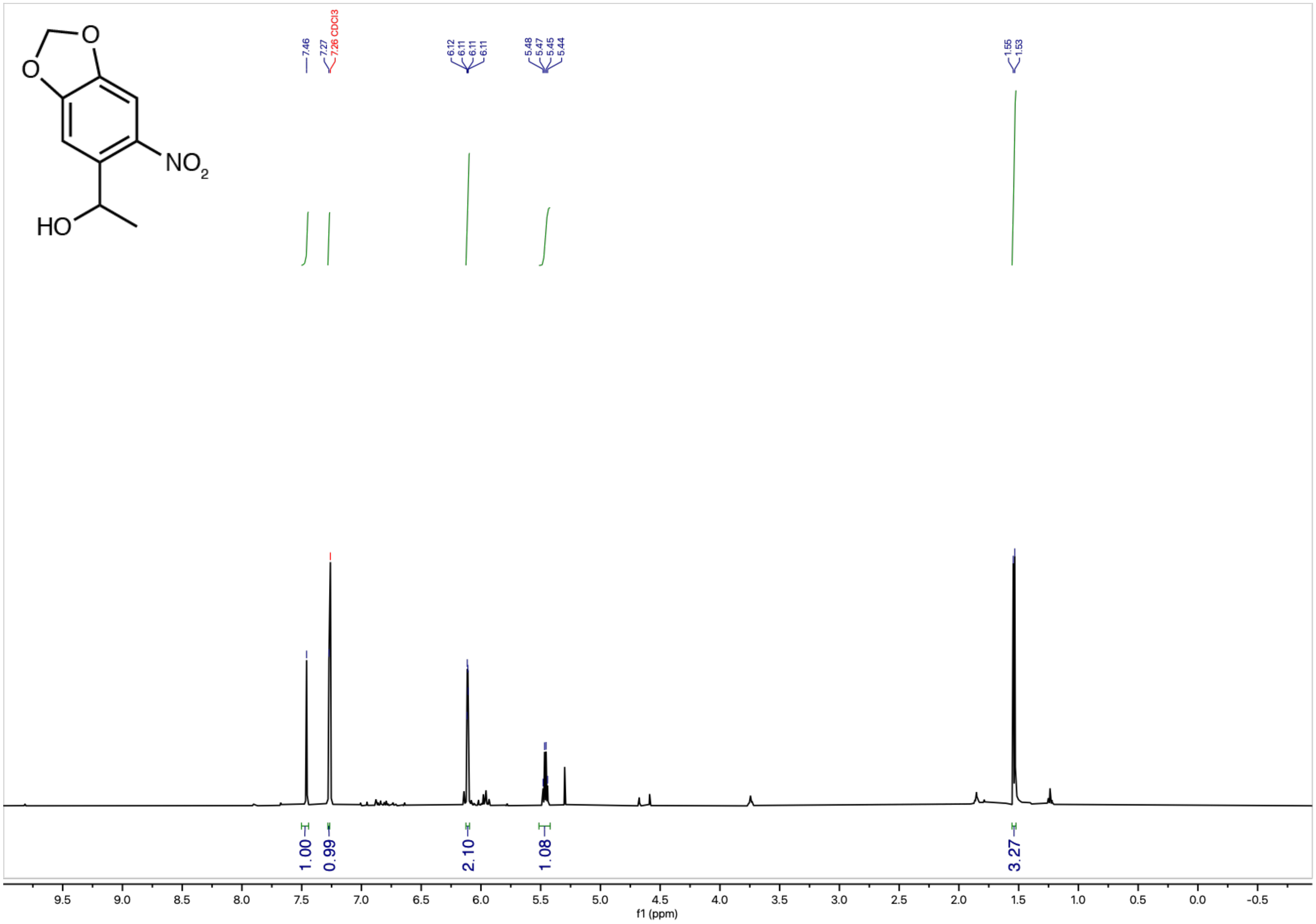

$^{13}C\{^1H\}$ NMR (126 MHz, $CDCl_3$) – Standard view **((±)-13)**

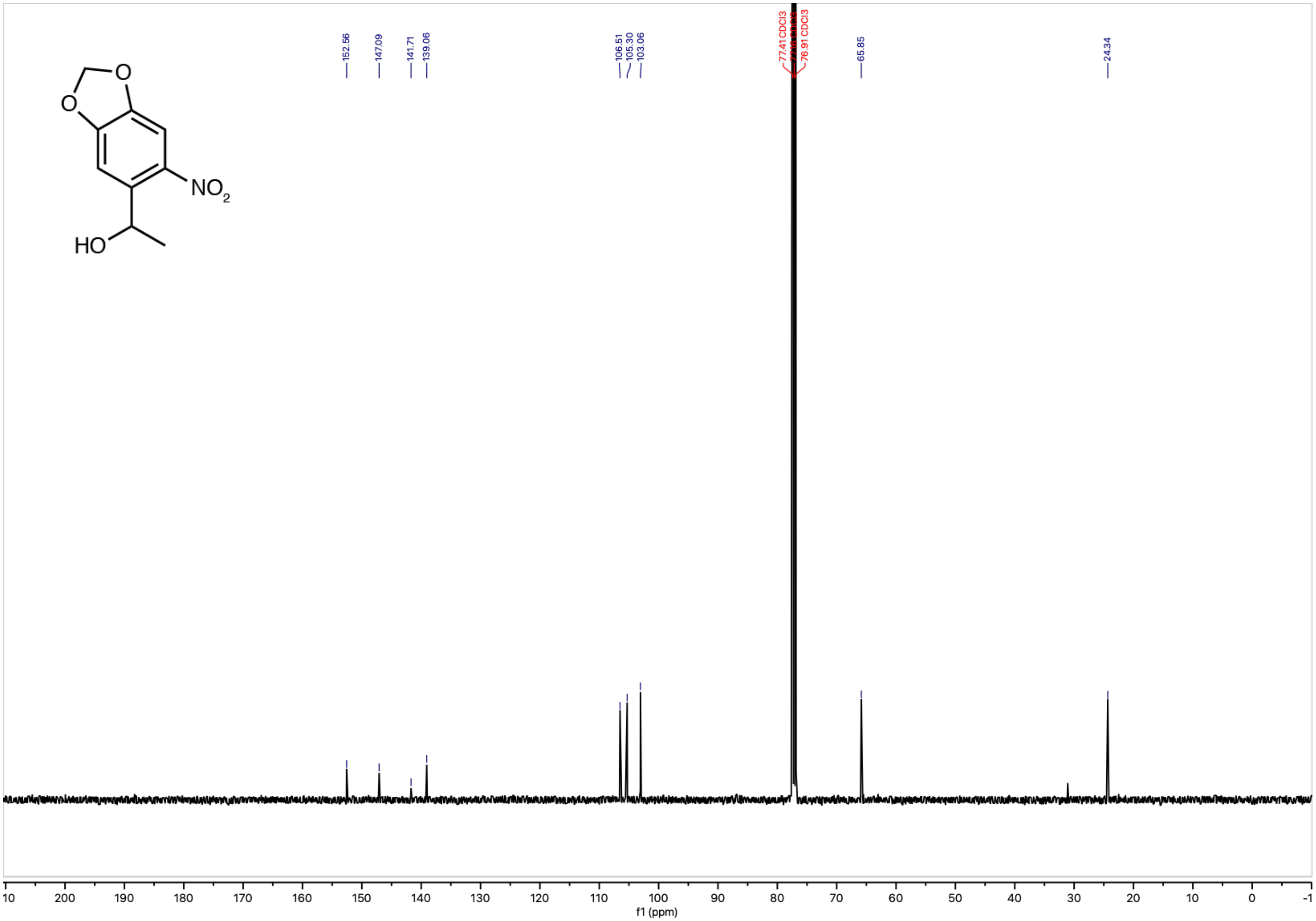

[1]H NMR (500 MHz, $CDCl_3$) – Standard view **((±)-14)**

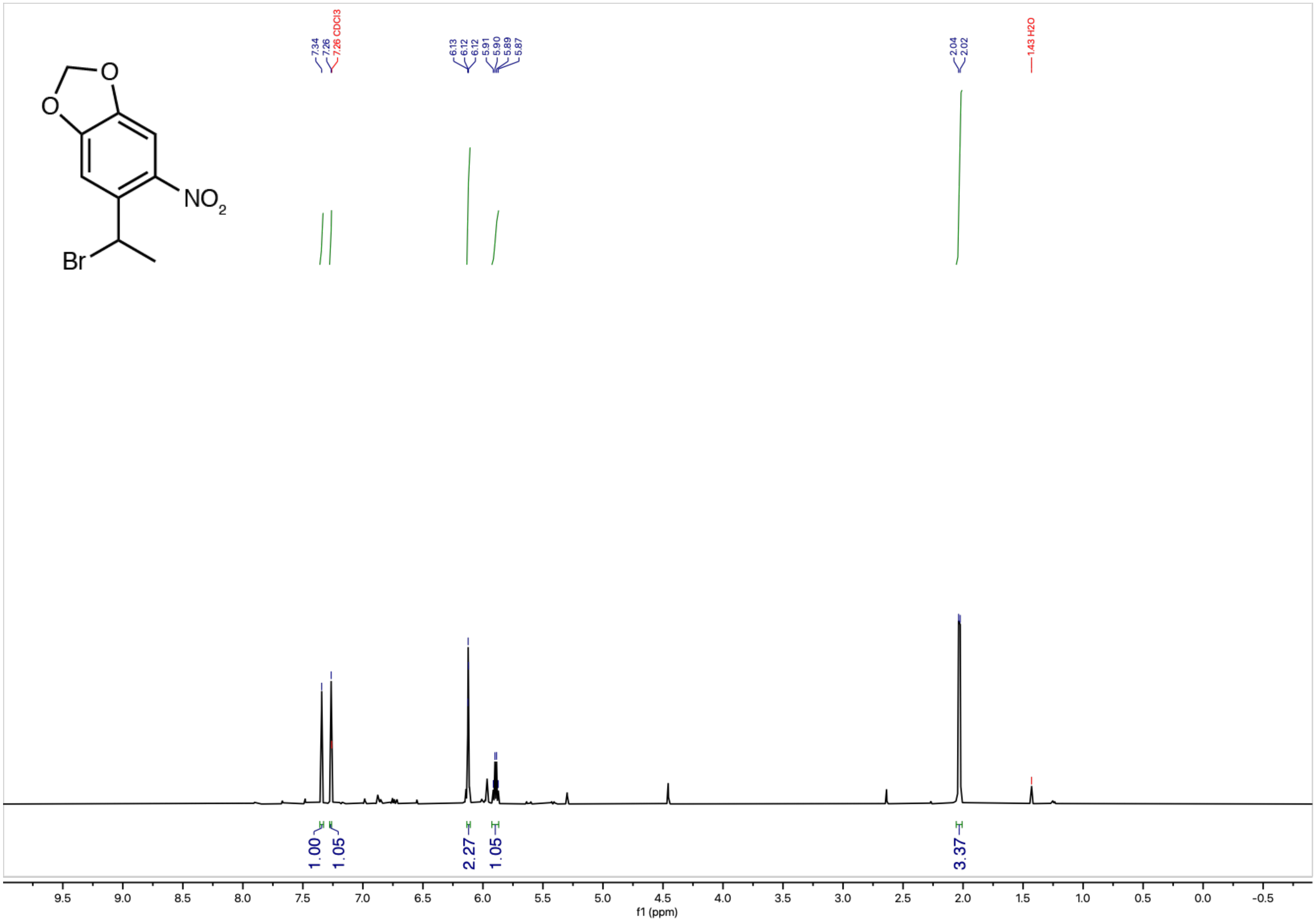

$^{13}C\{^{1}H\}$ NMR (126 MHz, $CDCl_3$) – Standard view **((±)-14)**

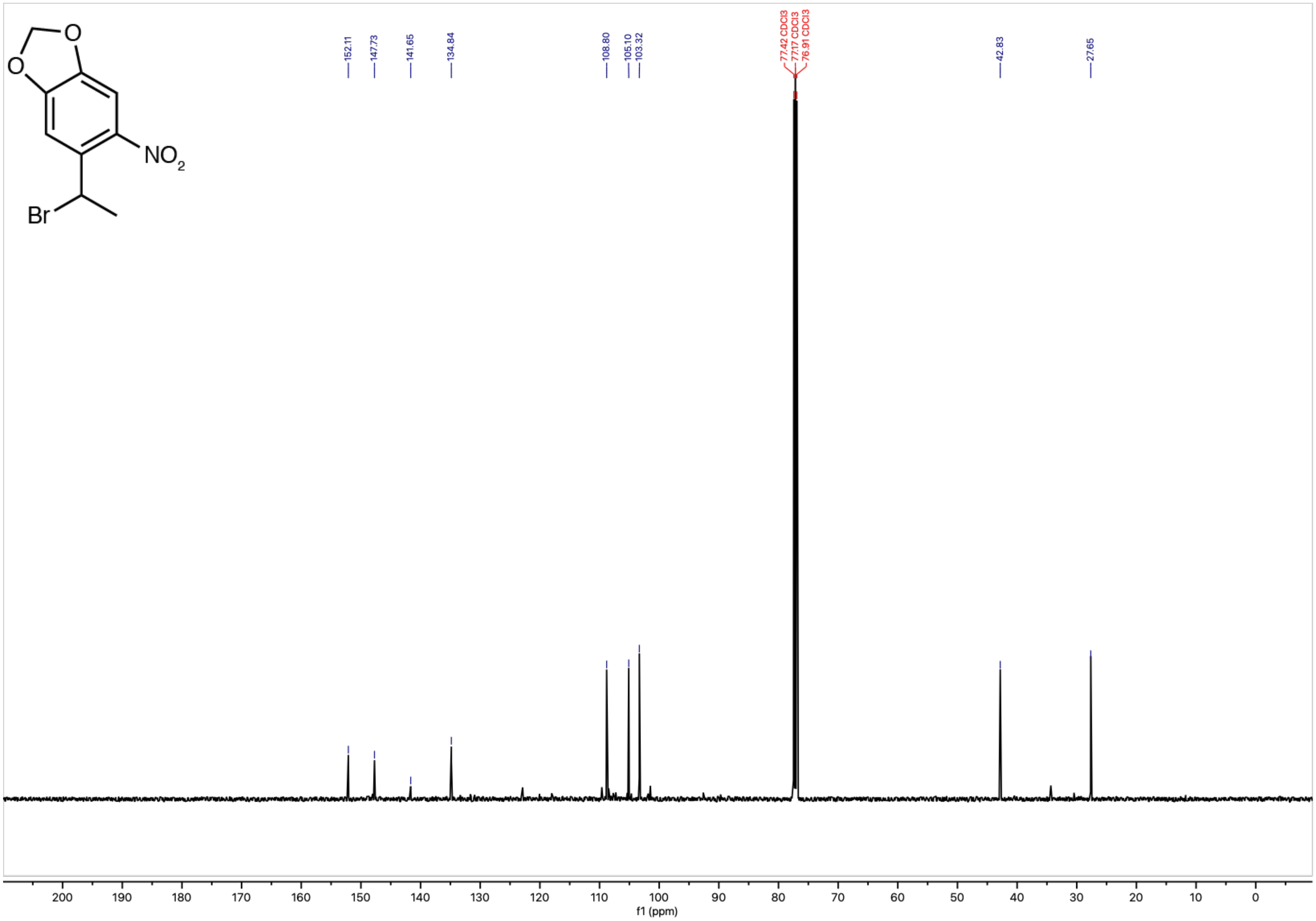

$^{1}H$ NMR (500 MHz, $CDCl_3$) – Standard view **((±)-15)**

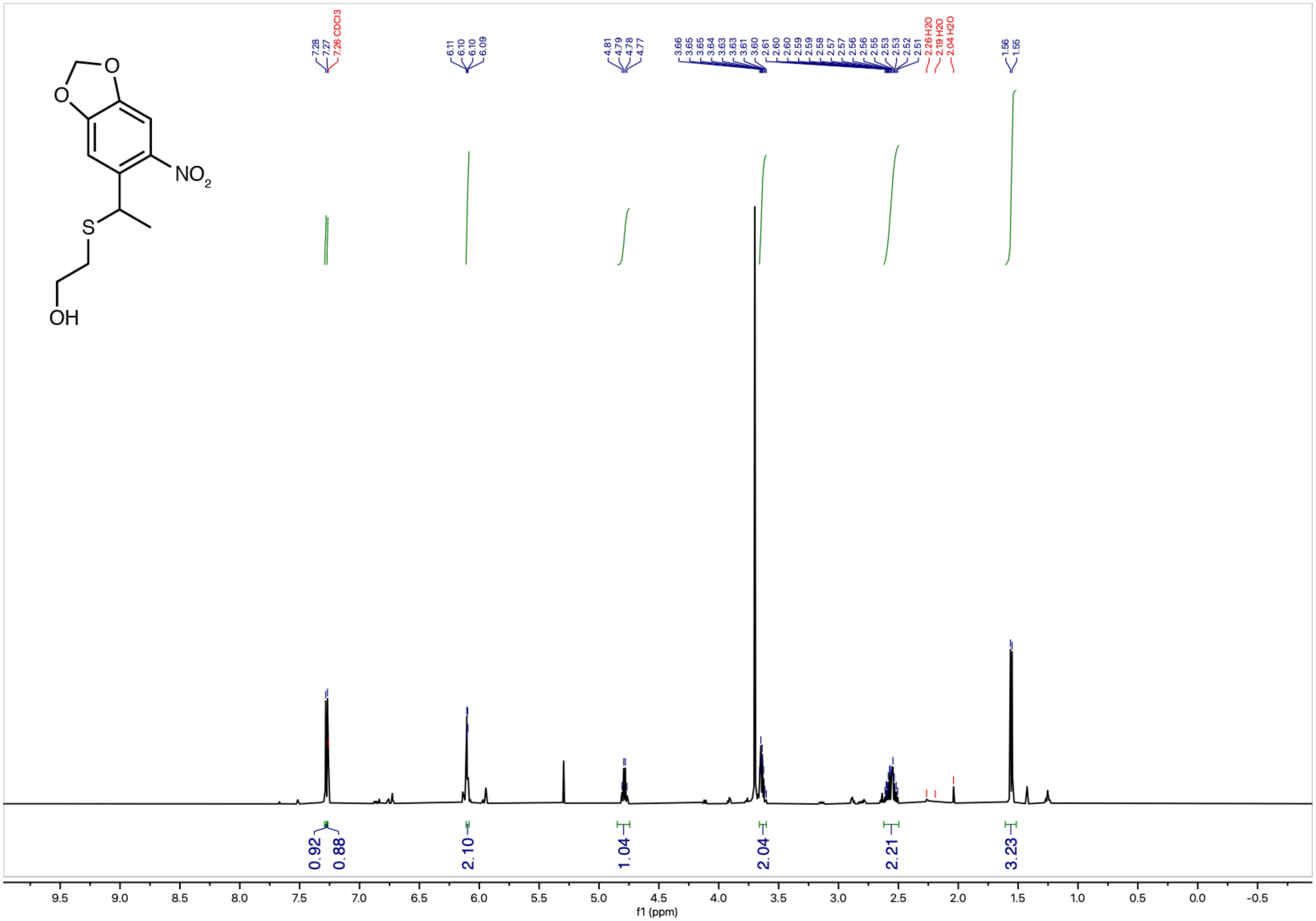

$^{13}C\{^{1}H\}$ NMR (126 MHz, $CDCl_3$) – Standard view **((±)-15)**

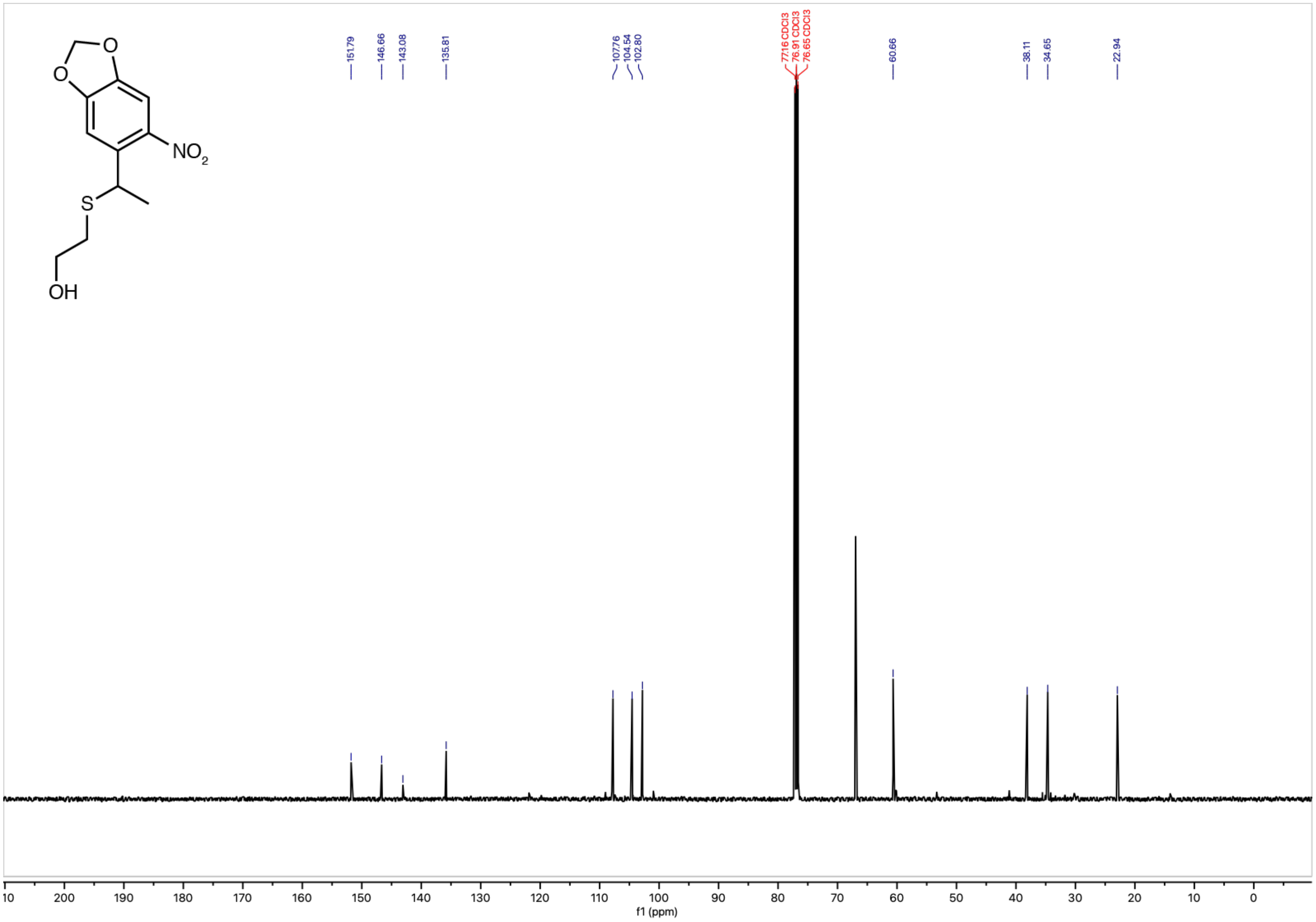

[1]H NMR (600 MHz, methanol-$d_4$) – Standard view **(17)**

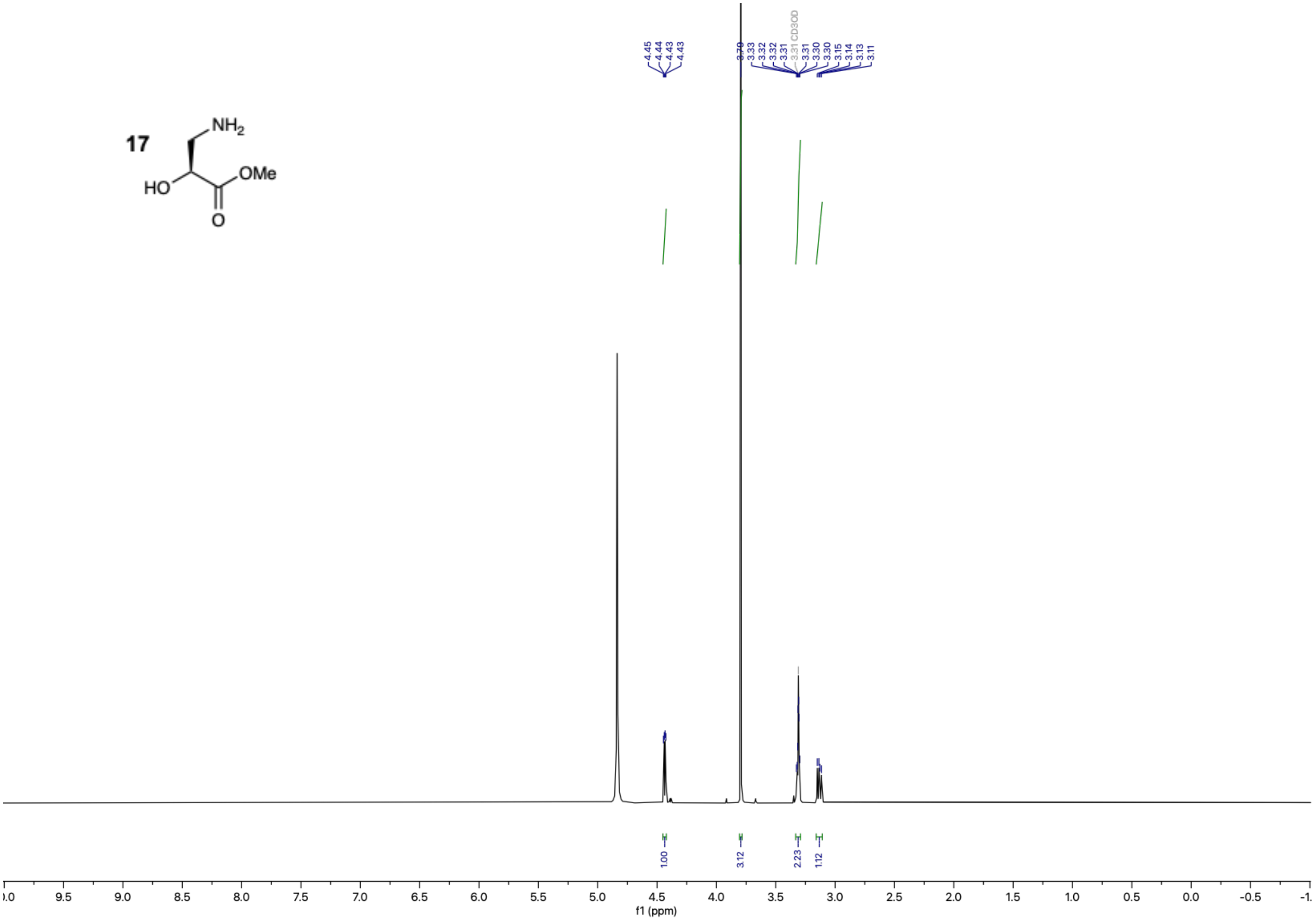

$^{13}C\{^{1}H\}$ NMR (151 MHz, methanol-$d_4$) – Standard view **(17)**

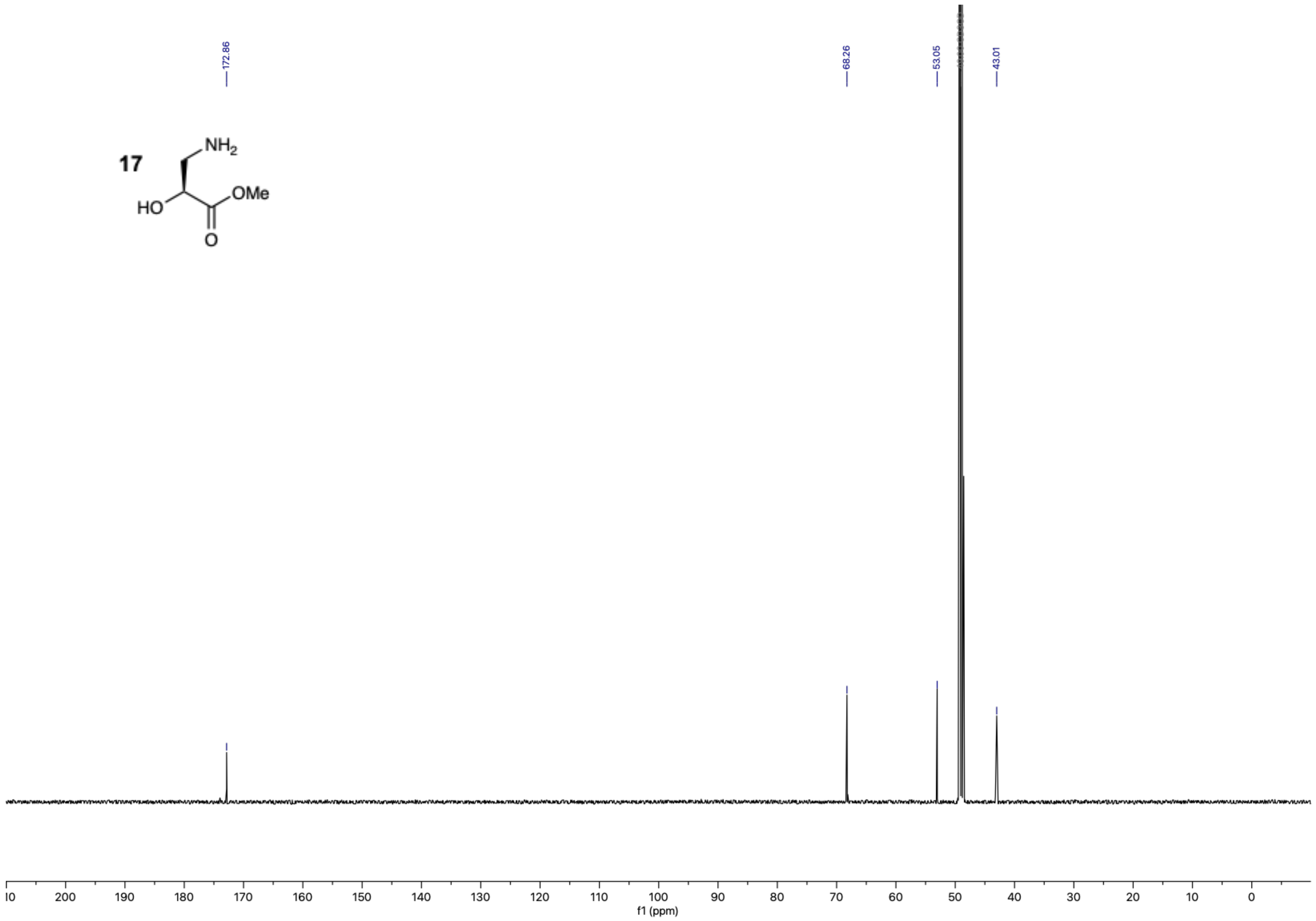

$^{1}$H NMR (600 MHz, methanol-$d_4$) – Standard view **(18)**

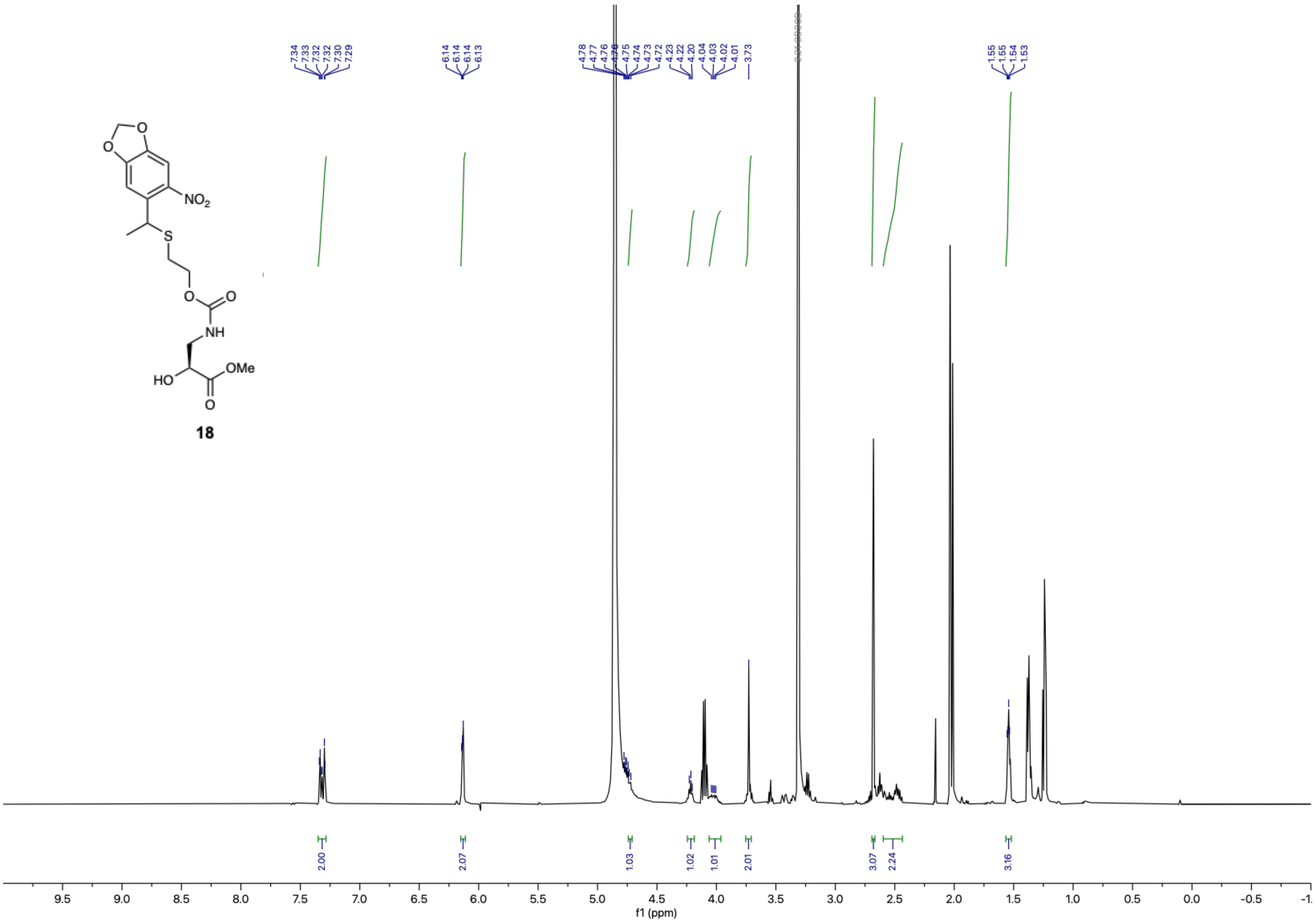

18

$^1$H NMR (600 MHz, methanol-$d_4$) – Standard view **(4)**

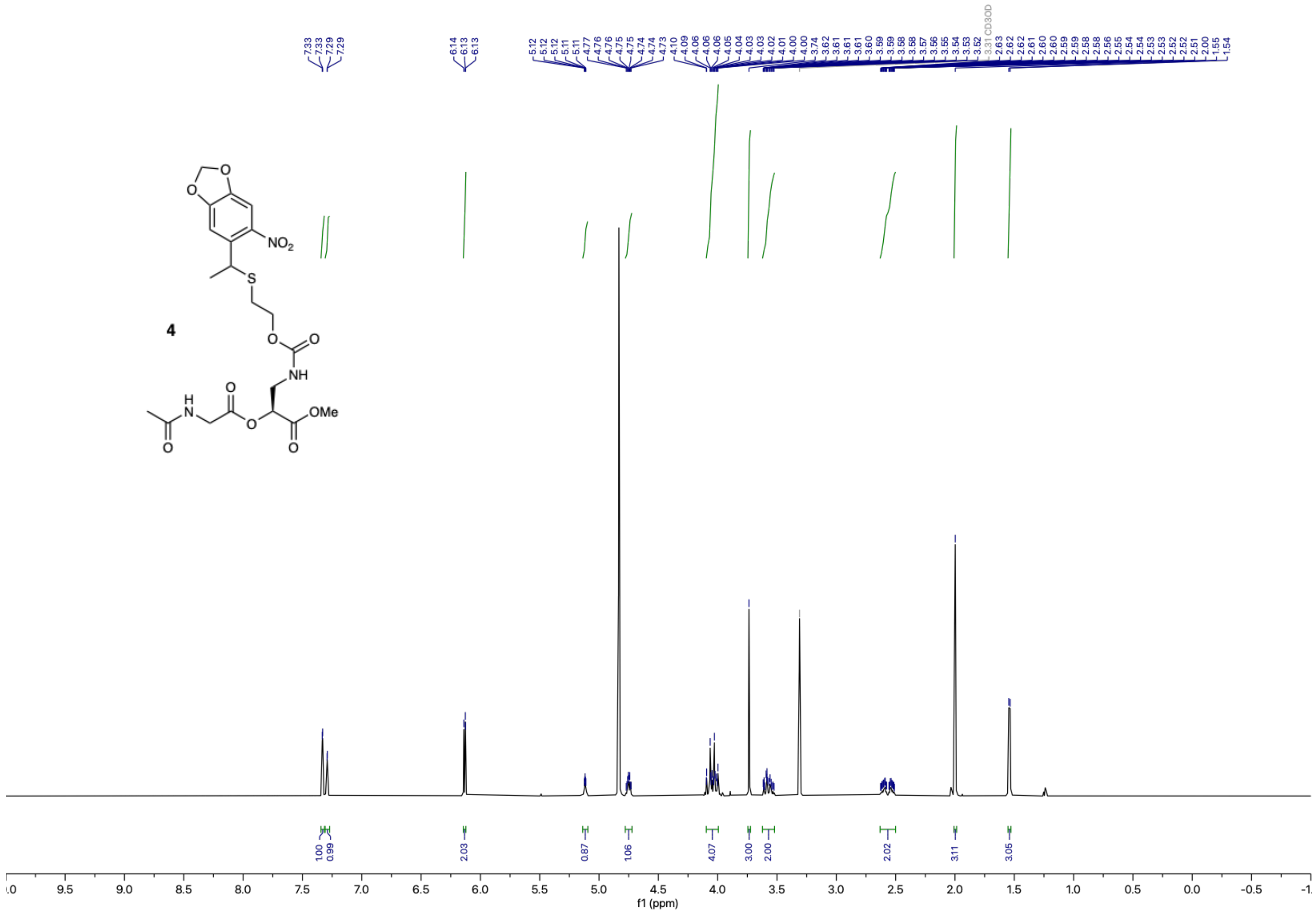

$^{13}C\{^1H\}$ NMR (151 MHz, methanol-$d_4$) – Standard view **(4)**

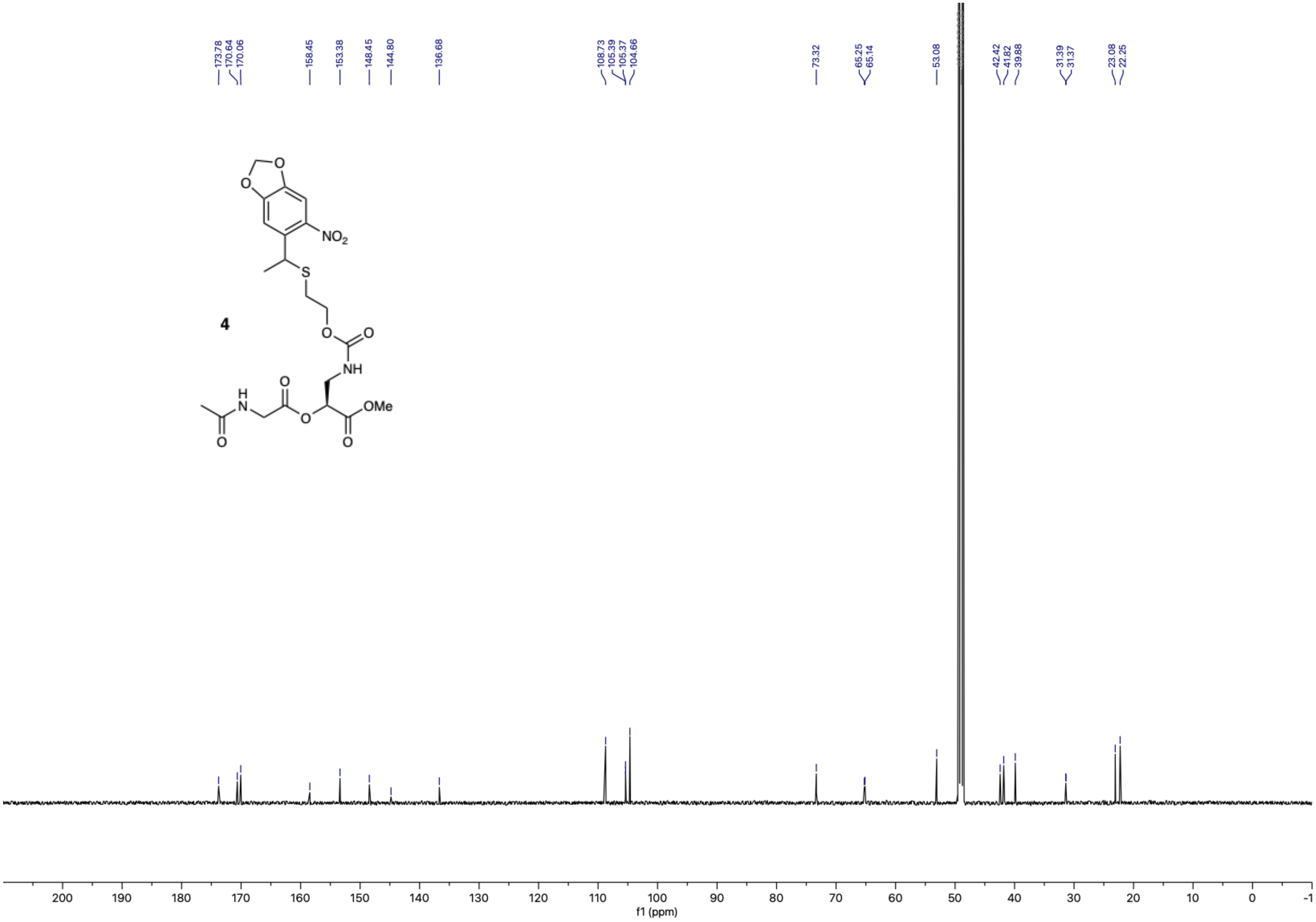

$^{1}H$ NMR (400 MHz, $CDCl_3$) – Extended view **((±)-20)**

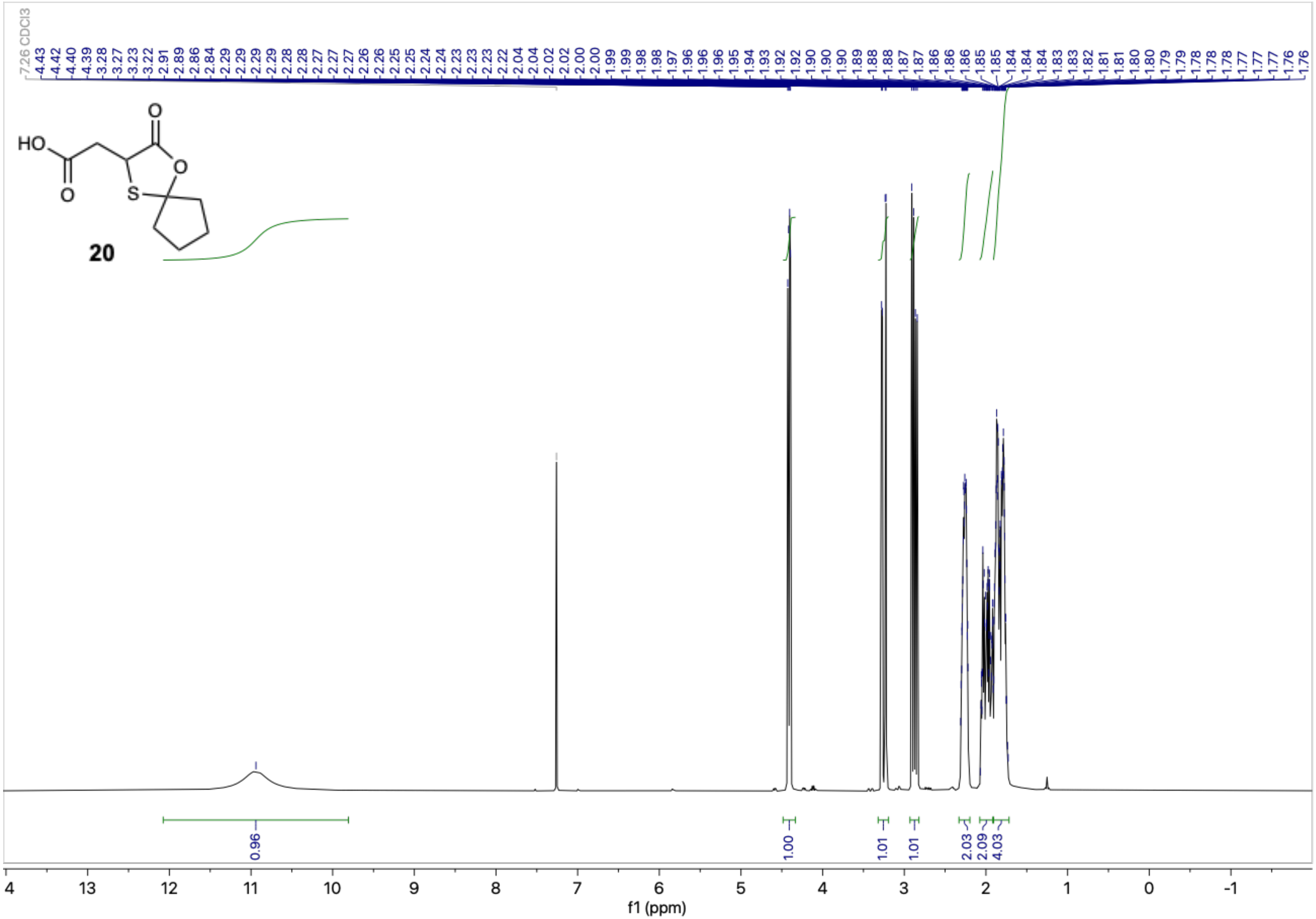

[1]H NMR (600 MHz, $CDCl_3$) – Standard view **((±)-21)**

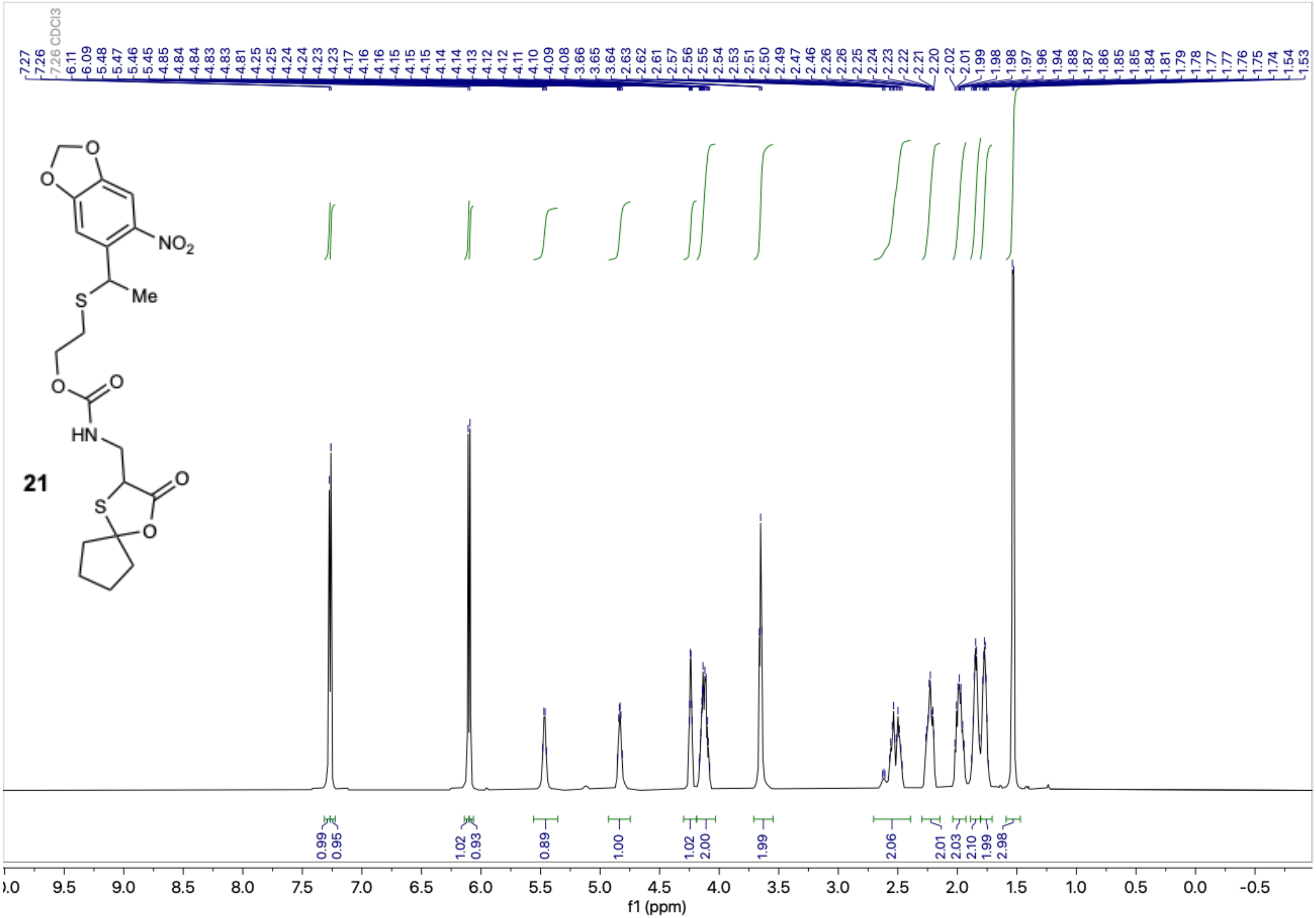

$^1H$-$^1H$ COSY ($CDCl_3$) **((±)-21)**

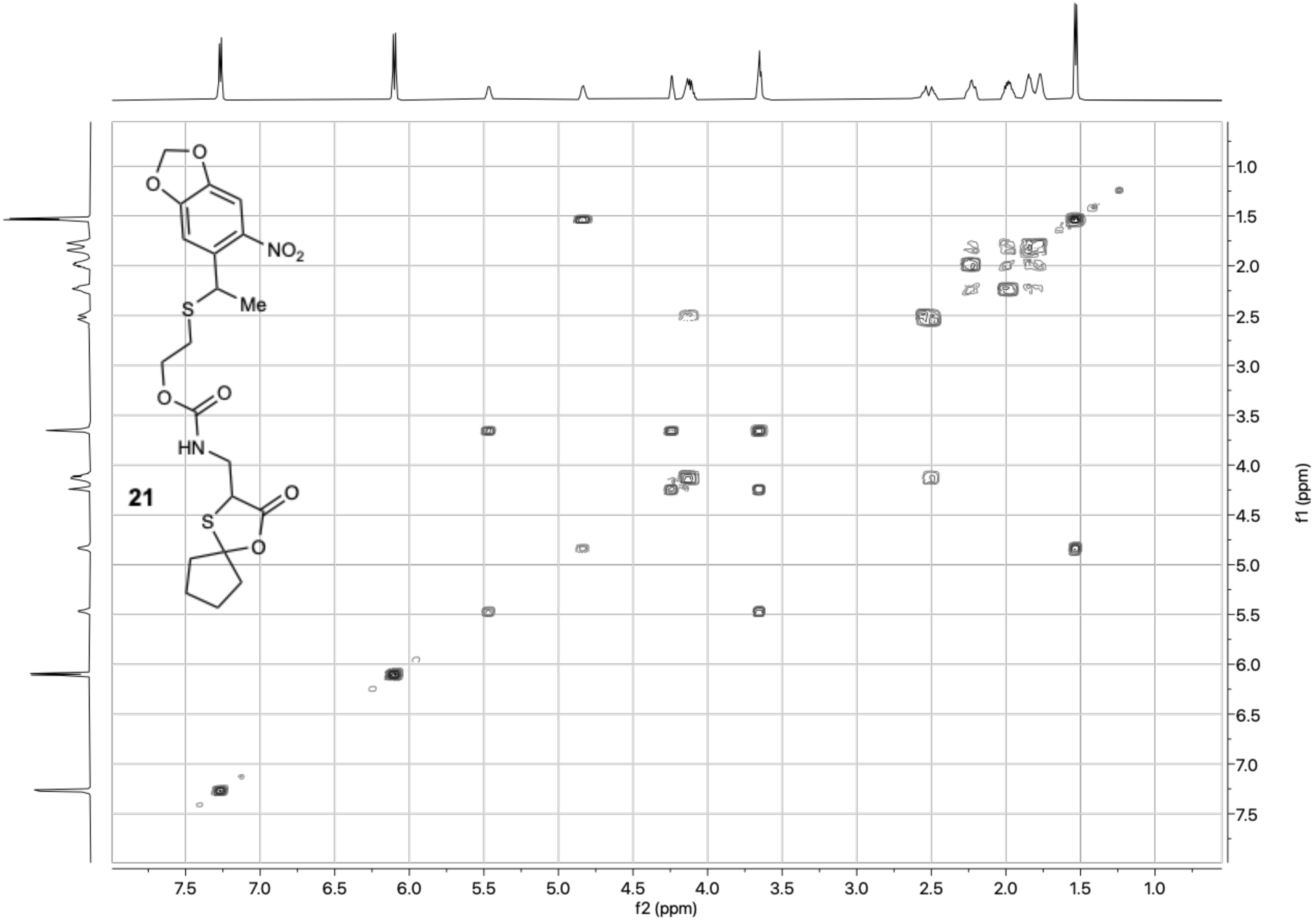

$^{13}C\{^1H\}$ NMR (151 MHz, $CDCl_3$) – Standard view **((±)-21)**

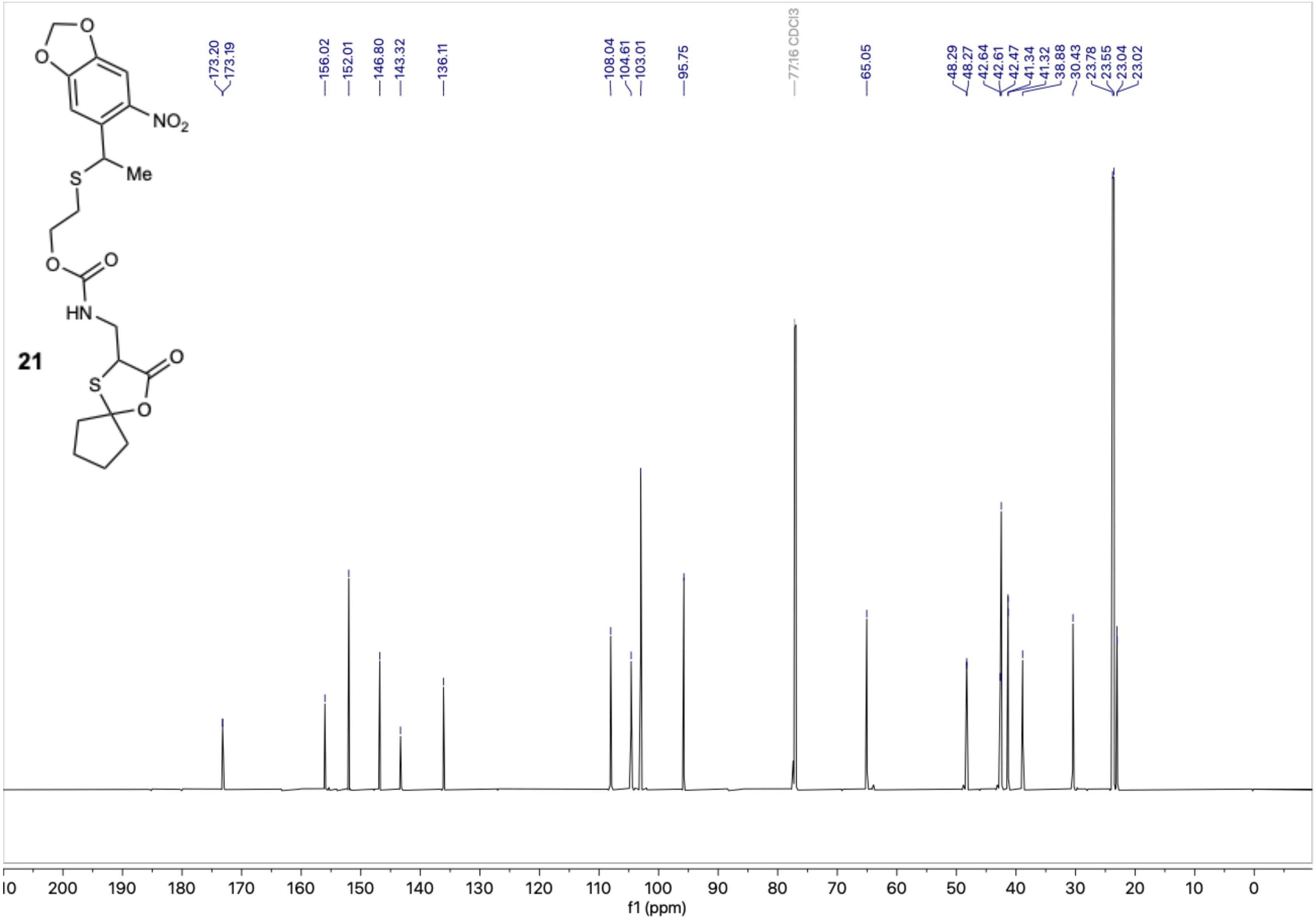

Multiplicity-Edited $^{1}H$-$^{13}C$ HSQC ($CDCl_3$) **((±)-21)**

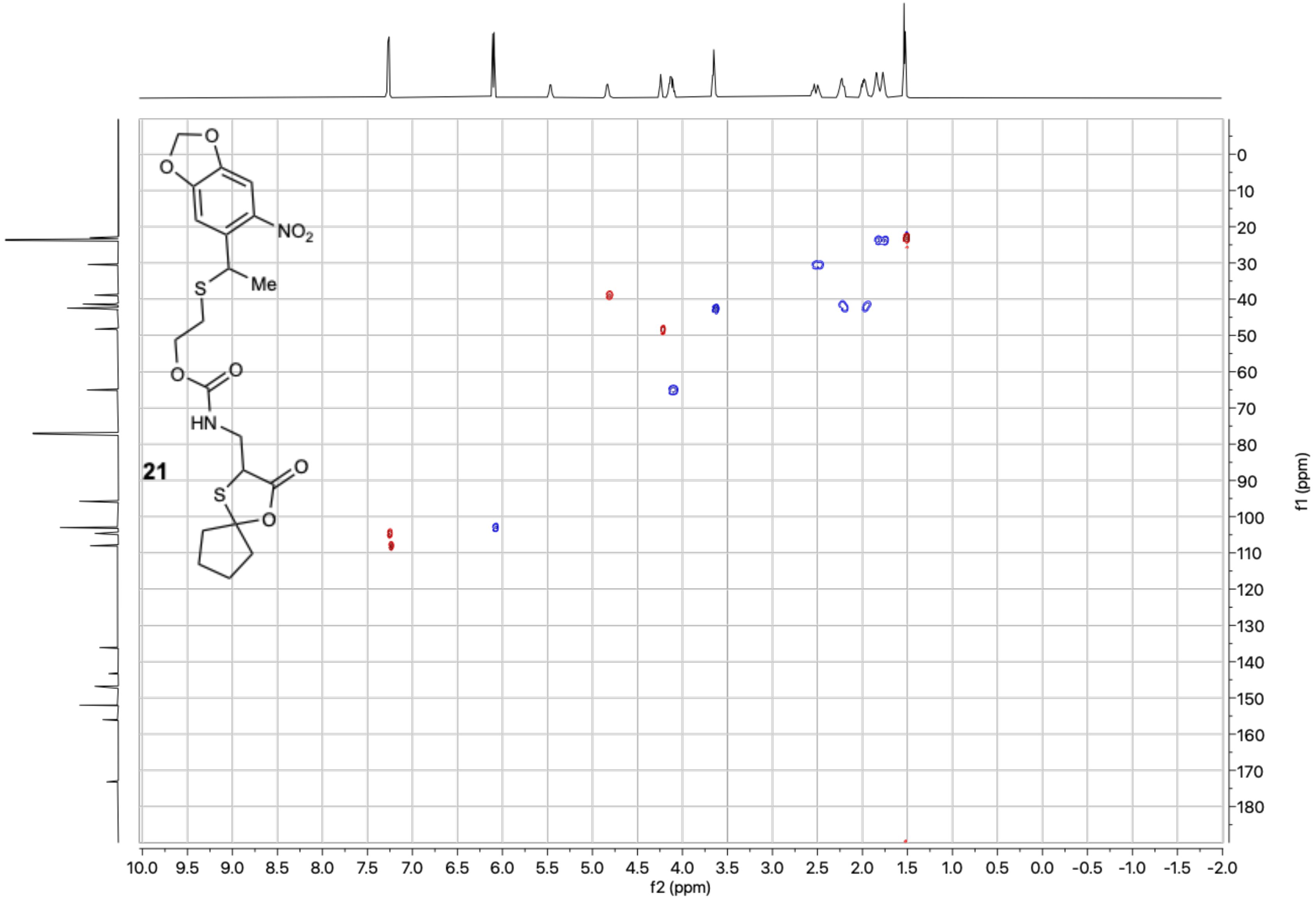

$^{1}H$ NMR (400 MHz, $CDCl_3$) – Standard view, crude **((±)-8)**

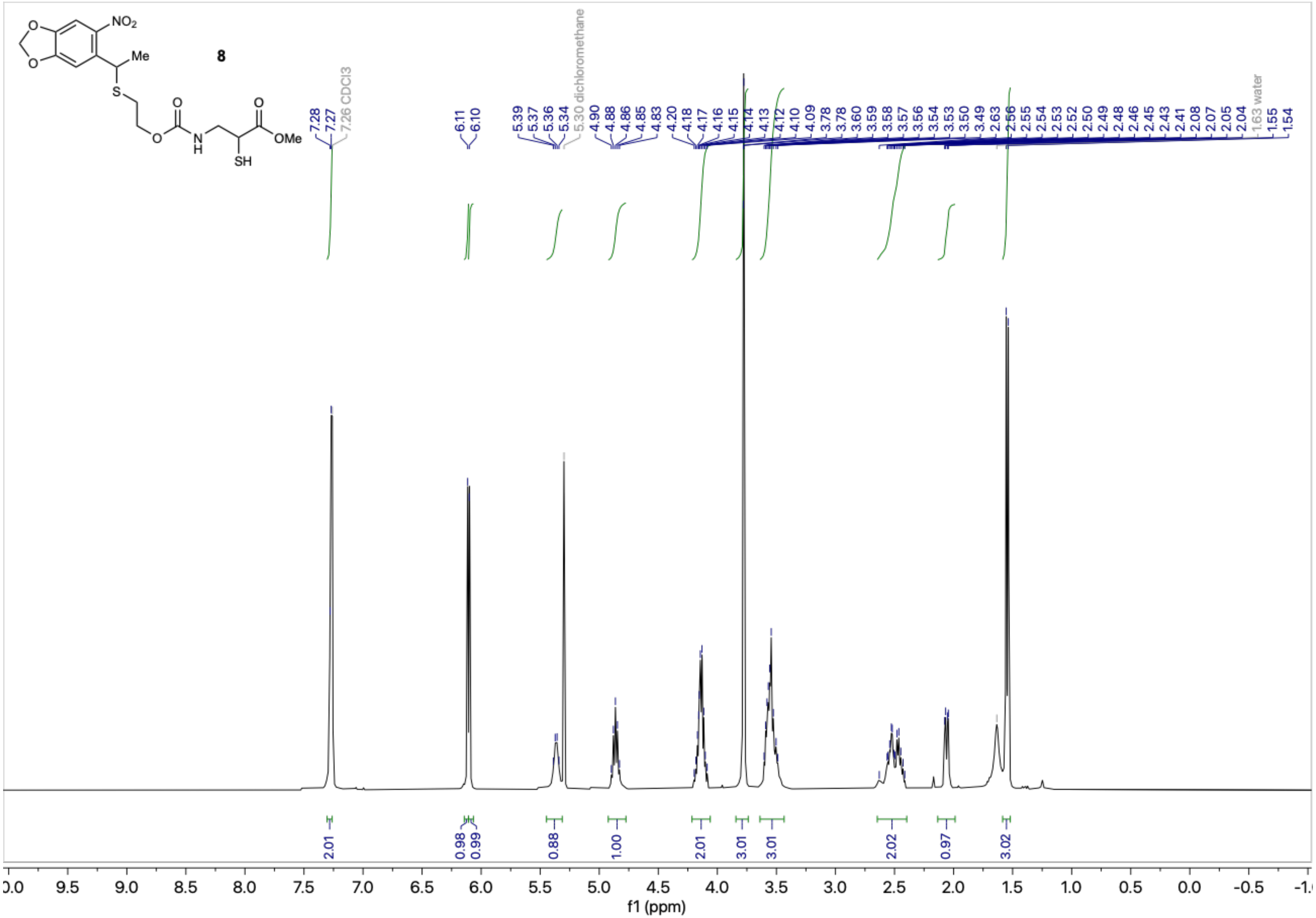

$^{1}H$ NMR (500 MHz, $CDCl_3$) – Standard view **((±)-5)**

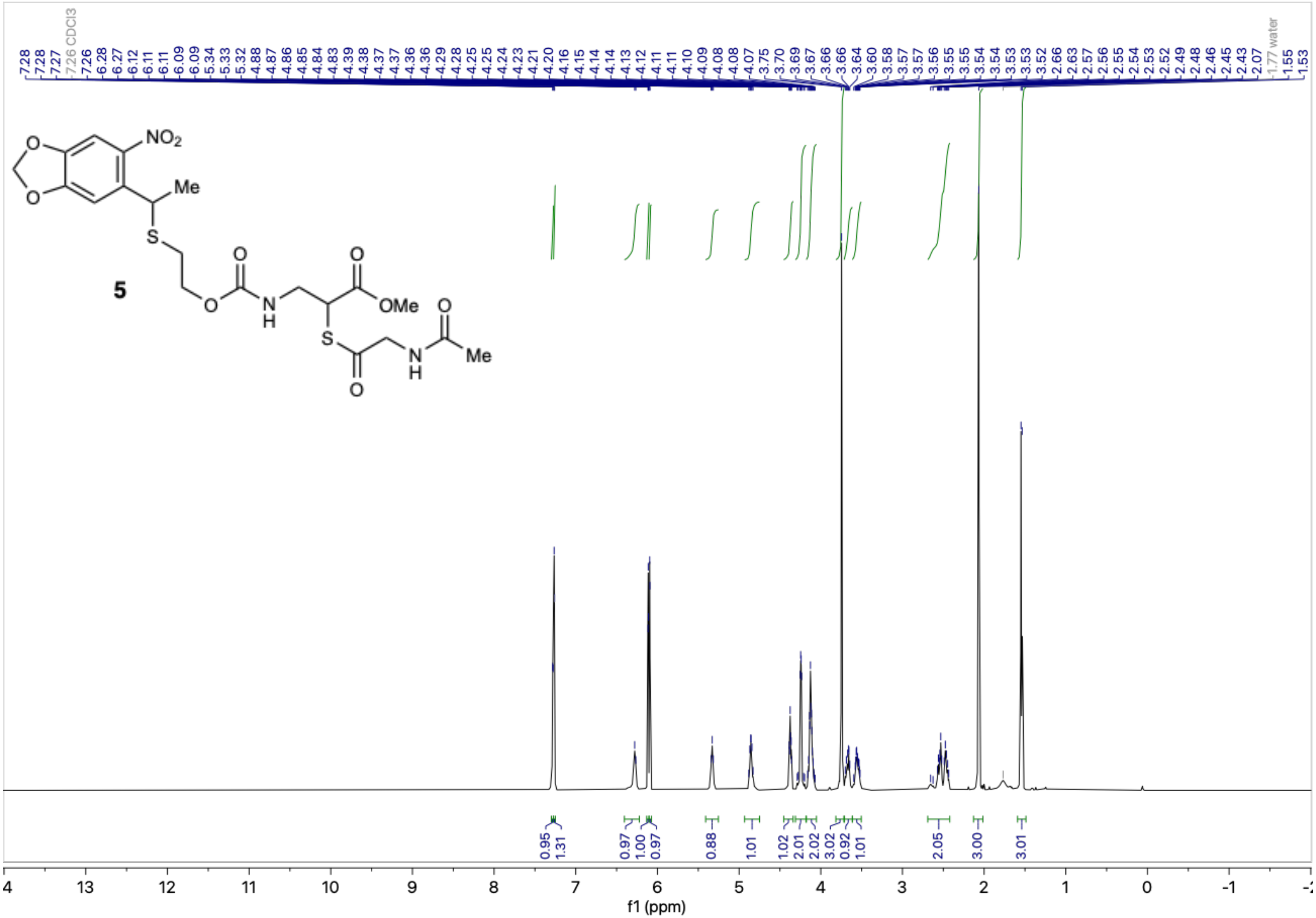

[1]H-[1]H COSY ($CDCl_3$) **((±)-5)**

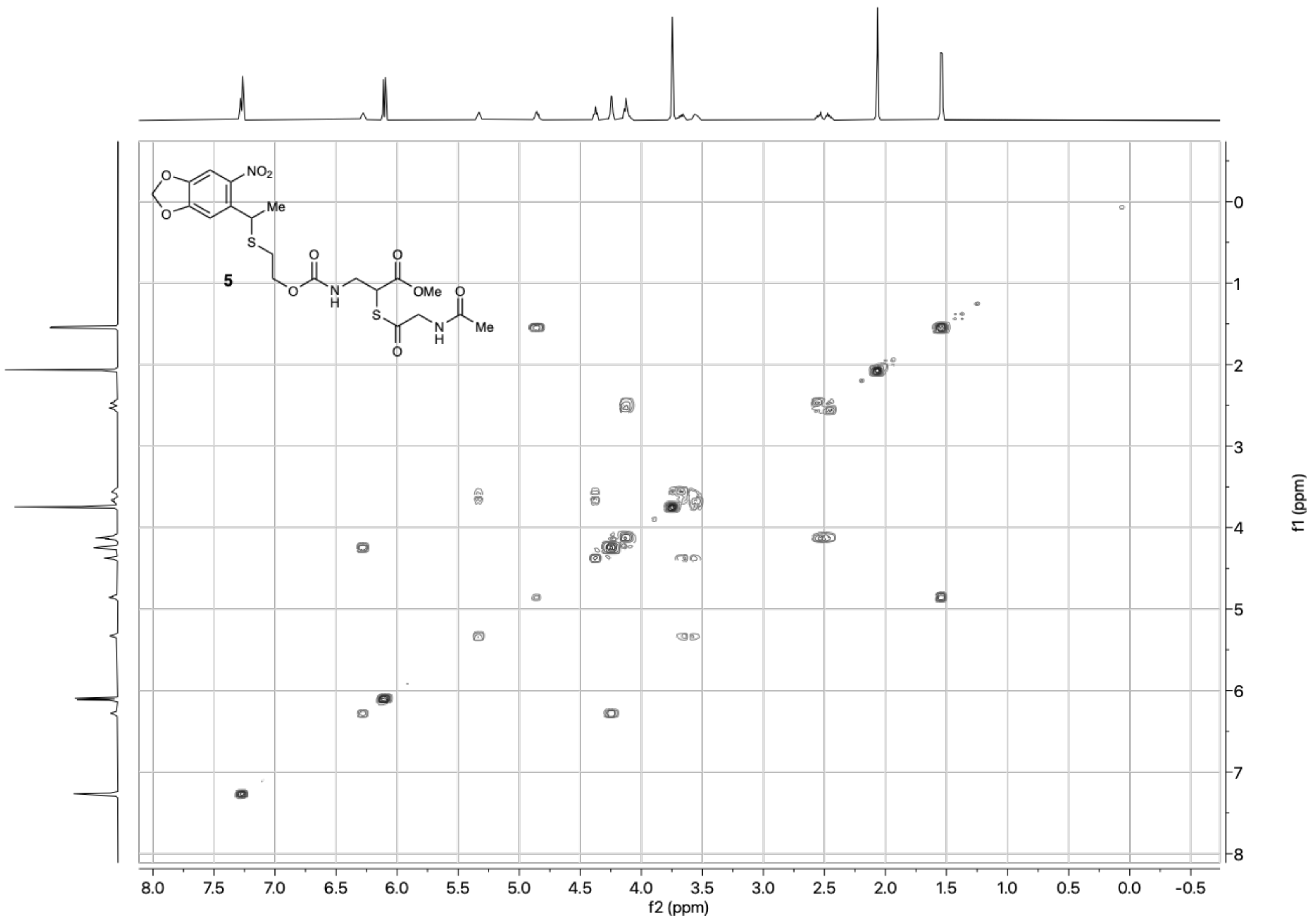

$^{13}C\{^1H\}$ NMR (126 MHz, $CDCl_3$) – Standard view **((±)-5)**

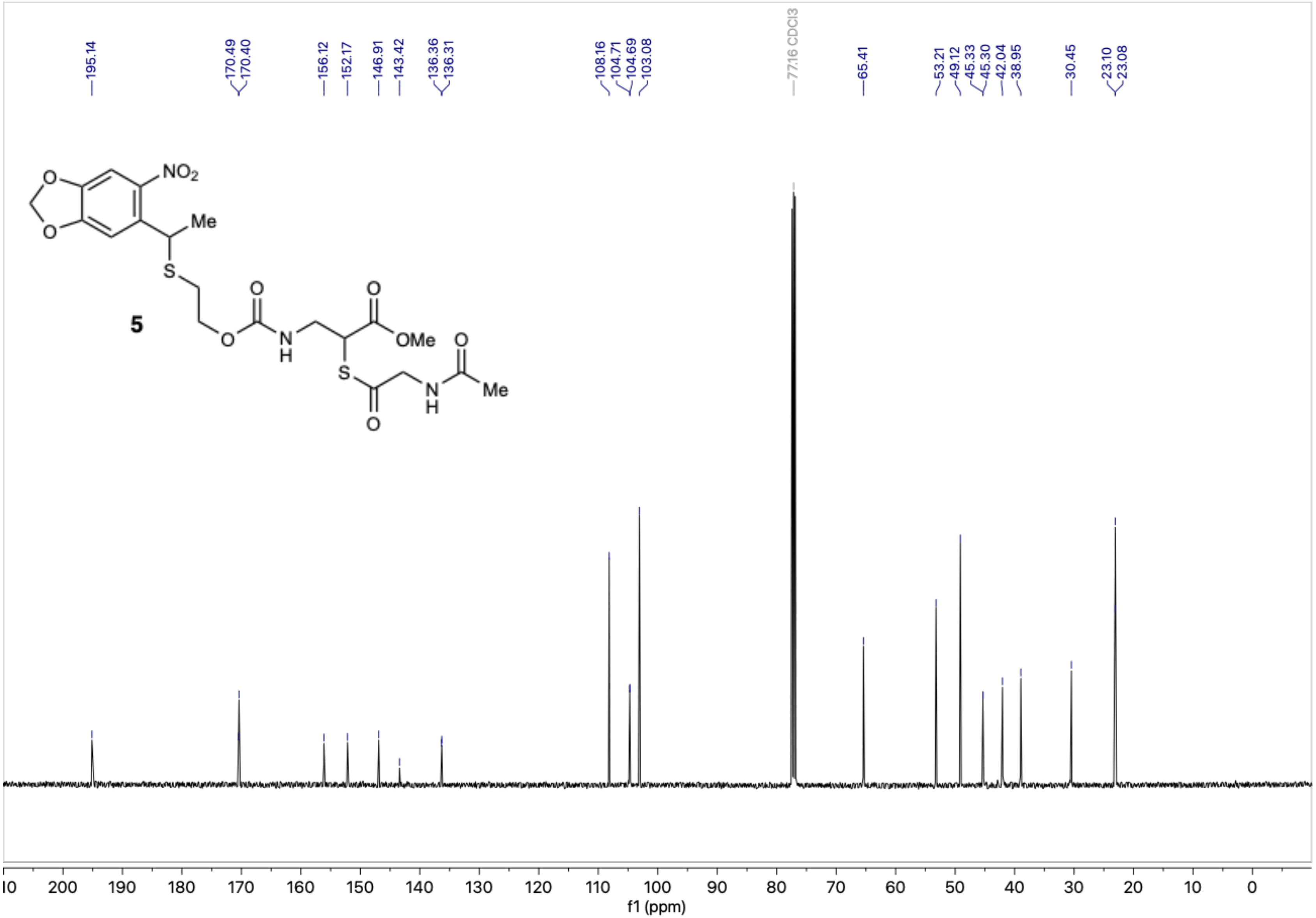

Multiplicity-Edited $^1$H-$^{13}$C HSQC ($CDCl_3$) – **((±)-5)**

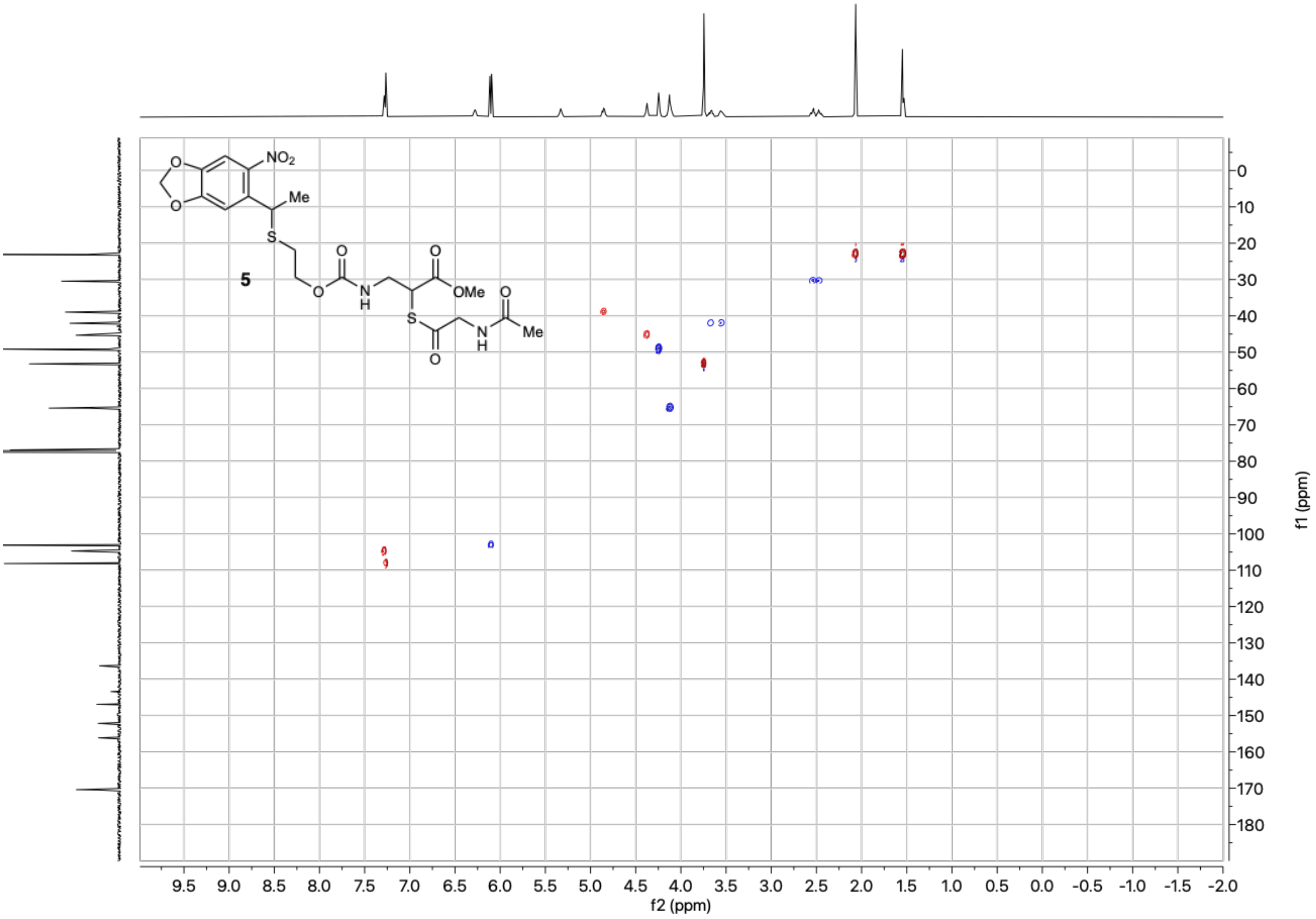

$^1$H NMR (600 MHz, methanol-$d_4$) – Standard view **(22)**

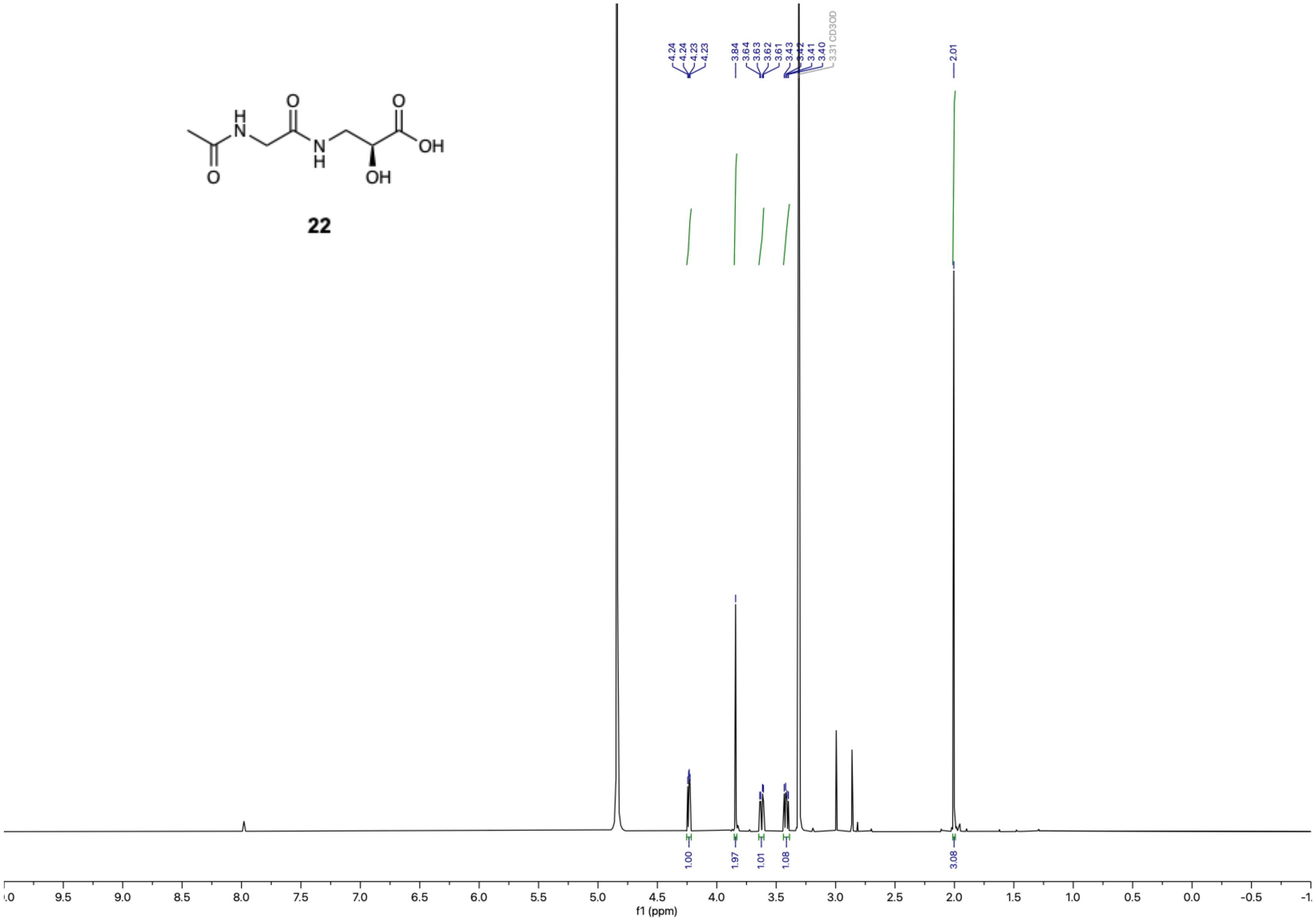

**22**

$^{13}C\{^{1}H\}$ NMR (151 MHz, methanol-$d_4$) – Standard view **(22)**

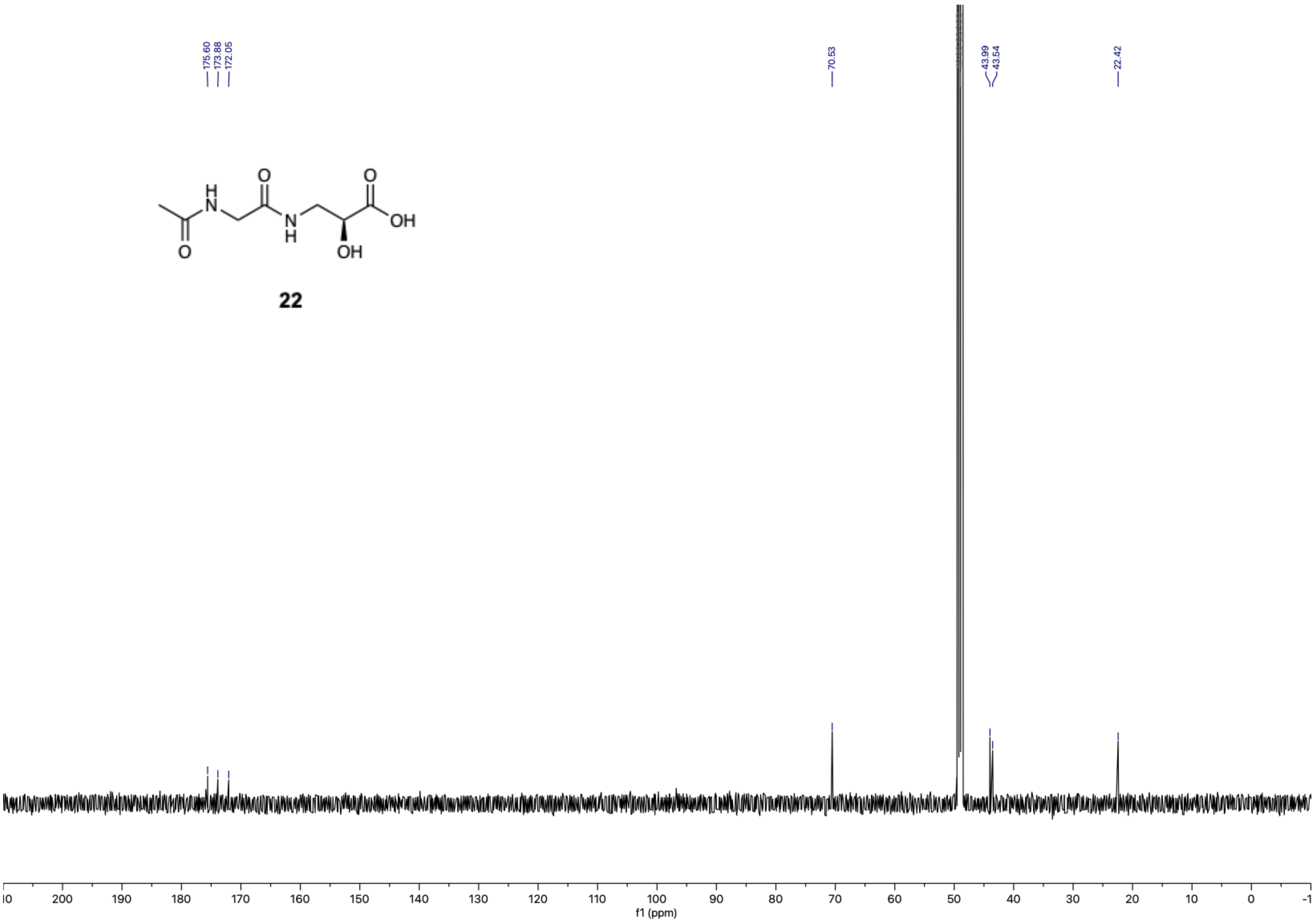

**22**

[1]H NMR (600 MHz, methanol-$d_4$) – Standard view **(6)**

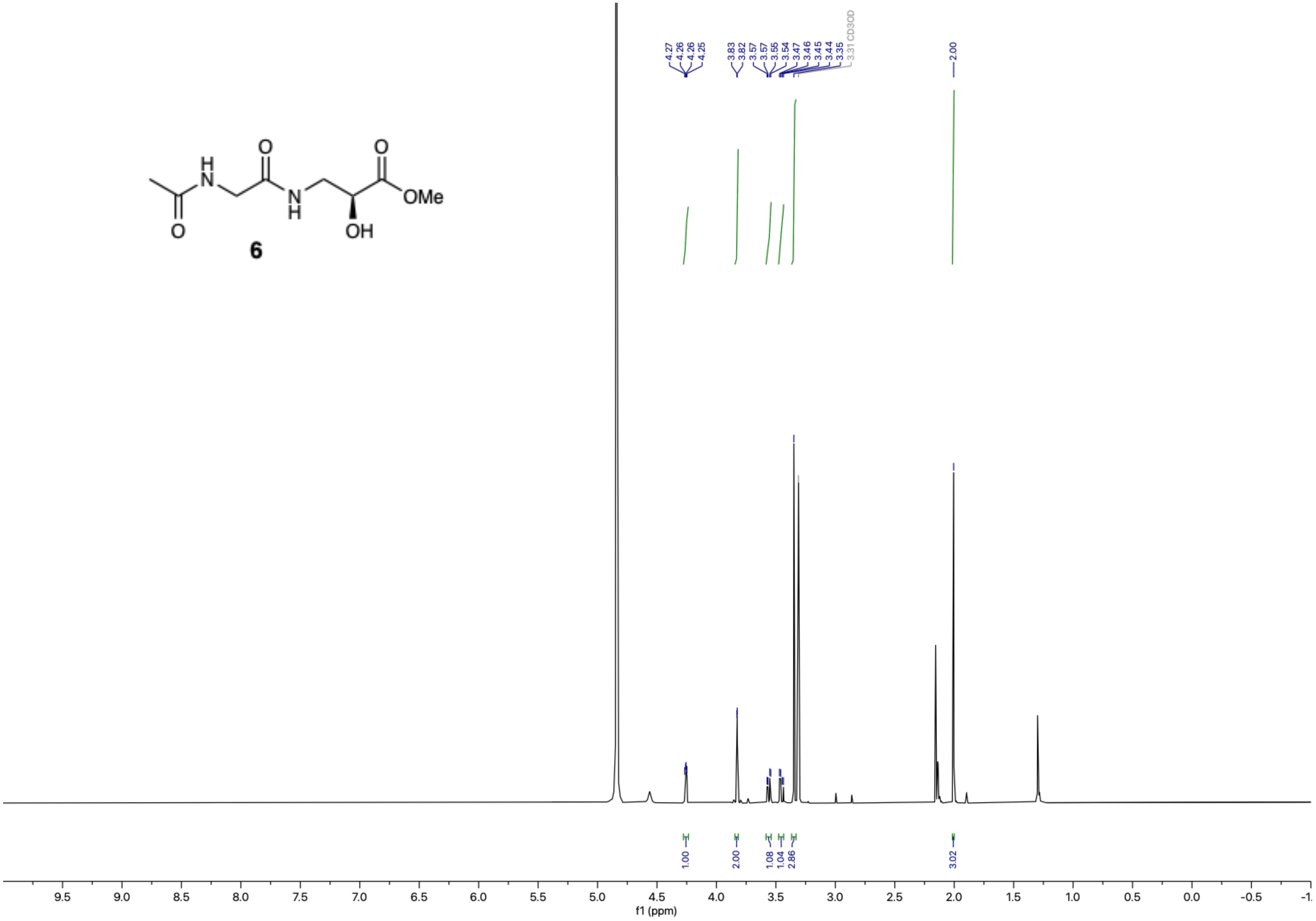

$^{13}C\{^{1}H\}$ NMR (151 MHz, methanol-$d_4$) – Standard view **(6)**

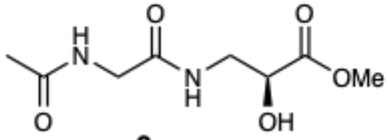

**6**

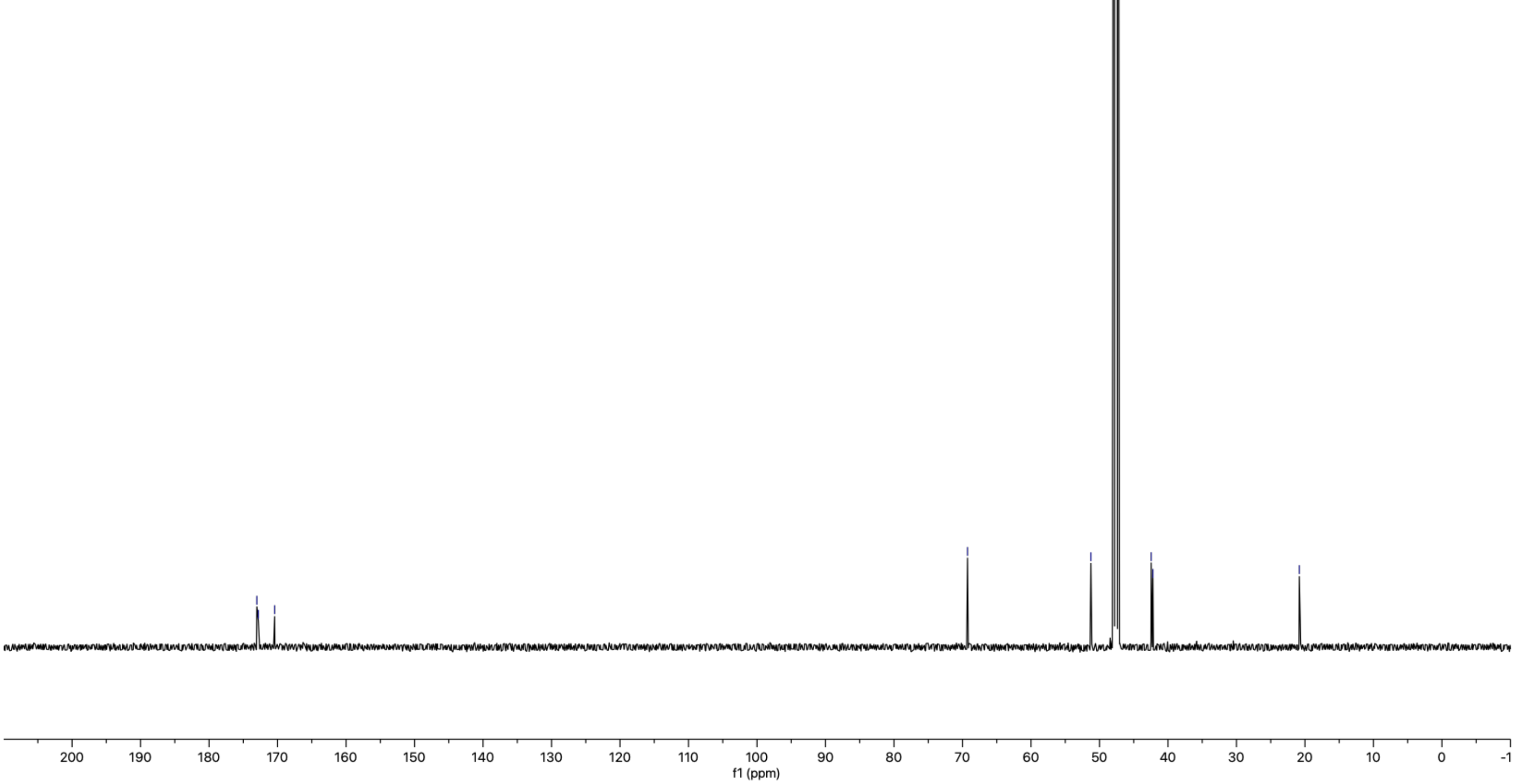

[1]H NMR (400 MHz, $CDCl_3$ + 1% v/v TMS) – extended view, crude **((±)-20,** as used in the preparation of **(±)-24)**

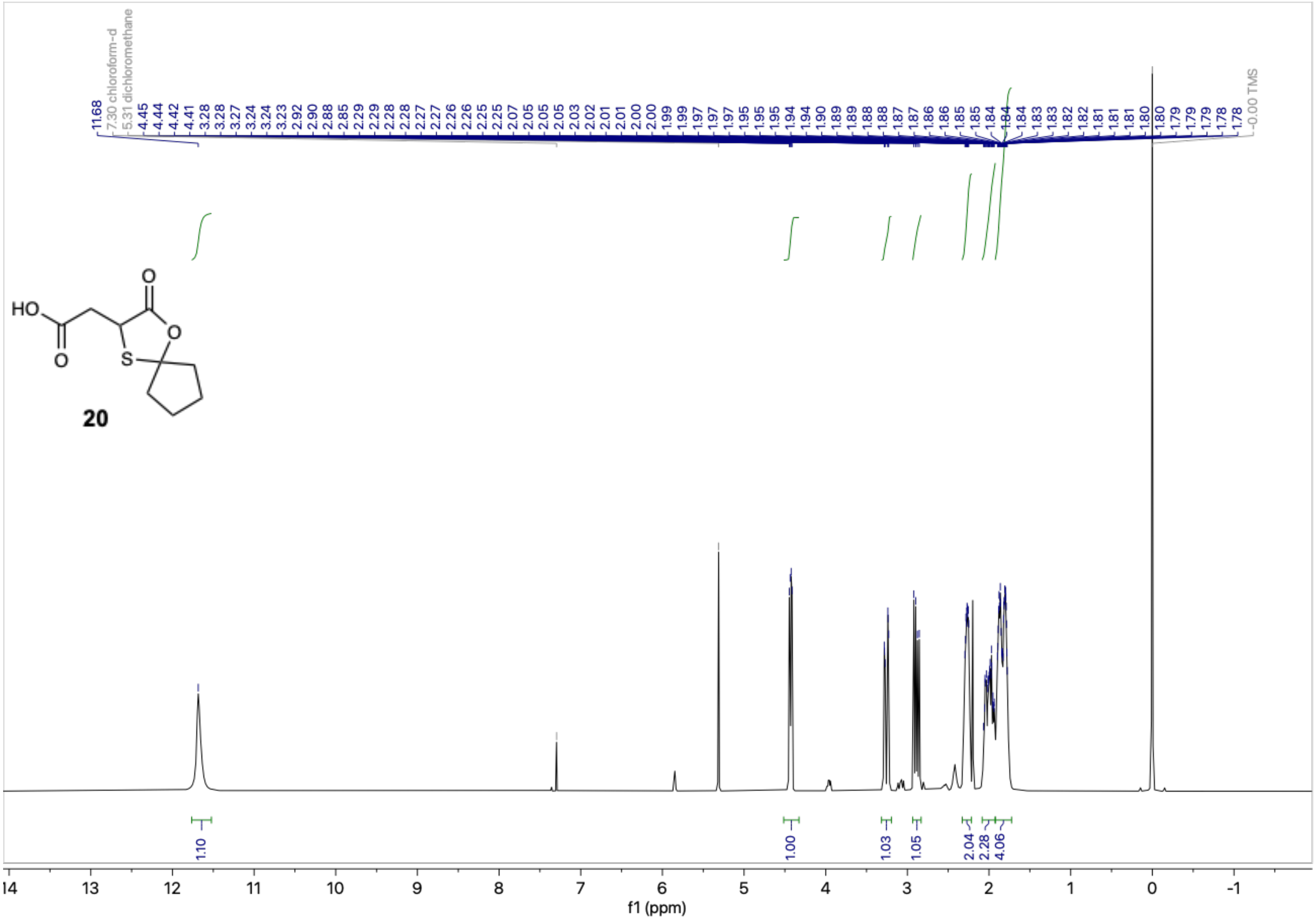

$^1$H NMR (400 MHz, DMSO-$d_6$) – standard view, crude **((±)-24)**

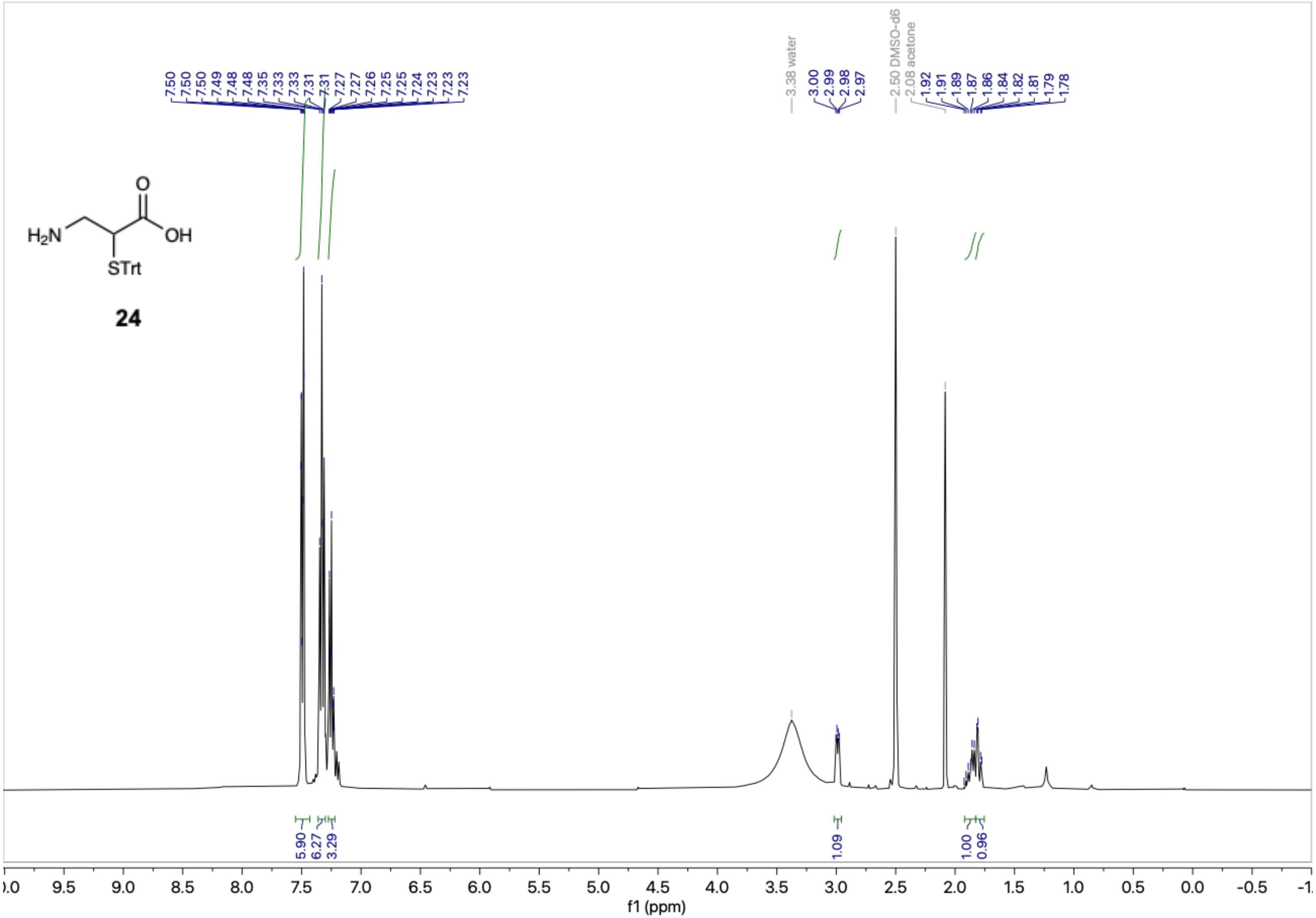

[1]H NMR (400 MHz, DMSO-$d_6$) – Extended view **((±)-25)**

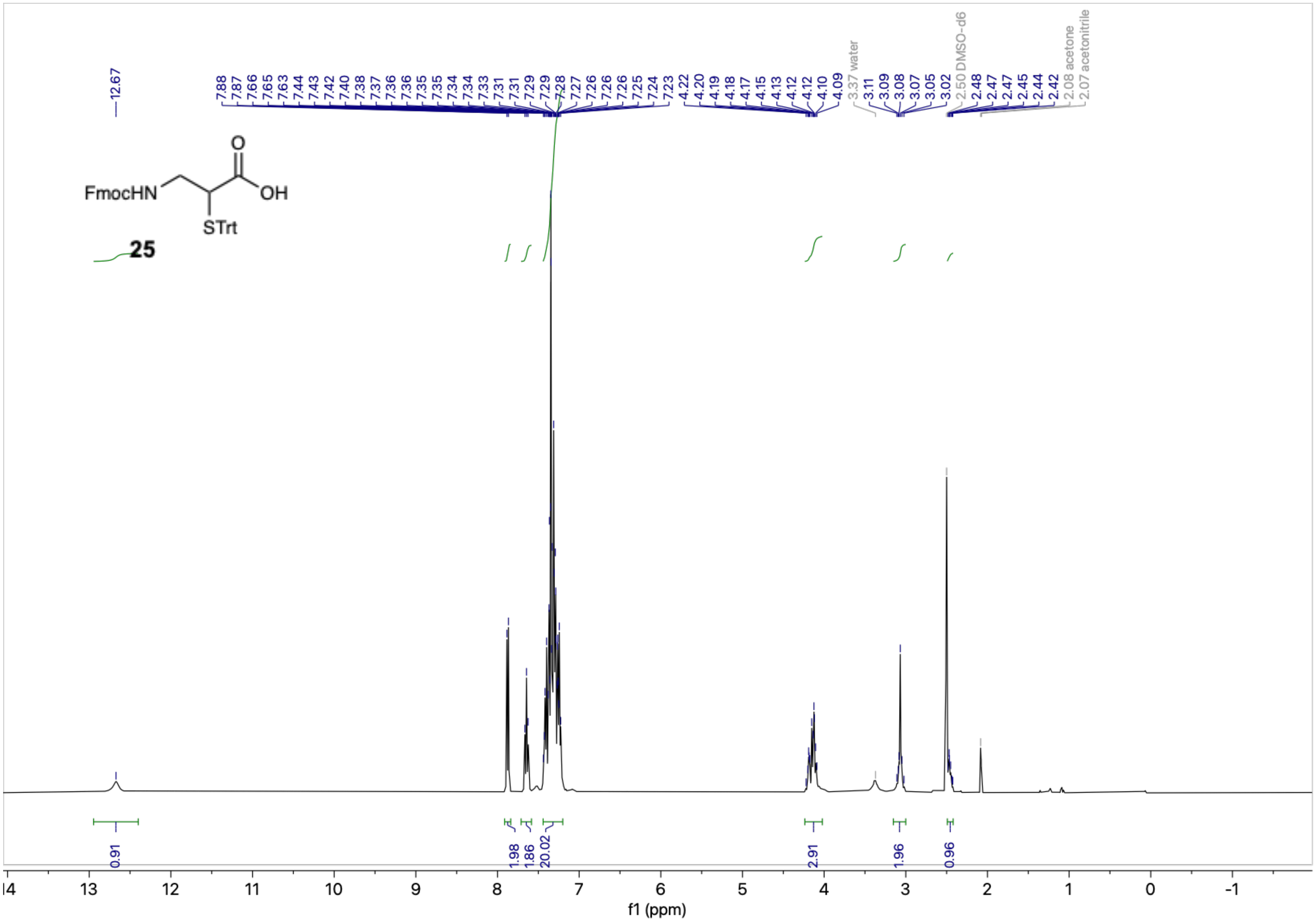

$^{1}H$-$^{1}H$ COSY (DMSO-$d_6$) (**(±)-25**)

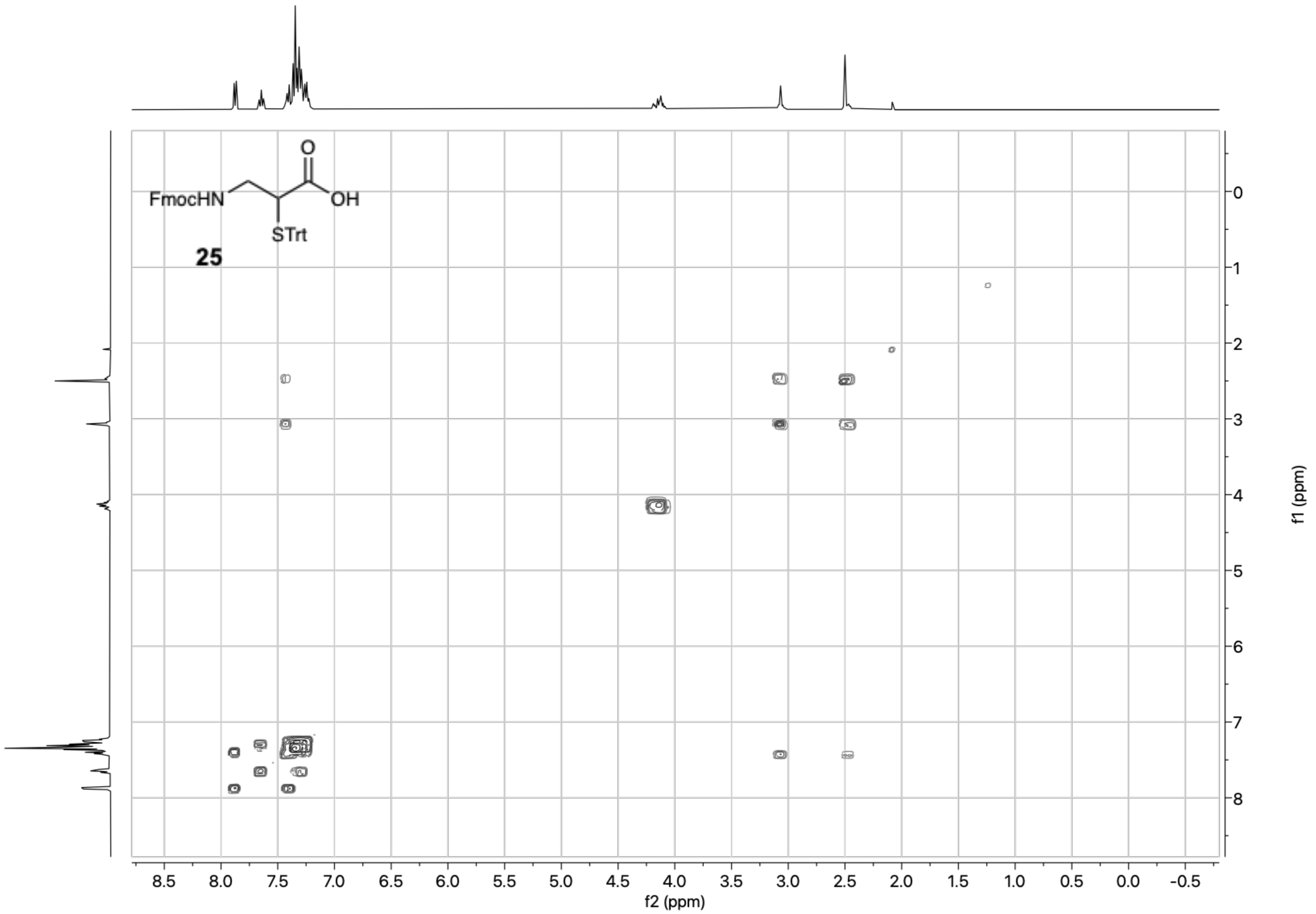

$^{13}C\{^{1}H\}$ NMR (151 MHz, DMSO-$d_6$) – Standard view **((±)-25)**

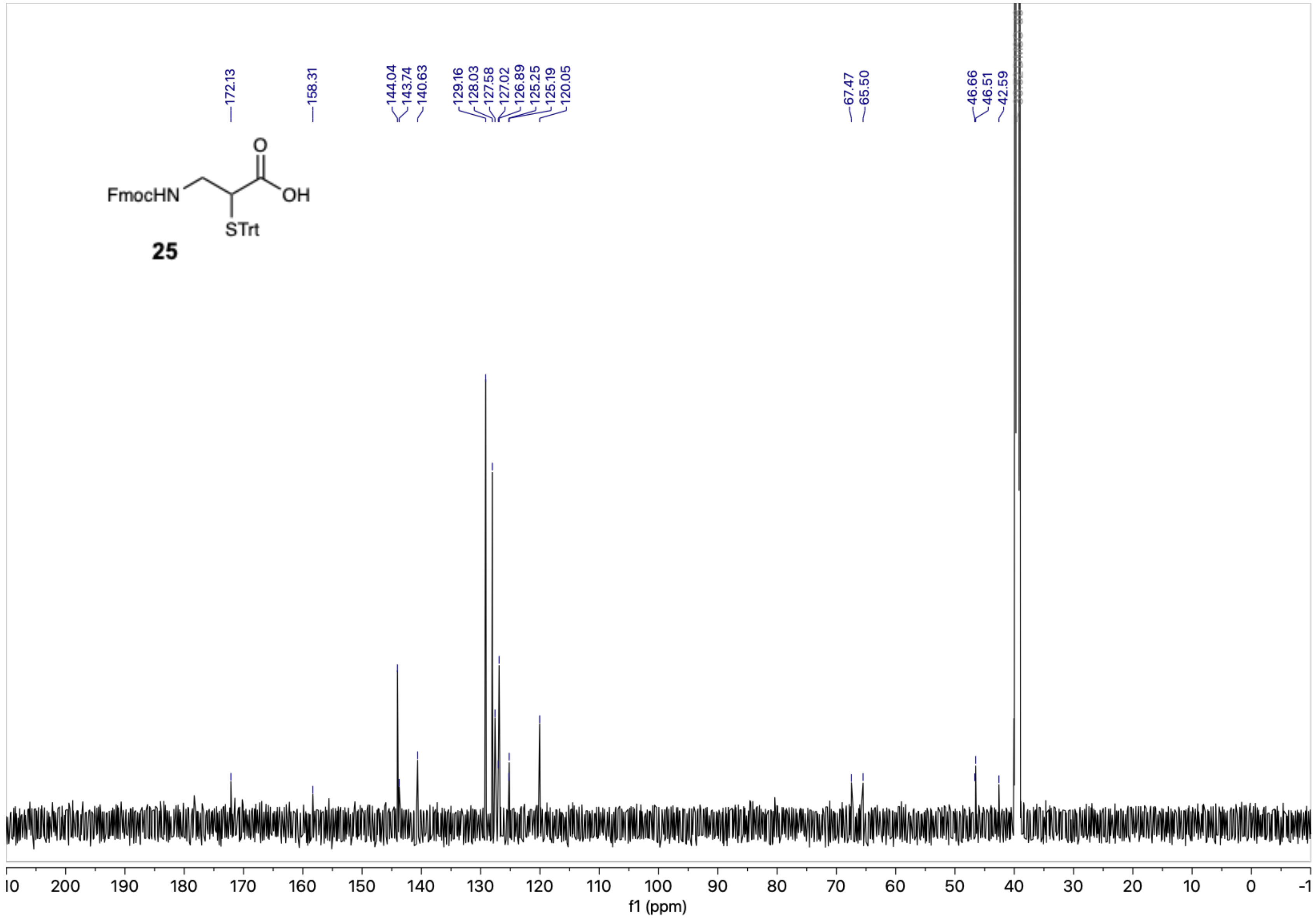

[1]H NMR (600 MHz, methanol-$d_4$) – Standard view **((±)-26)**

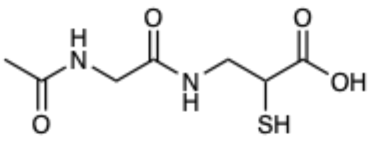

**26**

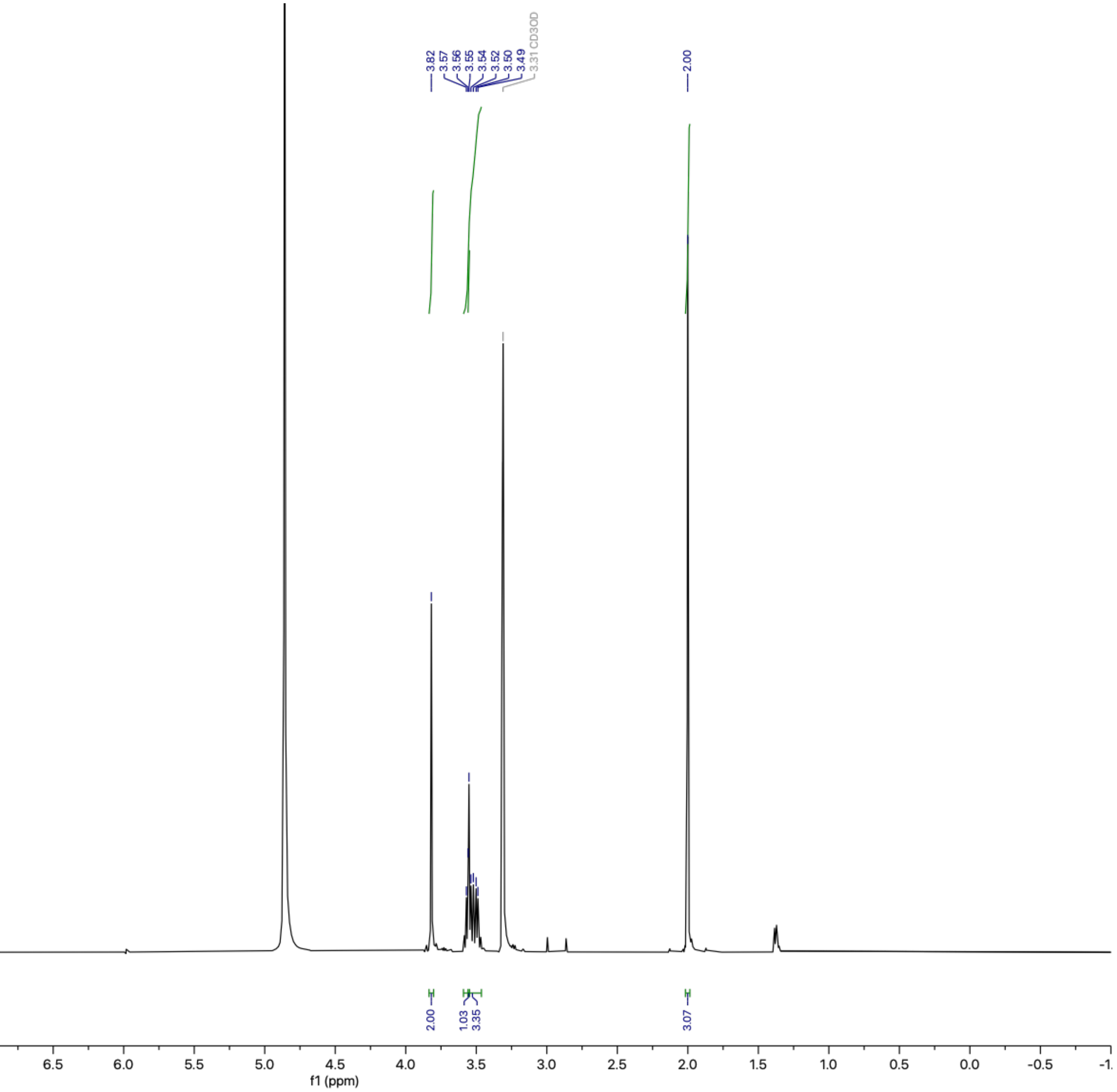

[1]H NMR (600 MHz, methanol-$d_4$) – Standard view **((±)-7)**

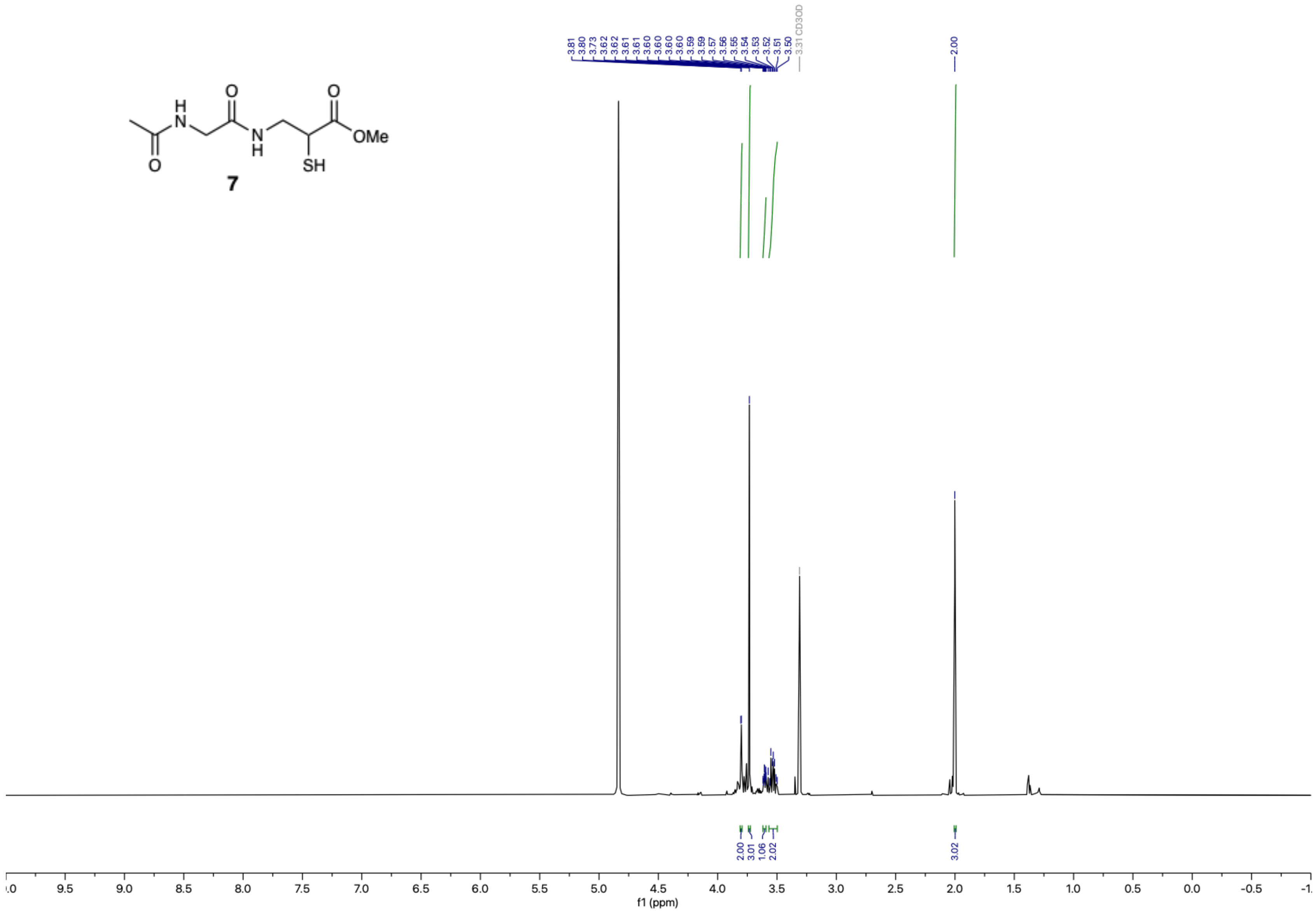

$^{13}C\{^1H\}$ NMR (151 MHz, methanol-$d_4$) – Standard view **((±)-7)**

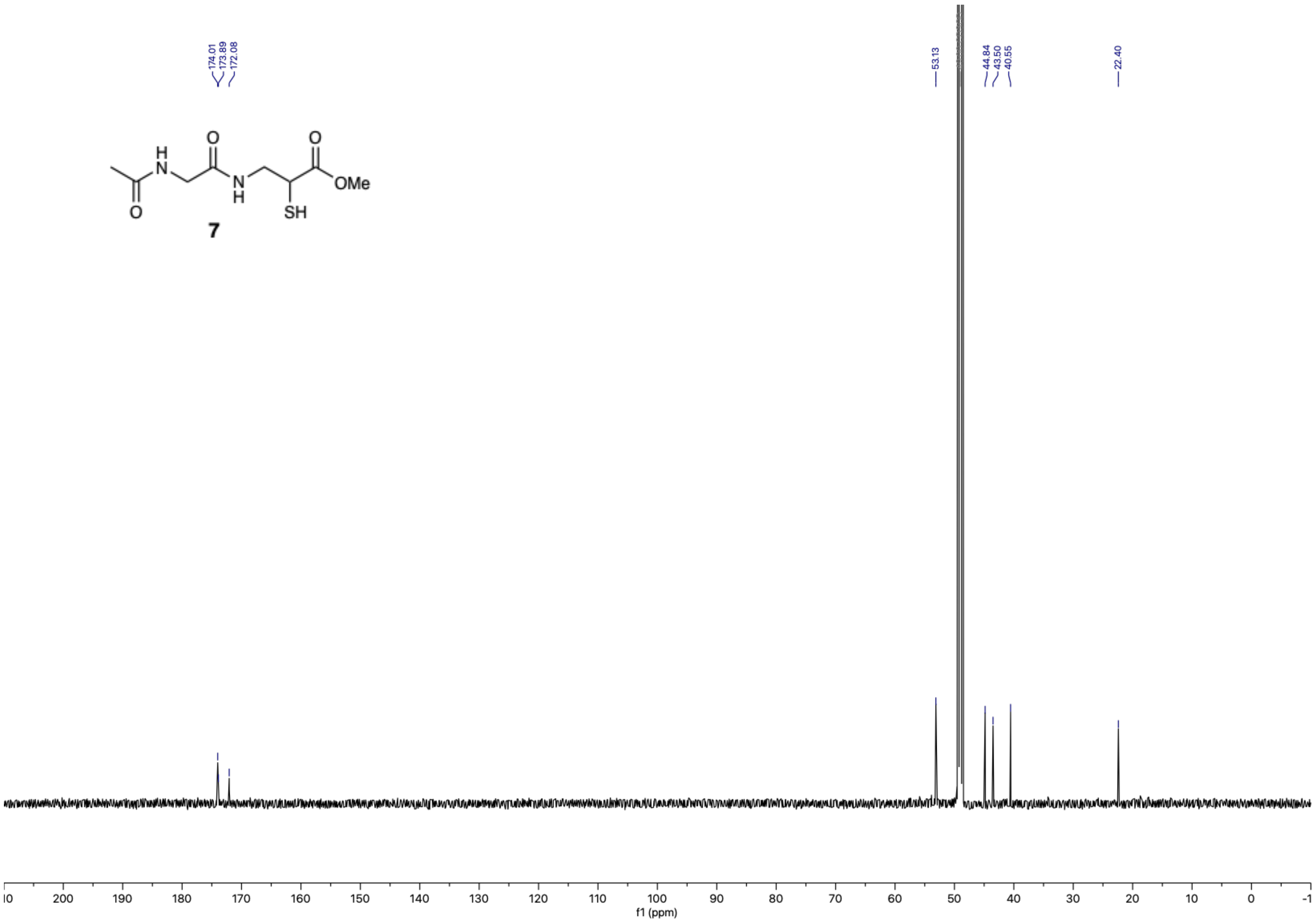

$^{1}$H NMR (500 MHz, methanol-$d_4$) – Standard view **(27)**

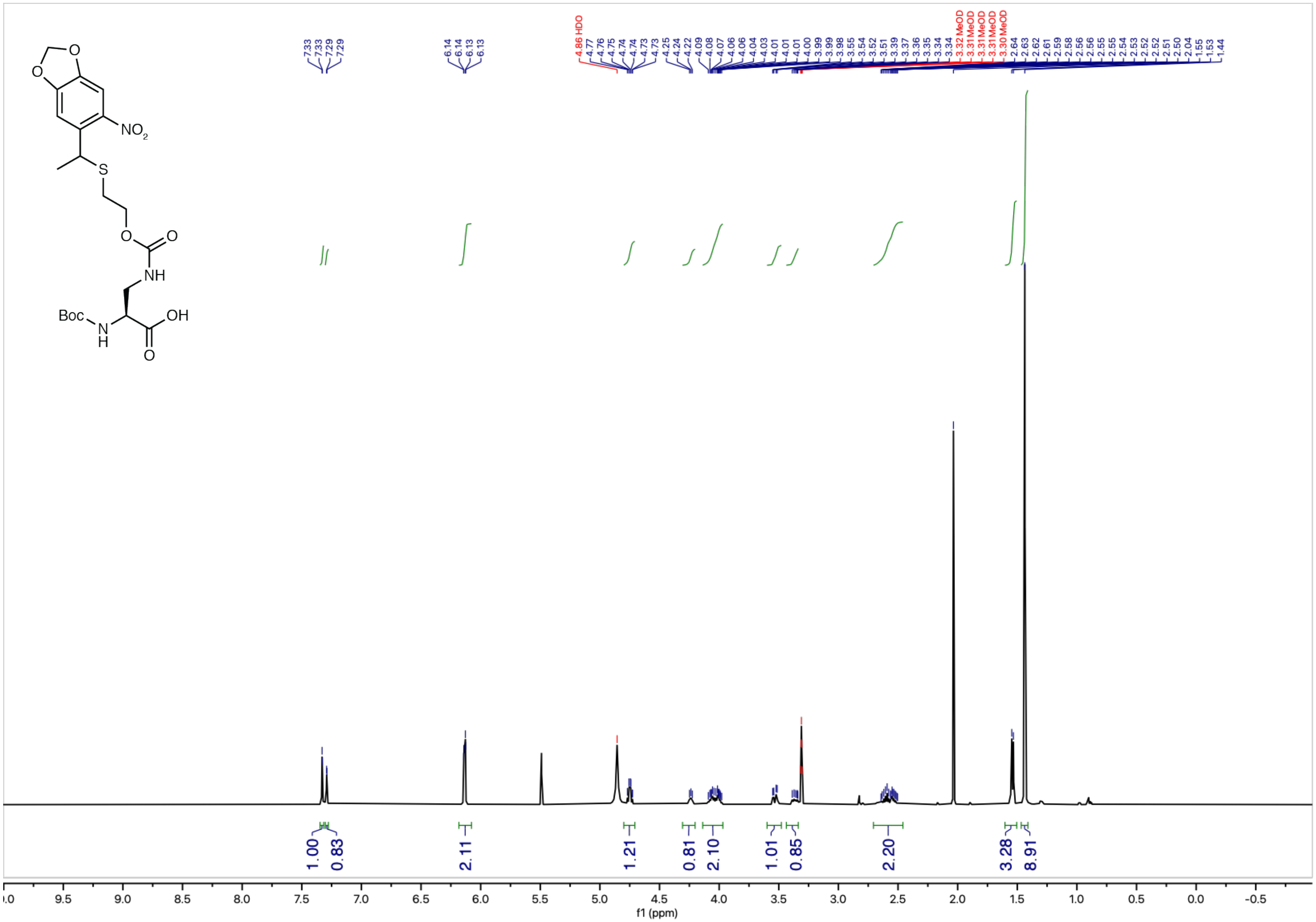

$^{13}C\{^{1}H\}$ NMR (126 MHz, methanol-$d_4$) – Standard view **(27)**

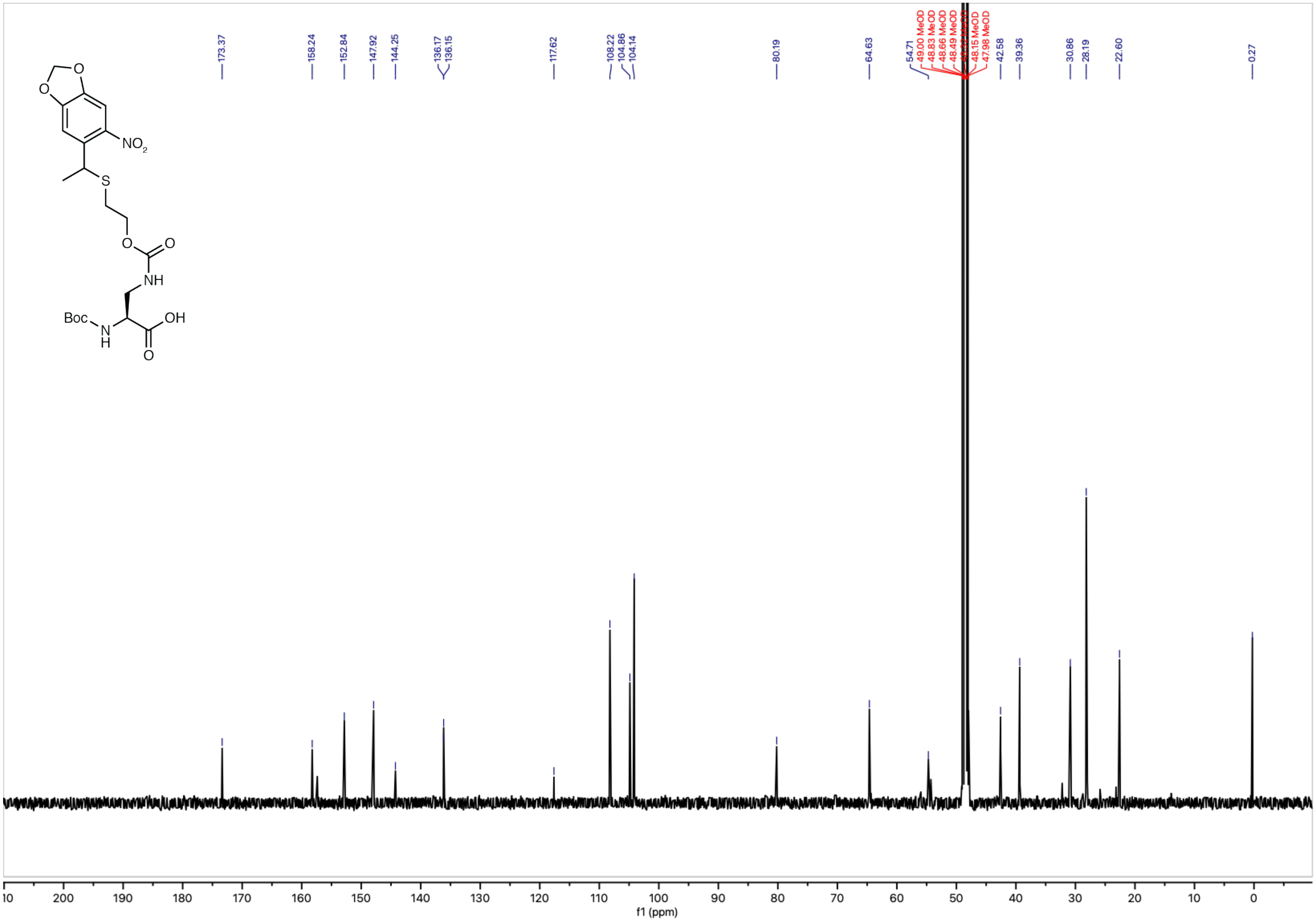

[1]H NMR (500 MHz, methanol-$d_4$) – Standard view **(1)**

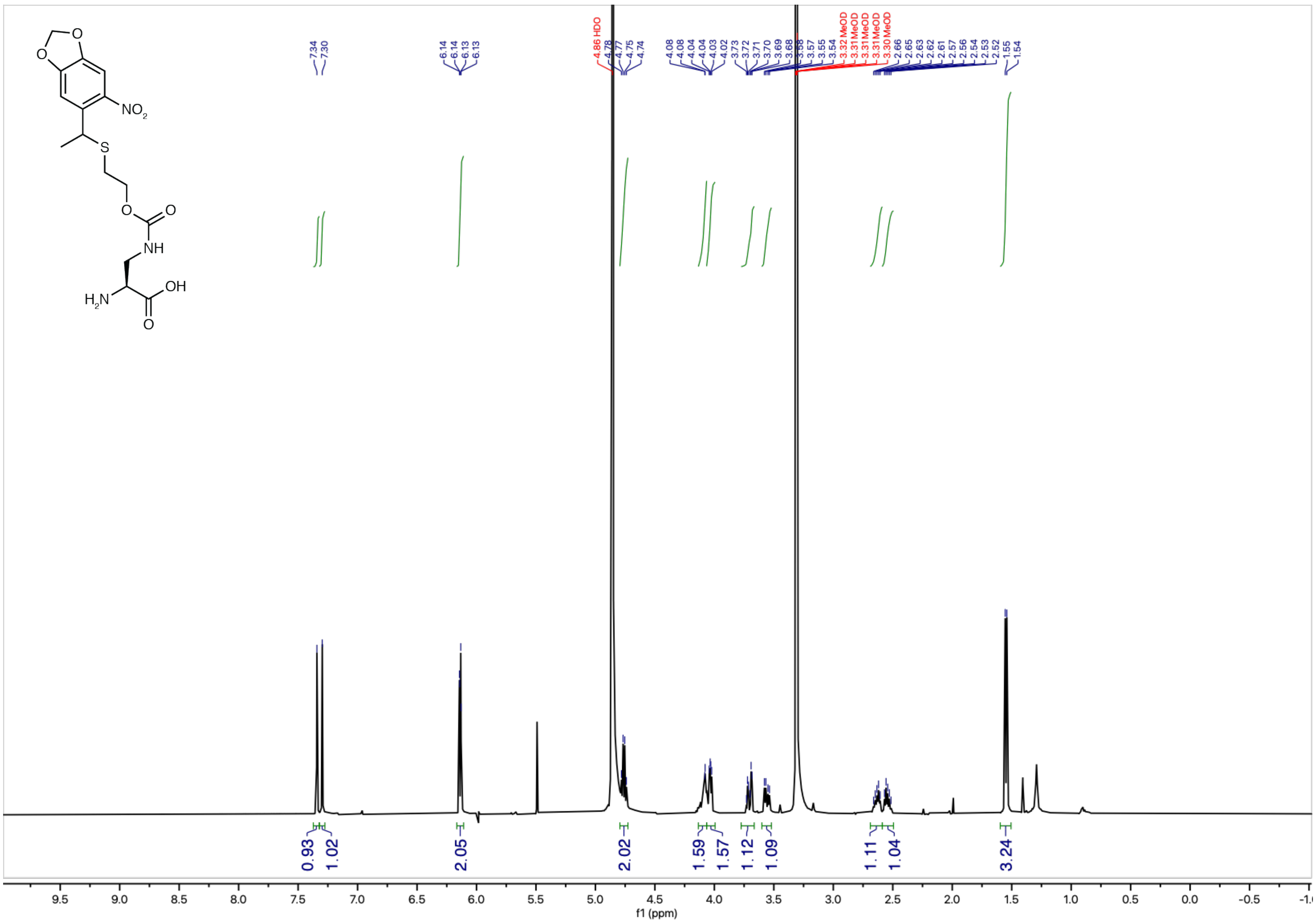

$^{13}C\{^{1}H\}$ NMR (126 MHz, methanol-$d_4$) – Standard view **(1)**

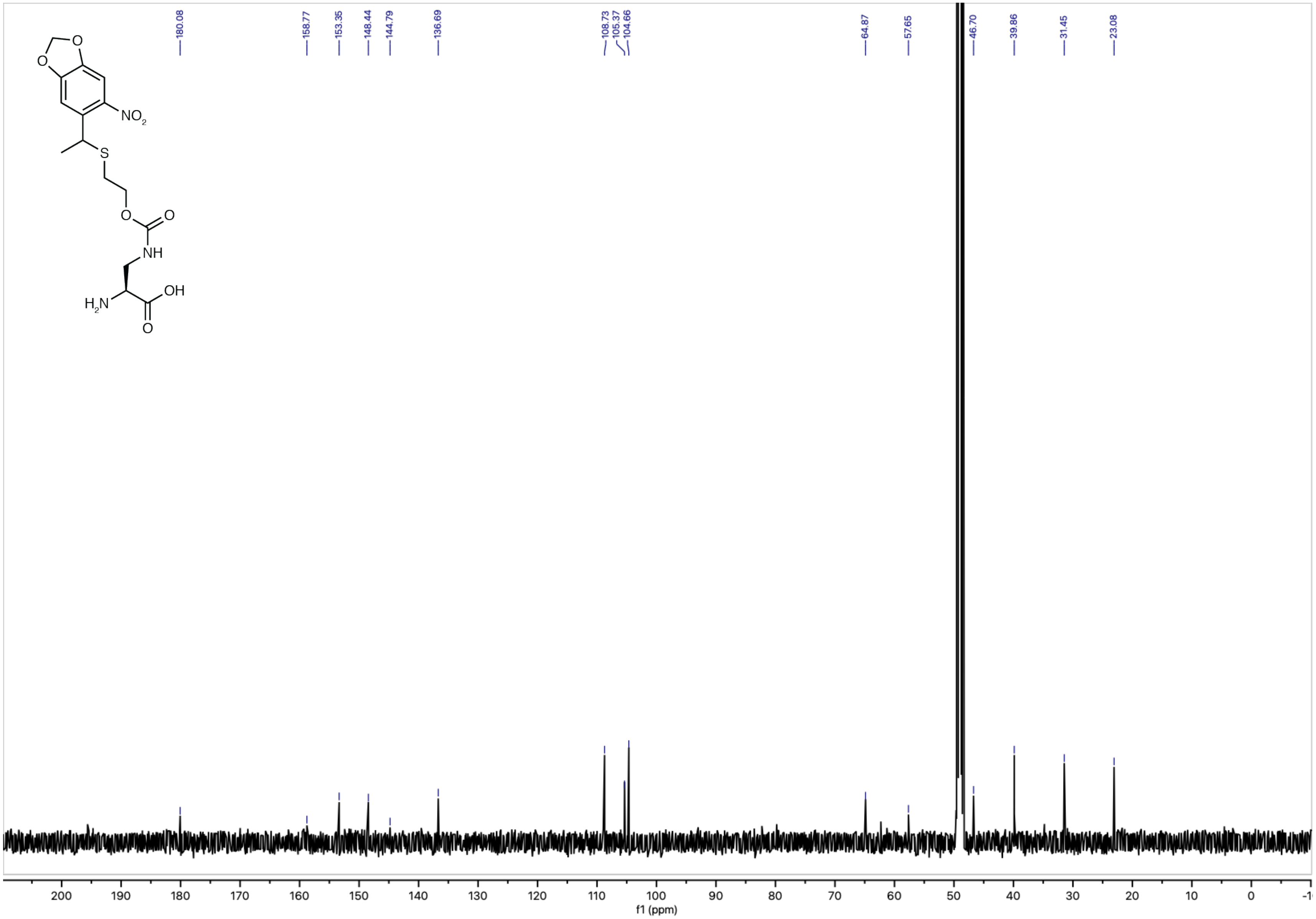

$^1$H NMR (500 MHz, methanol-$d_4$) – Standard view **(2)**

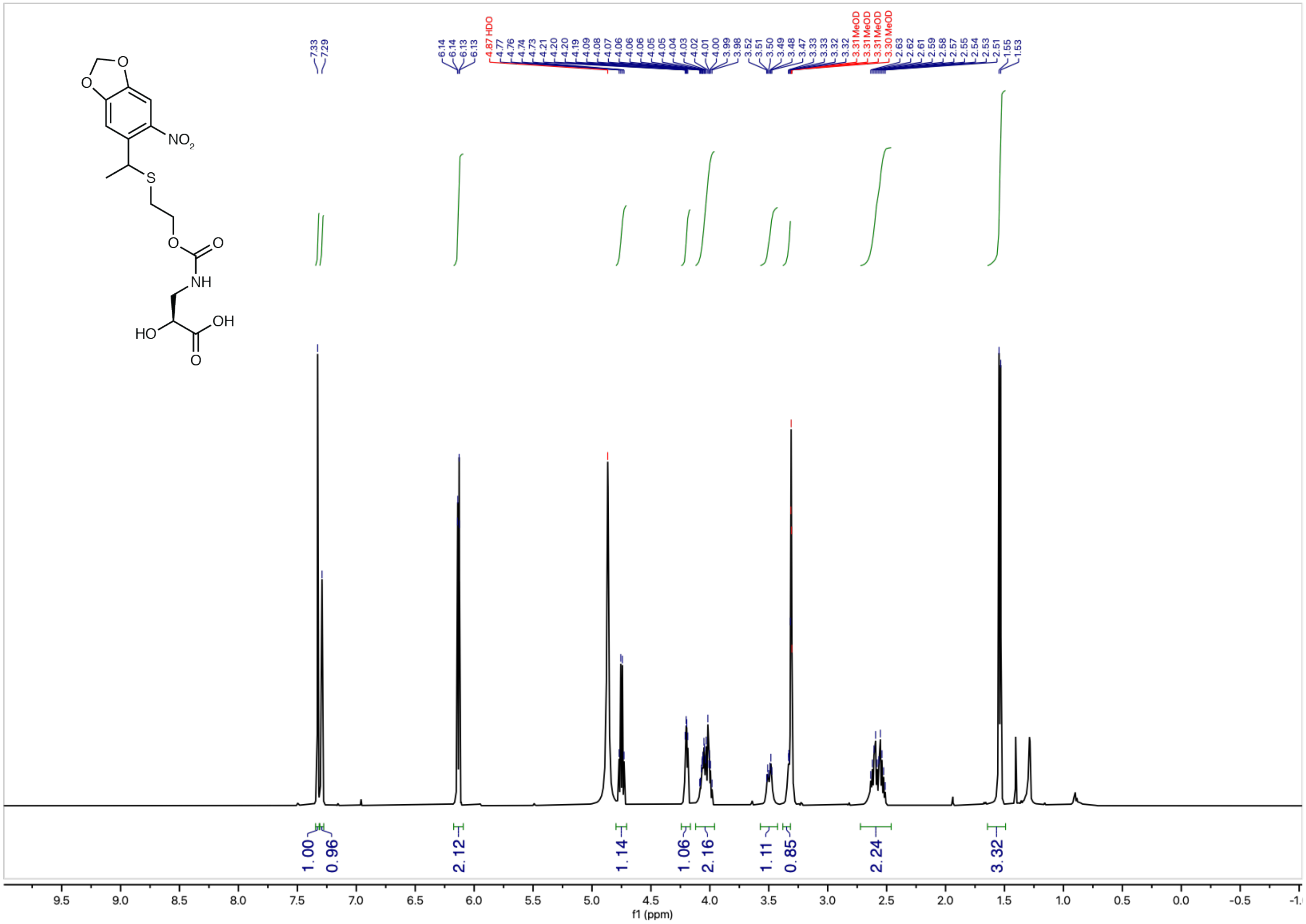

$^{13}C\{^{1}H\}$ NMR (126 MHz, methanol-$d_4$) – Standard view **(2)**

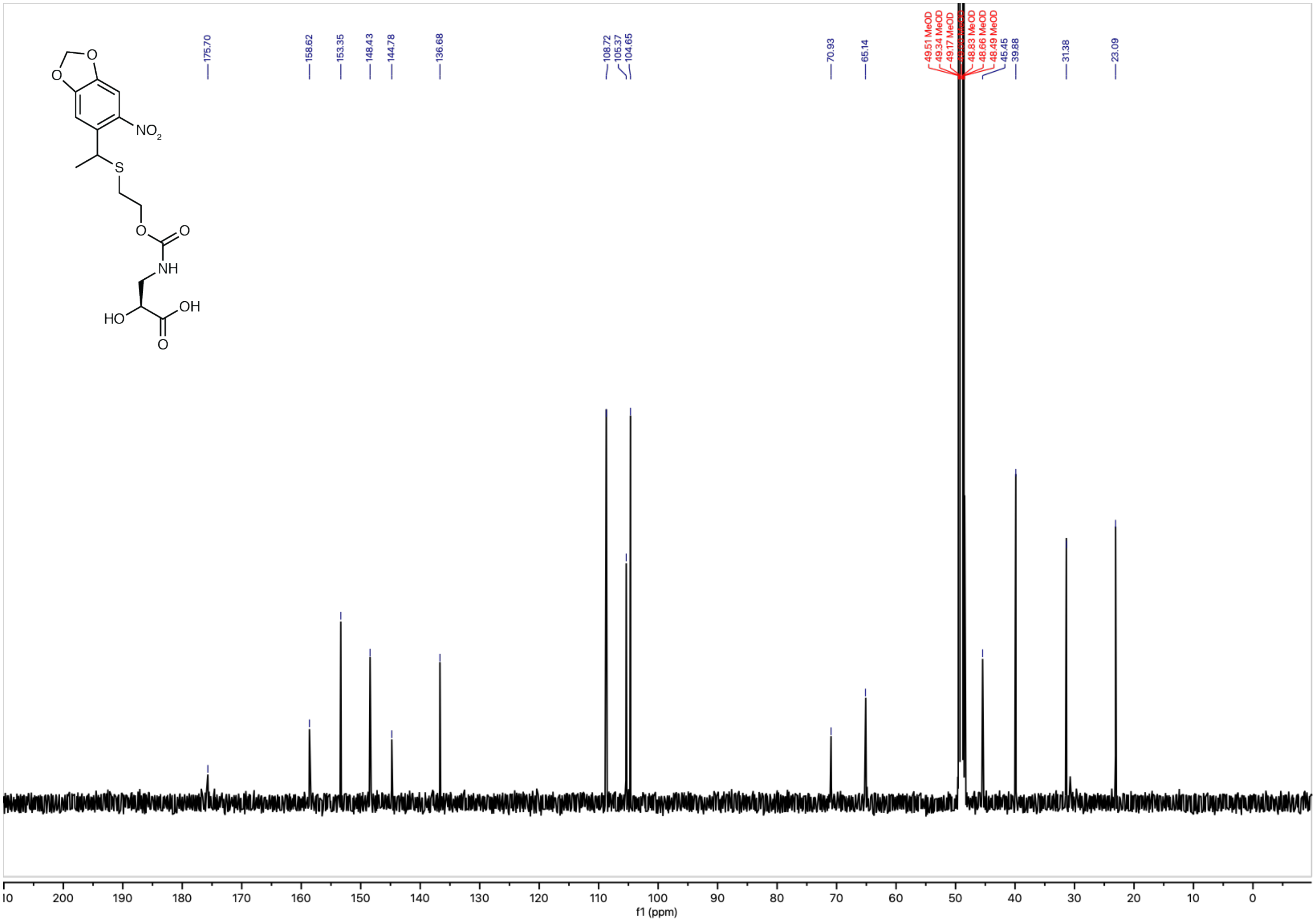

[1]H NMR (400 MHz, methanol-$d_4$) – Standard view **(3)**

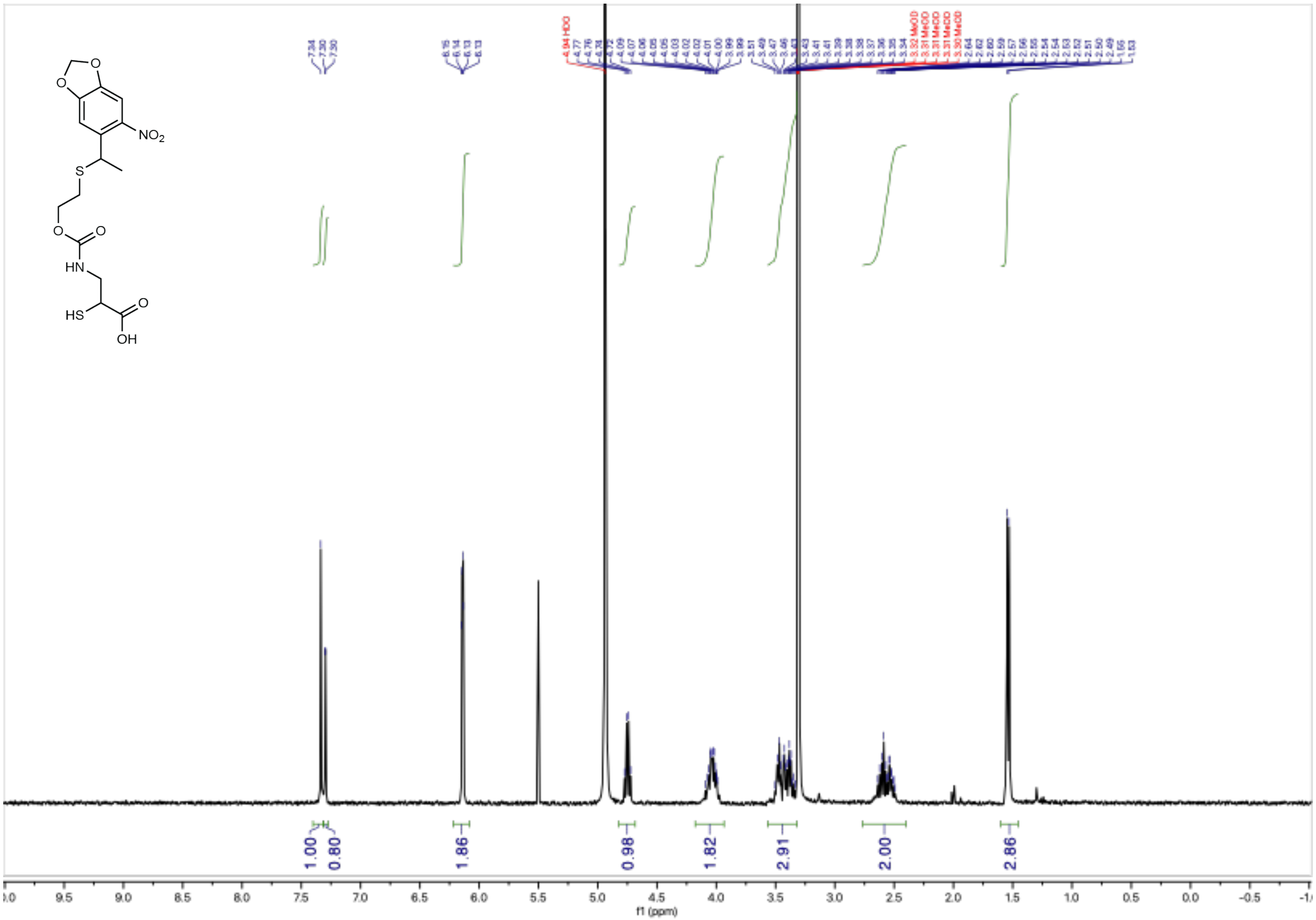

$^{13}C\{^1H\}$ NMR (126 MHz, methanol-$d_4$) – Standard view **(3)**

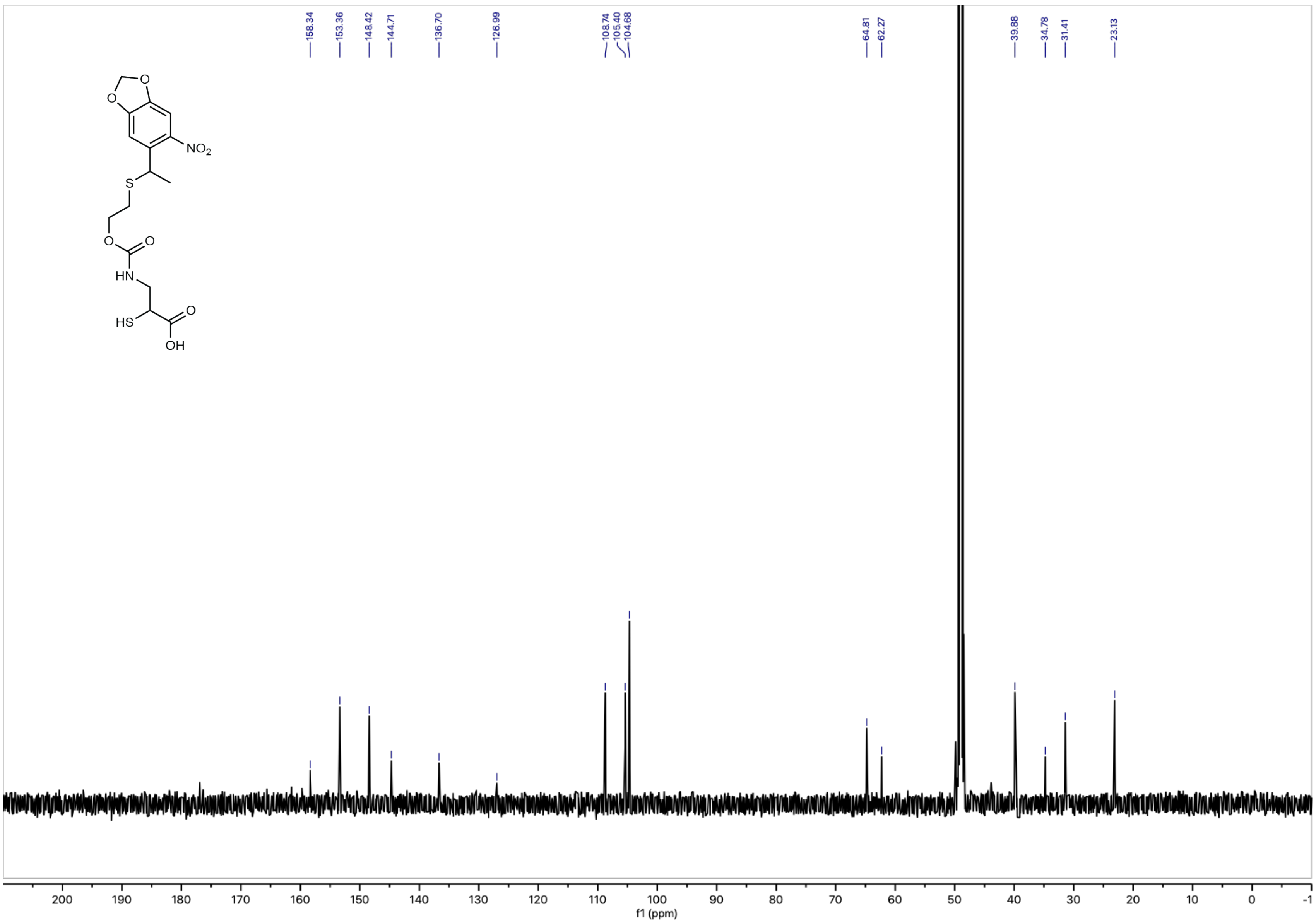

**Experimental NMR Spectra**

$^{1}H$ NMR (600 MHz, $H_2O$+$D_2O$, solvent suppression) – Standard view **(6)**

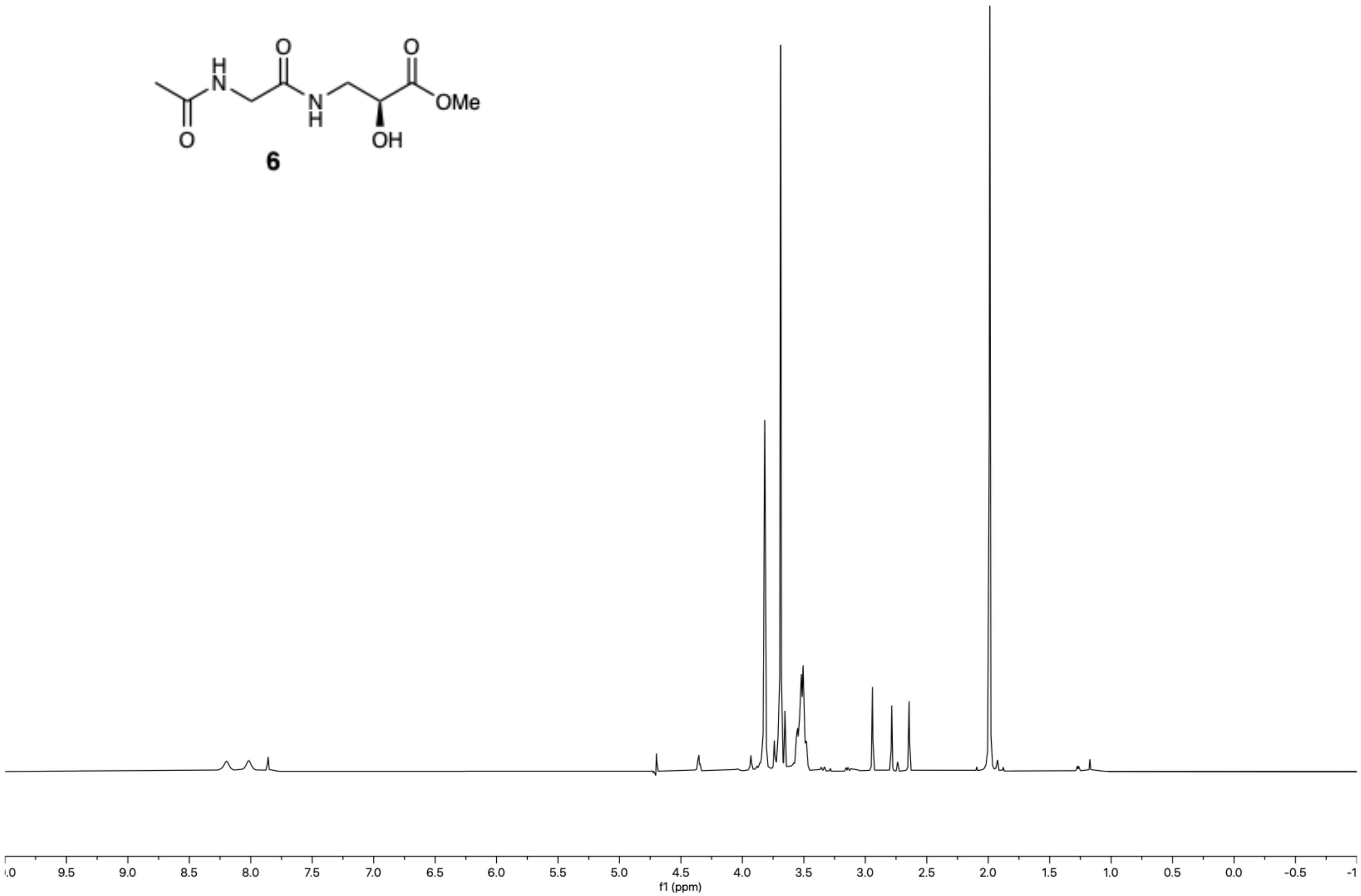

**6**

$^{1}H$-$^{13}C$ HSQC, Multiplicity edited ($H_2O$+$D_2O$) – Standard view **(6)**

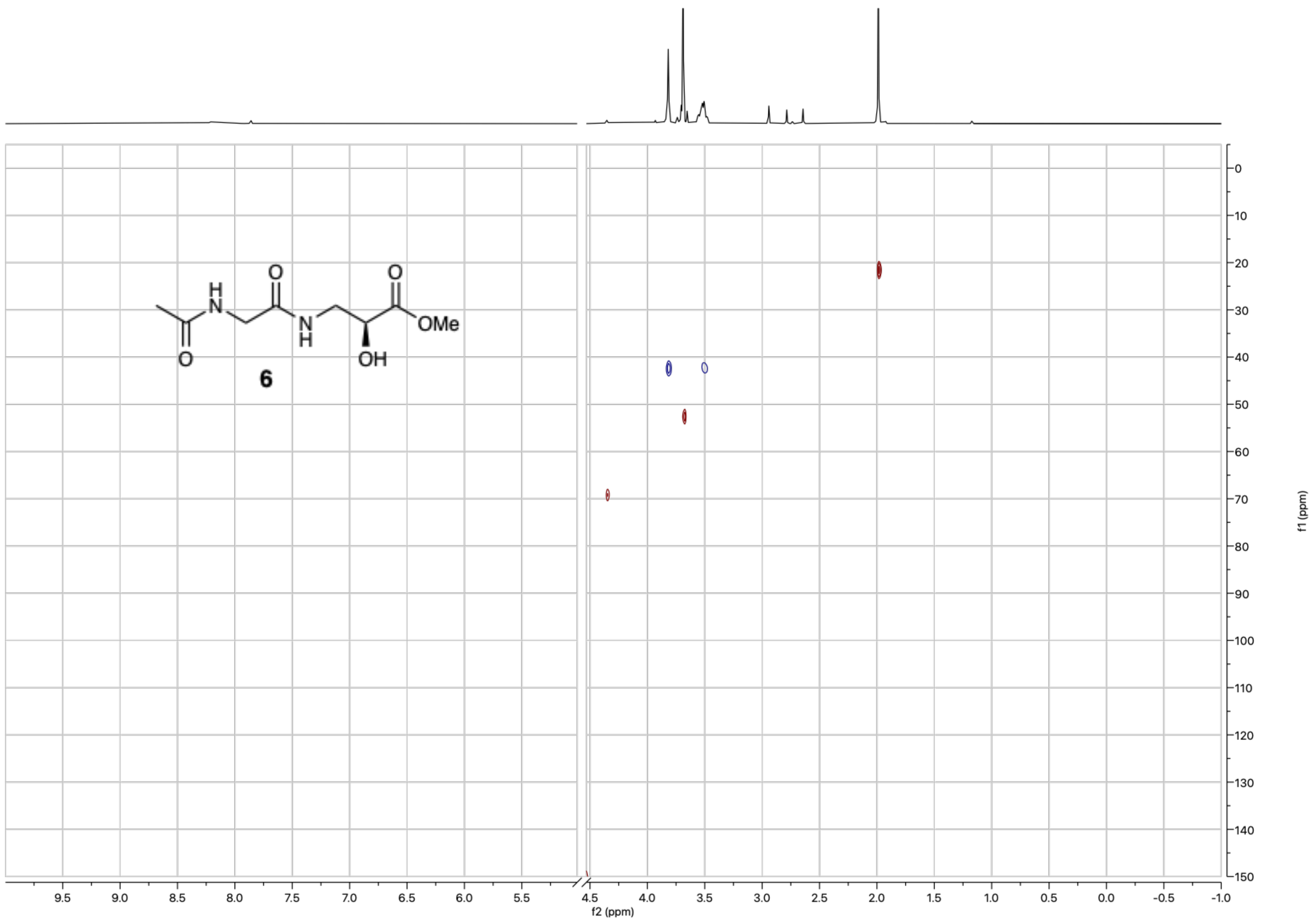

$^{1}H$ NMR (600 MHz, $H_2O$+$D_2O$, solvent suppression) – Standard view **(4)**

**4**

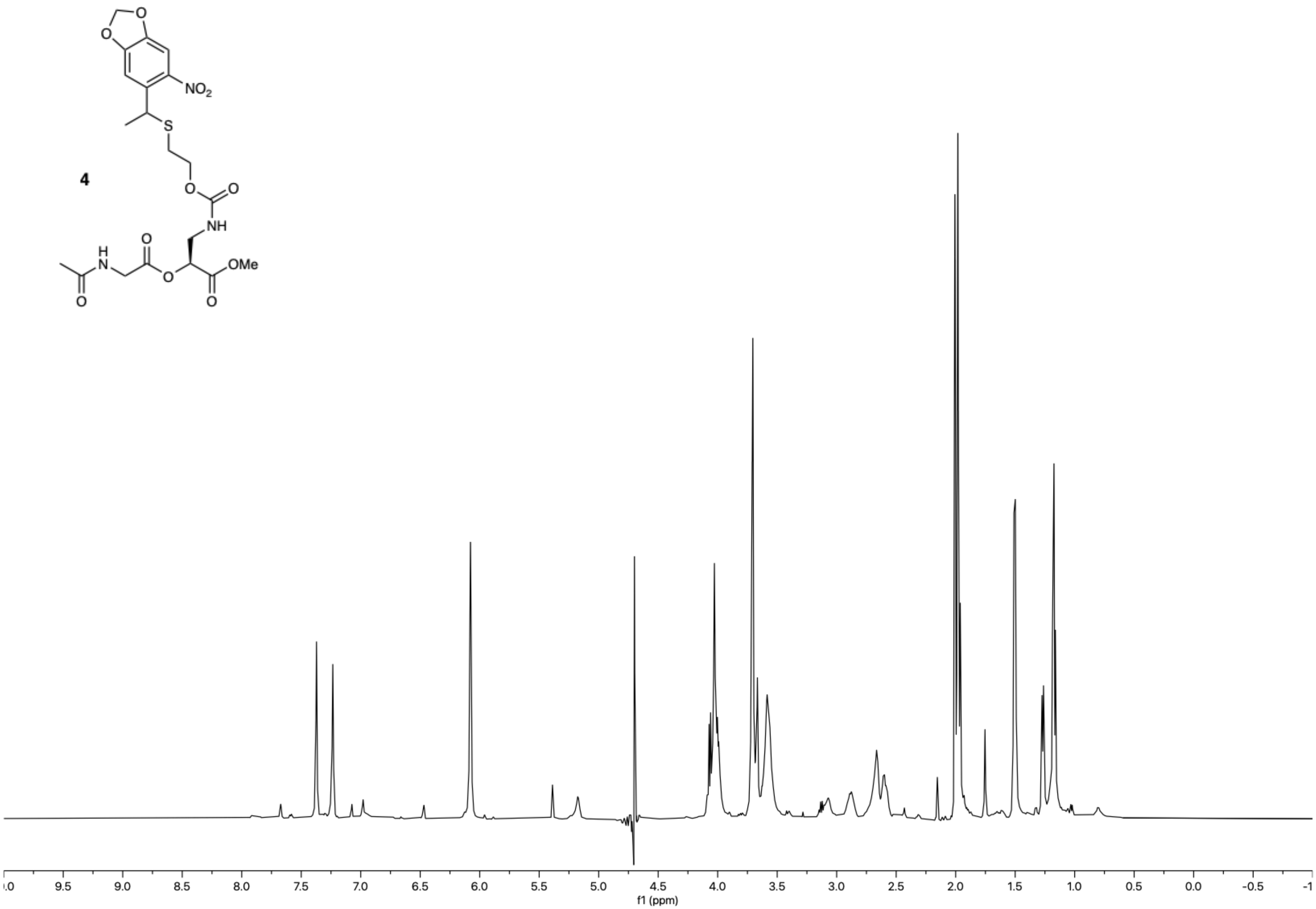

$^{1}H$-$^{13}C$ HSQC, Multiplicity edited ($H_2O+D_2O$) – Standard view **(4)**

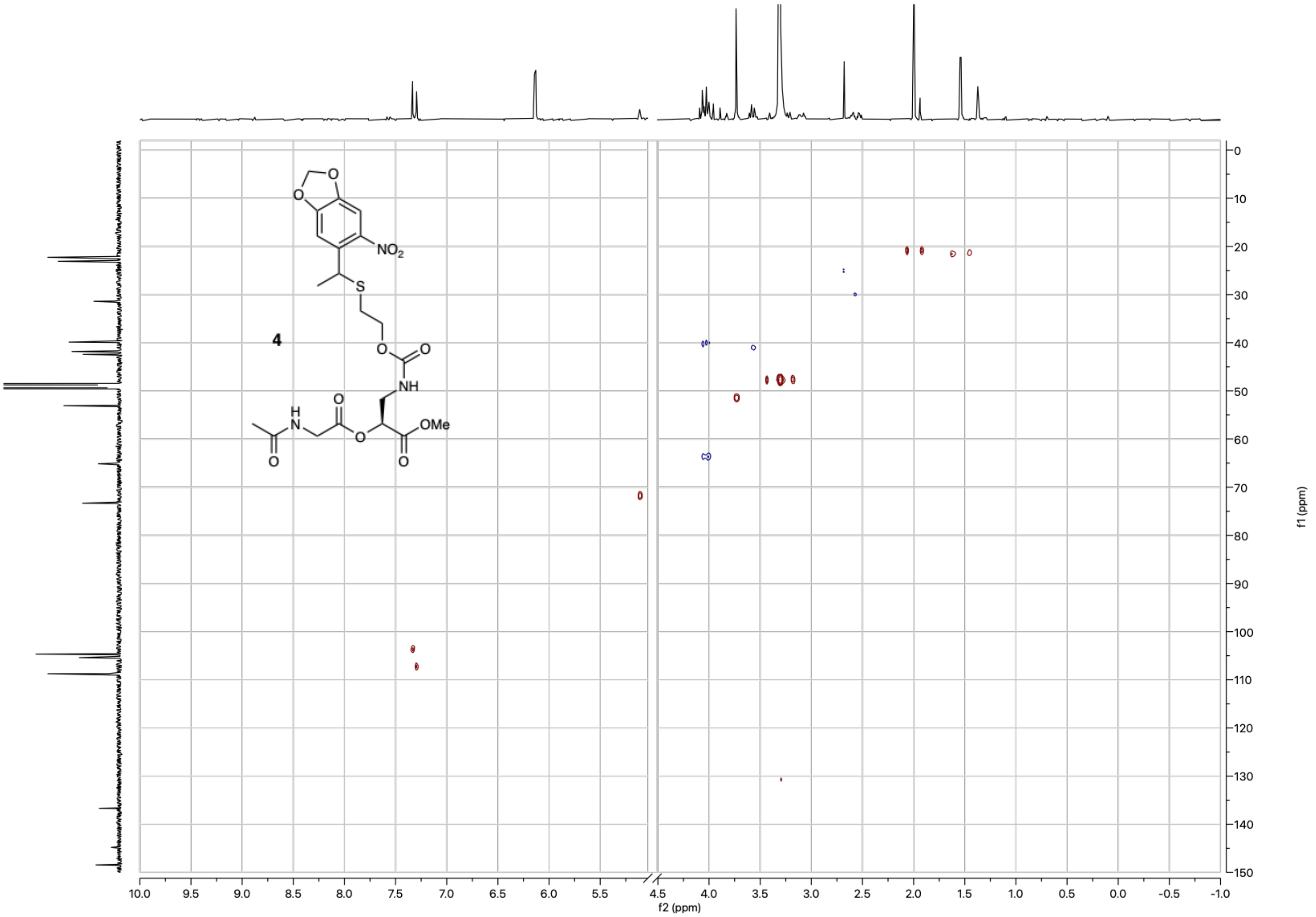

[1]H NMR (600 MHz, $H_2O$+$D_2O$, solvent suppression) – Standard view **(4),** following 2 min UV exposure

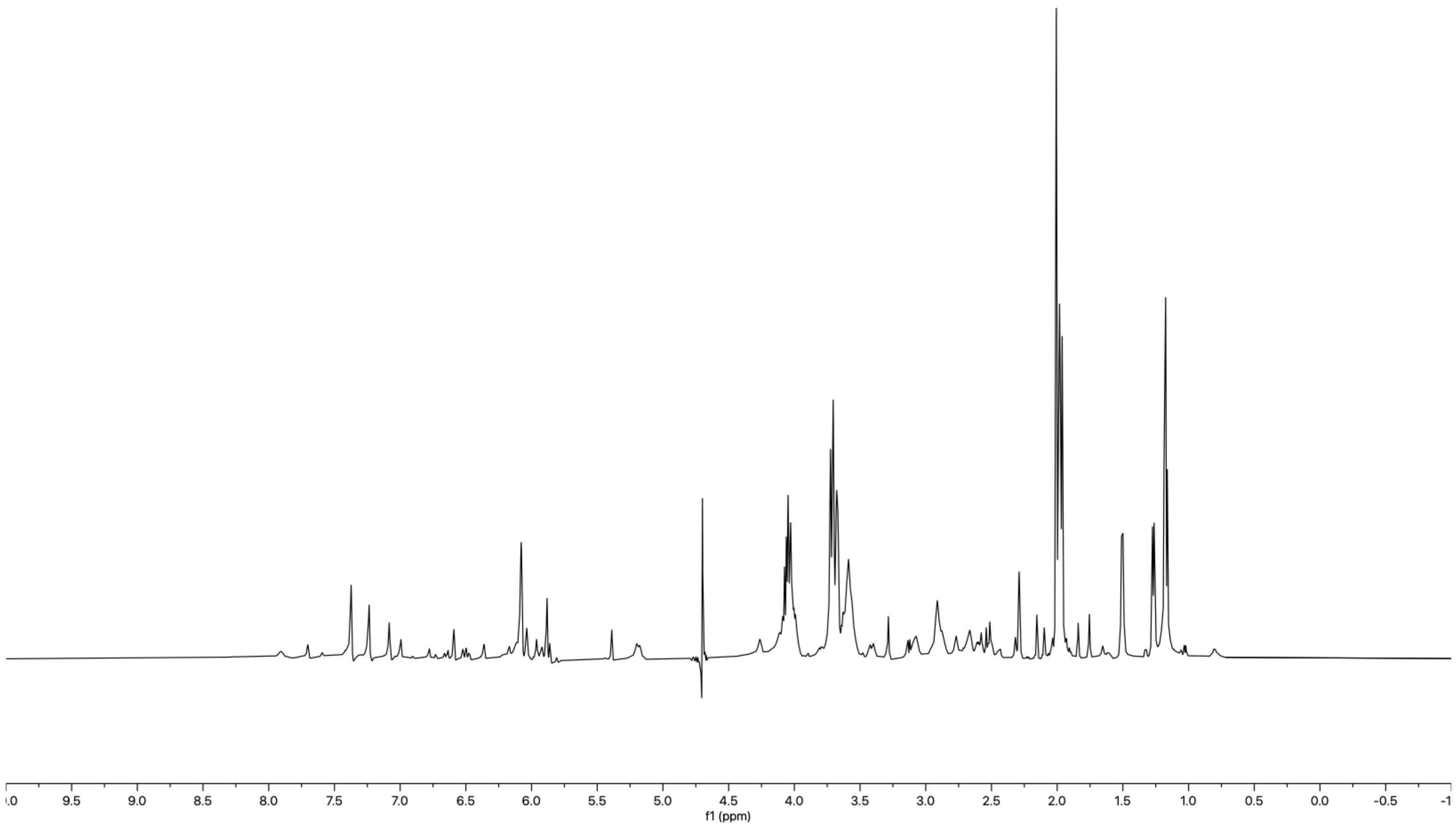

$^{1}H$ NMR (600 MHz, $H_2O+D_2O$, solvent suppression) – Standard view **(4),** following 2 min UV exposure and 4h incubation at 37 °C

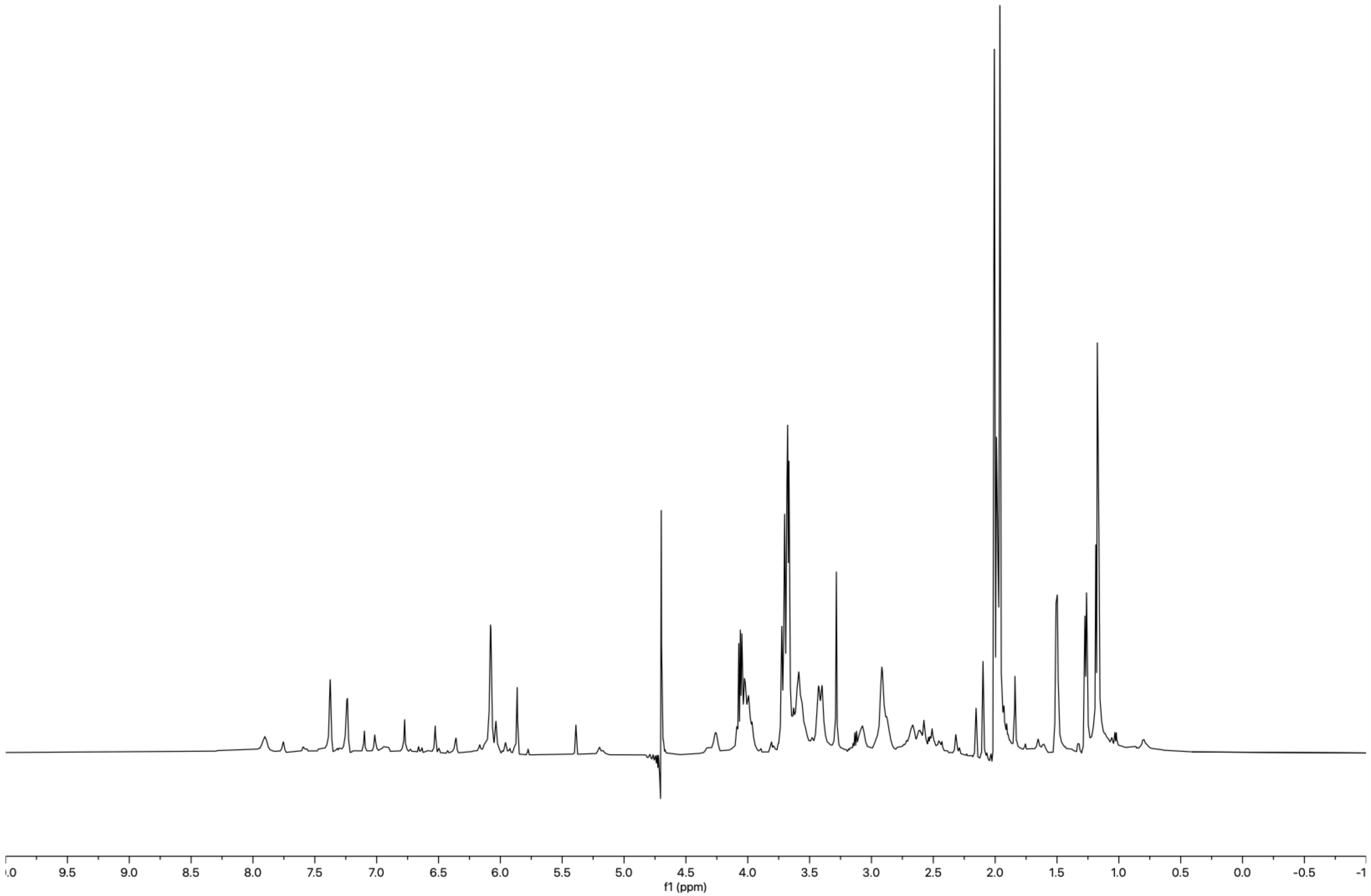

$^{1}H$ NMR (600 MHz, $H_2O$+$D_2O$, solvent suppression) – Standard view **(4),** following 2 min UV exposure and 16h incubation at 37 °C

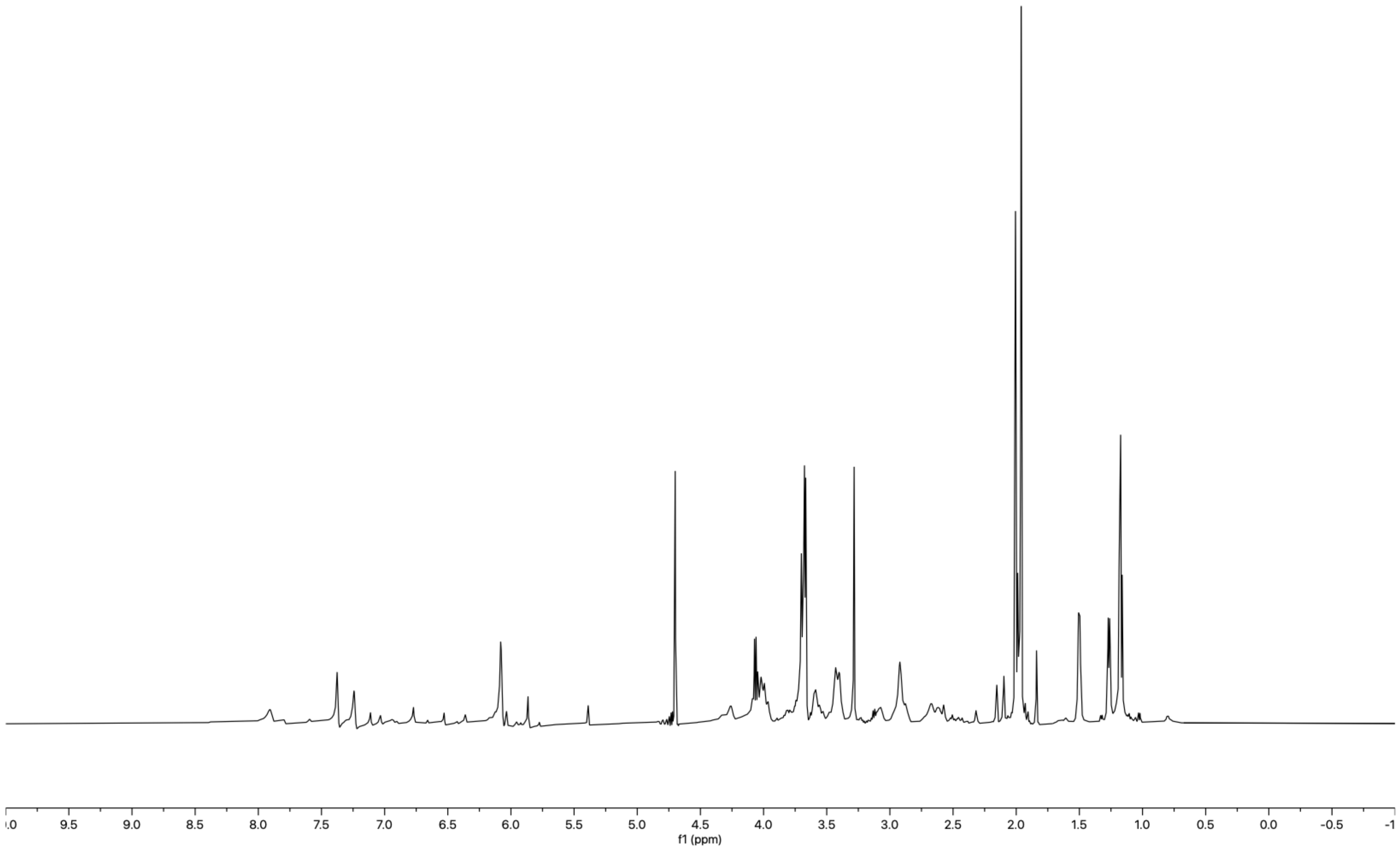

$^{1}H$-$^{13}C$ HSQC, Multiplicity edited ($H_2O$+$D_2O$) – Standard view **(4),** following 2 min UV exposure and 16h incubation at 37 °C

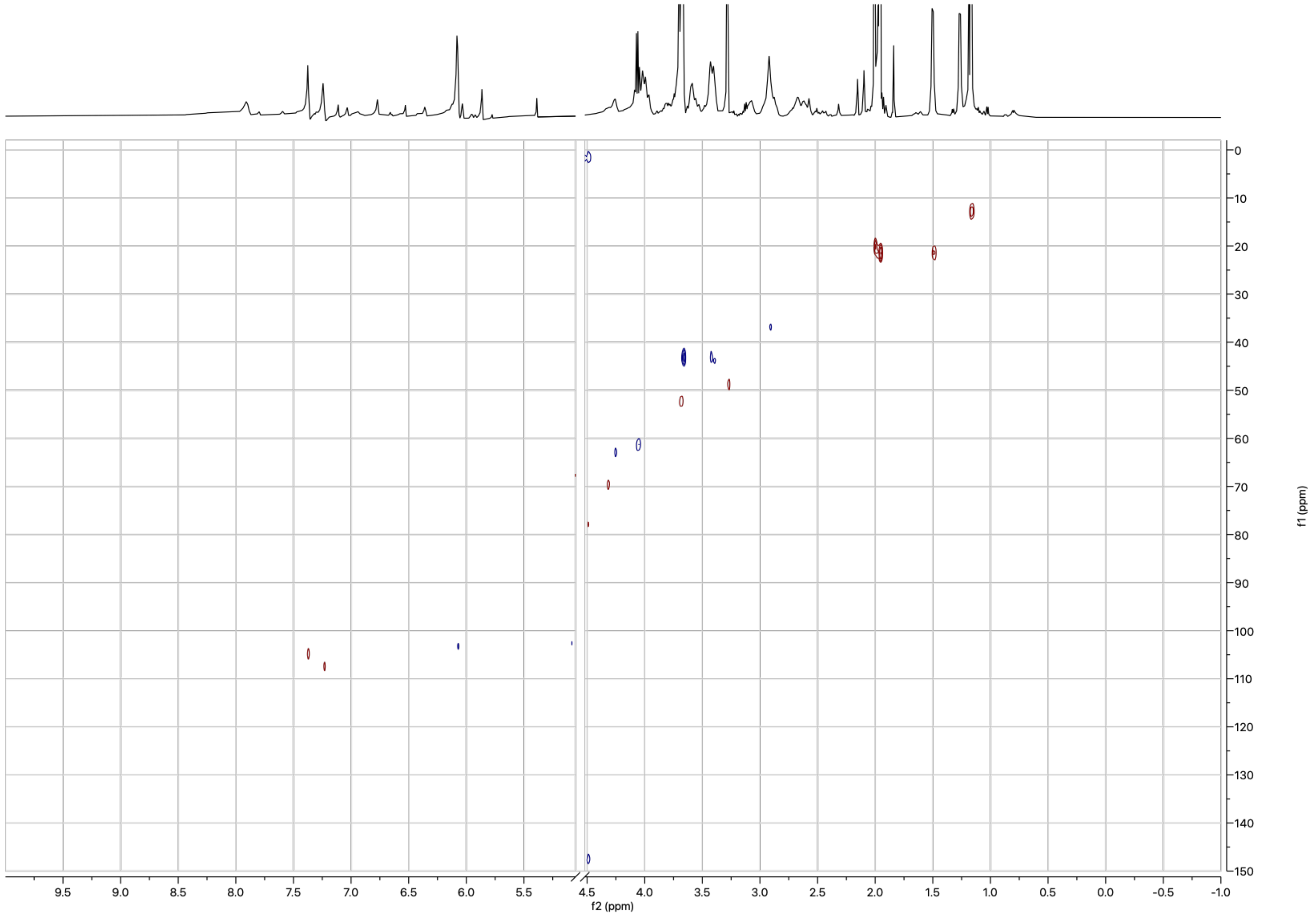
